# High-dimensional association detection in large scale genomic data

**DOI:** 10.1101/2020.11.18.388504

**Authors:** Hillary Koch, Cheryl A. Keller, Guanjue Xiang, Belinda Giardine, Feipeng Zhang, Yicheng Wang, Ross C. Hardison, Qunhua Li

## Abstract

Joint analyses of genomic datasets obtained in multiple different conditions are essential for understanding the biological mechanism that drives tissue-specificity and cell differentiation, but they still remain computationally challenging. To address this we introduce CLIMB (Composite LIkelihood eMpirical Bayes), a statistical methodology that learns patterns of condition-specificity present in genomic data. CLIMB provides a generic framework facilitating a host of analyses, such as clustering genomic features sharing similar condition-specific patterns and identifying which of these features are involved in cell fate commitment. We apply CLIMB to three sets of hematopoietic data, which examine CTCF ChIP-seq measured in 17 different cell populations, RNA-seq measured across constituent cell populations in three committed lineages, and DNase-seq in 38 cell populations. Our results show that CLIMB improves upon existing alternatives in statistical precision, while capturing interpretable and biologically relevant clusters in the data.

---

Uncovering changes across multiple biological conditions is a lasting theme in large-scale genomic data analyses across many types of studies. Examples include the analysis of tissue-specificity of gene expression patterns^1,2^, differential protein binding across cell types^3,4,5^, or causal single nucleotide polymorphisms (SNPs)^6,7,8,9^ and pleiotropic genetic variants^10^ across many genome-wide association (GWA) studies. We are specifically motivated by two contexts:

**Motivating context 1***Classification by association patterns*: if a set of subjects has been observed in many conditions, one may seek to assign subjects to classes based on the patterns of association they exhibit across biological conditions. For example, when studying plasticity of gene expression across multiple human tissues, joint analysis of these data might ask which sets of genes are collectively up-regulated together in some tissues, but down-regulated in others.

**Motivating context 2***Testing for consistent findings across many experiments*:one may desire to determine which signals are consistent across studies. For example, if one collects several ChIP-seq datasets under different experimental conditions, one may ask which loci are consistently bound in a fixed number of those conditions.

Both motivating contexts concern determining observations that have either null or significant associations across a collection of conditions. One standard approach to jointly analyzing a collection of conditions applies general clustering algorithms such as *K*-means or hierarchical clustering. Though these techniques can group signal profiles with similar association patterns together, their results do not directly provide information on condition specificity, such as which signals are consistent or differential across conditions. Somewhat similarly, time series-inspired methods such as the short time-series expression miner^11^ may be applied to genomic data collected at multiple time points. However, this approach assumes a temporal relationship across conditions and groups observations according to changes relative to a temporal baseline. This temporal assumption may not be applicable for studying genetic pleiotropy or plasticity in gene regulation, and again cannot be used to identify patterns of condition specificity. Alternatively, one may identify observations significantly associated with each condition separately, and use these individual outcomes to determine which relationships are significantly shared or differential across conditions. This technique, which is commonly used in expression quantitative trait locus (eQTL) analyses^1^, does not leverage any information-sharing among conditions, and is thus underpowered to identify shared or differential associations^12,13^. Urbut et al.^14^ improved upon single-condition analyses with a statistical model for joint eQTL analysis. This approach shows increased power; however, it makes some restrictive modeling assumptions, such as data symmetry, that are not always appropriate, especially when seeking consistent signals across conditions, as we will illustrate later. Pairwise analyses, commonly employed for differential expression analysis, also improve upon analyses of individual conditions, but still do not offer the power of a joint analysis when more than two conditions are present. Moreover, when more than two conditions are examined, it is unclear how to properly aggregate findings from a series of pairwise comparisons.

To provide interpretable joint analysis of multiple conditions, several others have introduced “association vectors” to describe an observation’s specific pattern of association across conditions; these approaches leverage mixture models to cluster observations into groups with different association vectors. For example, Andreassen et al.^10^ apply association vectors to the study of pairs of GWA studies. In this two-condition setting, they assume the presence of four association vectors {(0, 0), (0, 1), (1, 0), (1, 1)}, where a SNP described by the (0, 0) assocation vector is null in both studies, a SNP from (1, 1) is non-null in both studies, and a SNP from (0, 1) or (1, 0) is null in one of the studies, but non-null in the other. Some^15,16^ similarly use association vectors to find reproducible observations across replicated experiments, while others^17,18^ leverage them to determine which SNPs are eQTLs across various tissues.

These association vectors can be appreciated as an alternative to binarization or ternarization of genomic signals, since they assign binary or ternary *labels* to the data. A label directly reflects the pattern of condition specificity of the observations in its associated cluster. Further, as a mixture modeling approach, these labels naturally allow for heterogeneity in signals, resulting in greater model flexibility.

Yet, a remaining challenge is that models that leverage these association vectors suffer from computational intractability for even a modest number of conditions^15,17^. To understand this issue, consider *D* conditions: Let ℋ = {*H* = (*h*_[1]_, …, *h*_[*D*]_) : *h*_[*i*]_ ∈ *{*−1, 0, 1*}*} be the set of all 3^*D*^ possible configurations of association vector *H*, such that an observation described by an association vector with *h*_[*i*]_ = 1 (*h*_[*i*]_ = −1) has a positive (negative) association in condition *i*. It is clear that this model formulation becomes computationally prohibitive even for single-digit *D* because the total number of possible association vectors grows exponentially with *D*, possibly resulting in the number of model parameters exceeding the number of observations. In response to this, several restrictive assumptions are imposed. For example, Amar et al.^16^ somewhat alleviate computational burden by assuming all associations must be positive, and estimate partial latent associations for subgroups of conditions with a heuristic approach. This heuristic reduces statistical power and resolution to test for consistent findings and cannot provide a single unified clustering of observations since it is not a true joint analysis. Moreover, this approach does not distinguish an observation that is significant in opposite directions in two conditions from an observation that exhibits consistent direction of association across conditions. Alternatively, Urbut et al.^14^ make computational gains by assuming all observations come from a uni-modal distribution centered over zero, but this restriction does not always hold in practice.

We present a methodology we refer to as CLIMB (Composite LIkelihood eMpirical Bayes) that allows us to tractably estimate which latent association vectors are likely to be present in the data. Our method is motivated by the observation that the true number of latent classes, each described by a different association vector, cannot be greater than the sample size. Thus, in higher dimensions, the number of true classes is very small relative to 3^*D*^, and many candidate classes have no members. By identifying these classes through a computationally efficient pairwise composite likelihood (CL) model and rigorously filtering out unsupported latent classes, we elucidate sparsity in class membership. In doing so, the aforementioned computational intractability issue falls away, and a joint Bayesian analysis, informed by the initial CL modeling, can be performed. Using ChIP-seq, RNA-seq, and DNase-seq data collected from hematopoietic cell lineages, we demonstrate that CLIMB compares favorably against existing alternatives based on improved statistical power, precision, and model interpretability for investigating cell type-specific protein binding and chromatin accessibility, and lineage-specific gene expression patterns.

## Results

### Overview of CLIMB

We model the multi-conditional data using a constrained mixture model that encodes condition-specificity through latent association labels −1, 0, and 1 (Fig. 1a). The parameter constraints in the model enforce some general patterns commonly observed under condition-specificity: (1) observations that are associated with a condition (i.e., association label ±1) have a stronger average signal than those that are not (i.e., association label 0), and (2) observations that are associated with multiple conditions correlate with one another within a given cluster. Specifically, we assume the data are summarized as some score, and transformed to a *Z*-score, with larger values corresponding to stronger signals.

**Figure 1:**
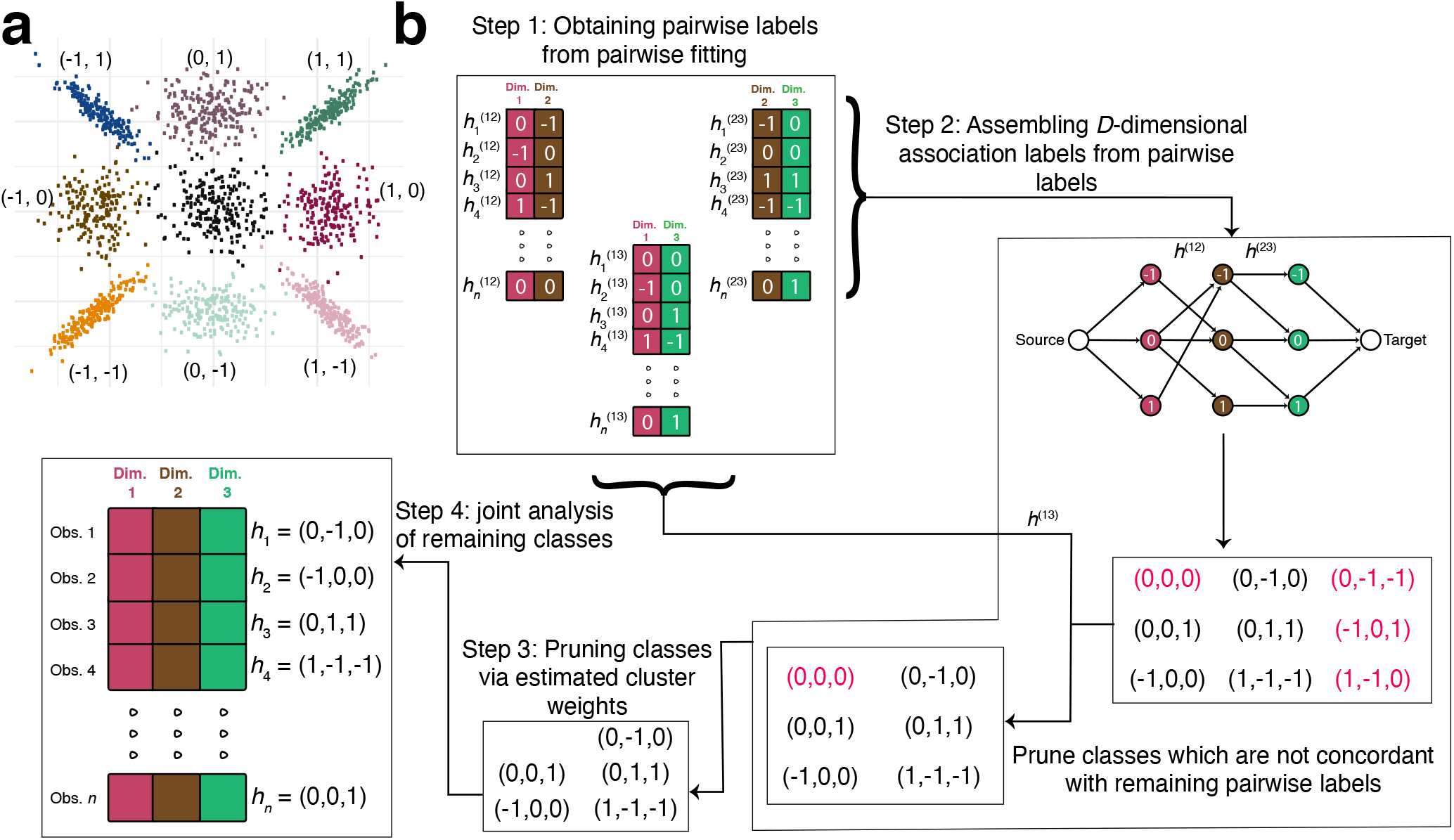
Toy examples of CLIMB. **a**, Illustration of the considered model using a simulated dataset with two dimensions. The 9 classes are annotated by their corresponding latent association vectors. The null class (0, 0) lies in the center over the origin. Classes that are non-null in at least one dimension exhibit a location shift. Only observations from classes that are non-null in both dimensions are correlated. **b**, Flowchart of CLIMB with a 3-dimensional example, with true classes whose association vectors are denoted *h*_1_, *h*_2_, *h*_3_, *h*_4_, and *h*_*n*_. Step 1 fits 3 pairwise models. Pairwise association vectors are estimated for each observation in each pairwise fit. In Step 2, we enumerate candidate 3-dimensional association vectors using a graph-based algorithm based on the estimated pairwise association vectors (shown as edges) between dimensions 1 and 2, and the estimated pairwise association vectors between dimensions 2 and 3. 9 candidate association vectors are found on the graph, but those that are colored in red are not truly present in the data. Association vectors that are not concordant with estimated association vectors from the pairwise fit between dimensions 1 and 3 are pruned. With 6 remaining candidates, one computes their prior weights (Step 3), then in Step 4 fits a Bayesian mixture model to the original, 3-dimensional data using the number of classes remaining after Step 3.

Then, letting *n* be the sample size, *D* be the dimension of the data, and *H* = (*h*_[1]_, …, *h*_[*D*]_) be a ternary latent association vector, the observed data **x** across *D* conditions follow the normal mixture model

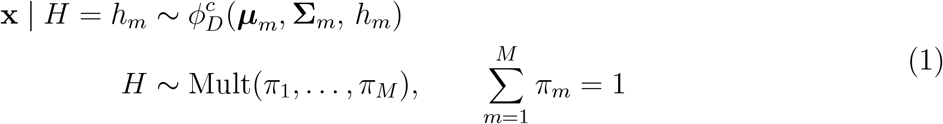

where *h*_*m*_ is the *m*^*th*^ latent class, *m* ∈ 1, …, *M*, and 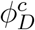 is a *D*-dimensional constrained normal distribution. The constrained normal distribution, defined presently, is used to impose association label-driven constraints:

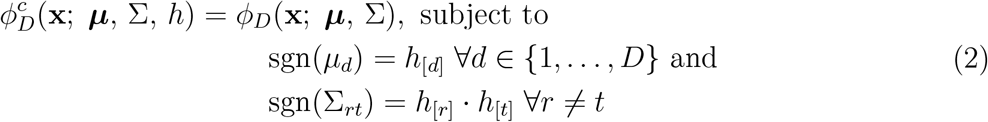

where *µ*_*d*_ is the *d*^*th*^ element of ***µ*** and ∑_*rt*_ is the (*r, t*)^*th*^ element of ∑.

Though the possible number of latent classes *M* explodes combinatorially, many latent classes likely have no members. In order to estimate the actual number of classes, we leverage information about association patterns between pairs of conditions through a pairwise composite likelihood model to eliminate classes that are unlikely to be present in the data, making the final model computationally tractable. This filtering works as depicted through a toy example in Fig. 1b, and is briefly described in four major steps:

1. *Pairwise fitting*. Fit a bi-dimensional model for each of the 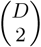 pairwise combinations of dimensions through a pairwise composite likelihood framework. The total number of possible latent classes in each bi-dimensional case is 9, and therefore tractable for typical genomic datasets. For each pair of dimensions, we estimate which subset of the 9 possible configurations of the latent association vector are supported by the data across those 2 dimensions by utilizing a penalized mixture model^19^. This mixture model penalizes the class mixing weights, such that classes that are likely without members are removed from the pairwise model. Unlike many composite likelihood approaches that assume independence across dimensions^15,20^, the pairwise model takes account of dependence between each pair of conditions.
2. *Assembling D-dimensional association labels from pairwise labels*. Use the estimated pairwise association vectors to assemble a preliminary list of feasible *D*-dimensional association vectors. *D*-dimensional association vectors that are inconsistent with inferred pairwise labels will be deemed infeasible and pruned.
3. *Pruning association labels with insufficient cluster weights*. Estimate the mixing weights for the remaining latent classes using the estimates obtained from the pairwise fits, pruning classes with insufficient weight and ensuring that *M* ≤ *n*.
4. *Empirical Bayesian estimation of the full D-dimensional model*. Reestimate parameters for the *D*-dimensional mixture model based on the final list of classes using a Bayesian approach. Inform prior hyperparameters with parameter estimates obtained from the pairwise fits. This final step ensures information across all dimensions is considered.

CLIMB’s model output is useful for a host of analyses, including: (1) using association labels and class membership to elucidate condition-specificity, (2) using class membership probabilities to test for consistency in signals across conditions, (3) using estimated cluster covariances to infer similarity between conditions, and (4) using estimated cluster means to obtain a parsimonious characterization of dominant patterns of condition-specificy. See *Methods* and supplement for details on these downstream analyses.

### Simulations

We used simulations to compare CLIMB to the available methods for multiconditional analysis, Urbut et al.’s mash^14^ and Amar et al.’s SCREEN^16^. We selected these two methods to compare against because they are also designed to analyze many conditions for obtaining information on condition specificity. In a separate simulation, we also compare CLIMB to DESeq2^21^, a widely used tool for pairwise differential expression analysis. Although DESeq2 focuses on pairwise comparisons, its wide adoption makes it a worthy comparison in the context of RNA-seq analysis.

We consider three data types commonly encountered in genomic analyses: ChIP-seq data, differential analysis output from RNA-seq data collected from treatment/control tissue pairs, and RNA-seq data. The first simulation aims to study cell type-specificity of patterns of protein binding across different cell types (motivating context 1), the second aims to identify which genes are dysregulated in a consistent manner across different diseased tissues when compared against normal tissues, and the final simulation aims to identify genes whose expression levels change across cell differentiation (motivating context 2). These datasets exhibit different distributional structures. For example, signals in simulation 1 have a positive sign (Supplementary Fig. S1a), but signals in simulations 2 and 3 can be positive or negative. The strictly positive nature of signals in simulation 1 arises from the fact that identified protein binding sites from ChIP-seq data are output from a peak-calling routine, where each signal indicates evidence for the presence of a ChIP-seq peak at a given genomic location. In contrast, the data in simulation 2 are derived from *P*-values that indicate whether genes are relatively over- or under-expressed in a diseased tissue relative to a normal counterpart tissue. This translates to *Z*-scores exhibiting both positive and negative signals, and data that are more symmetrically distributed about the origin (e.g., see Supplementary Fig. S1b). A unifying goal of all simulations is to evaluate the capacity of all methods to adapt to data types with different distributions. See *Testing consistency of effects* for description of statistical test used; see *Simulations and comparisons* and supplement for further details on the simulation procedure. A computational cost analysis is also conducted (Supplementary Fig. S2). CLIMB uniformly performed better than SCREEN and mash in simulations 1 and 2 across several quantitative metrics (Supplementary Fig. S3–S9), including sensitivity and precision. CLIMB, mash, and SCREEN respectively had average F1-scores of 0.97, 0.77, and 0.74 for simulation 1, and 0.46, 0.45, and 0.12, for simulation 2, at an *α*-level of 0.05. CLIMB also outperformed DESeq2 in simulation 3, for identifying differentially expressed genes in a multi-condition setting (Supplementary Fig. S5). For this simulation, CLIMB and DESeq2 had F1-scores of 0.65 and 0.48, respectively, at a confidence threshold of 0.05. If effects are not shared in more than 2 conditions, as they were in our simulations, then CLIMB gains no power over DESeq2 or other pairwise methods. These results indicate that CLIMB is well-suited for identifying patterns of association in the data as well as consistent and differential signals.

### Case studies

We showcase CLIMB’s utility by analyzing multiple datasets collected as part of the VISION (ValIdated Systematic IntegratiON of hematopoietic epigenomes)^22,23,24^ and ENCODE^25^ projects. These VISION and ENCODE data were collected from, respectively, 17 murine and 38 human hematopoietic cell populations across differentiation. The primary goal of the VISION project is to understand the interplay between transcriptomic variation and mechanisms of gene regulation during hematopoiesis, while the ENCODE project aims to describe functional elements in the human genome more broadly.

First, we study VISION CTCF ChIP-seq data in 17 hematopoietic cell populations^26^. While CTCF binding sites that are invariant across cell types are known to maintain chromatin structures^27^, the function of more cell type-specific CTCF binding sites remains largely unknown^5,28,29^. We show how CLIMB can be used to aid in tackling this question. Next, we examine VISION RNA-seq data collected from a subset of these cell populations to probe the transcriptomic changes that commit multipotent cells to different fates. Results from these analyses demonstrate CLIMB’s ability to elucidate interrelationships between cell populations in different genomic data types, produce interpretable classes, and conduct lineage-specific differential analyses. Finally, with ENCODE’s DNase-seq data, we illustrate CLIMB’s ability to identify novel classes of tissue-specific regulatory elements.

### VISION CTCF ChIP-seq

We applied CLIMB to CTCF ChIP-seq of chromosome 11 from 17 murine cell populations. This analysis yielded a final model that included 15 non-empty classes. Among these, 2 classes described constitutive binding behavior, while the remaining were more cell type-specific (see Supplementary Fig. S10 for an illustration of all classes). Similar results are obtained for chromosome 7 (see Supplementary Section *Analysis of CTCF ChIP-seq on chromosome 7*).

#### Constitutively bound CTCF is the dominant class

Previous work has noted that CTCF binding is largely consistent across cell types^5,27,30^. We identified two such classes of conserved loci from CLIMB’s model fit. The first is the class of all ones, corresponding to the collection of loci bound by CTCF across all cell types. The second is the class of all ones except for the CFUE population, corresponding to the collection of loci bound by CTCF in all but the CFUE cell population, likely reflecting lower signal-to-noise ratio in the CFUE dataset. Indeed, the CFUE experiment had the lowest quality as measured by Fraction of Reads in Peaks (FRiP) score^31^ (0.031, compared against next lowest iMK with FRiP score 0.054 and CMP with FRiP score 0.097). In agreement with previous studies, these two classes make up ~36% of all loci in the analysis. Moreover, consistent with others^30,32^, the average signal strength (based on the estimated class means) for bound loci within the two constitutive classes is significantly larger than the average signal strength for bound loci that are not widely shared across cell populations (one-sided *t*-test, *P* = 5.02 × 10^−12^).

#### Differential CTCF binding is predictive of cell population relationships

Although CTCF binding is largely consistent across cell types, previous studies suggested that changes in its binding patterns modify gene expression programs, affecting developmental cues or cell function^5,32,33^. We asked whether the classes discovered by CLIMB support the idea that changes in CTCF binding relate to hematopoietic development. To address this question, we clustered the cell populations based on the estimated class covariance matrices^34^ (see supplementary *Implementation details*). CLIMB’s clustering, shown in Fig. 2b, closely reflects the expected lineage relationship in Fig. 2a. This result supports the claim that changes in CTCF binding occur in a lineage-specific manner, and that CLIMB is well-suited to tease out this information from the data. In contrast, the clusterings based on mash and the standard hierarchical clustering using Pearson correlation depart further from the expected lineage relationship (Baker’s Gamma^35^ correlation coefficients, which measures the similarity between two hierarchical tree structures, of 0.251, 0.096, and 0.209 for CLIMB, mash, and Pearson, respectively, when compared against the ground truth tree in Supplementary Fig. S11). This suggests that mash does not sufficiently capture CTCF binding patterns across cell types, and that simple correlation measures cannot effectively distinguish between different classes of signals in the data. The low signal in the CFUE experiment likely caused the hierarchical clusterings by both CLIMB and Pearson correlation to isolate the CFUE cell from the remaining cell populations on the hierarchical tree. CLIMB exhibits robustness to this challenge, identifying this cell as an outlier among all experiments, while still achieving a hierarchical clustering that reflects the expected relationship among the remaining cell populations.

**Figure 2:**
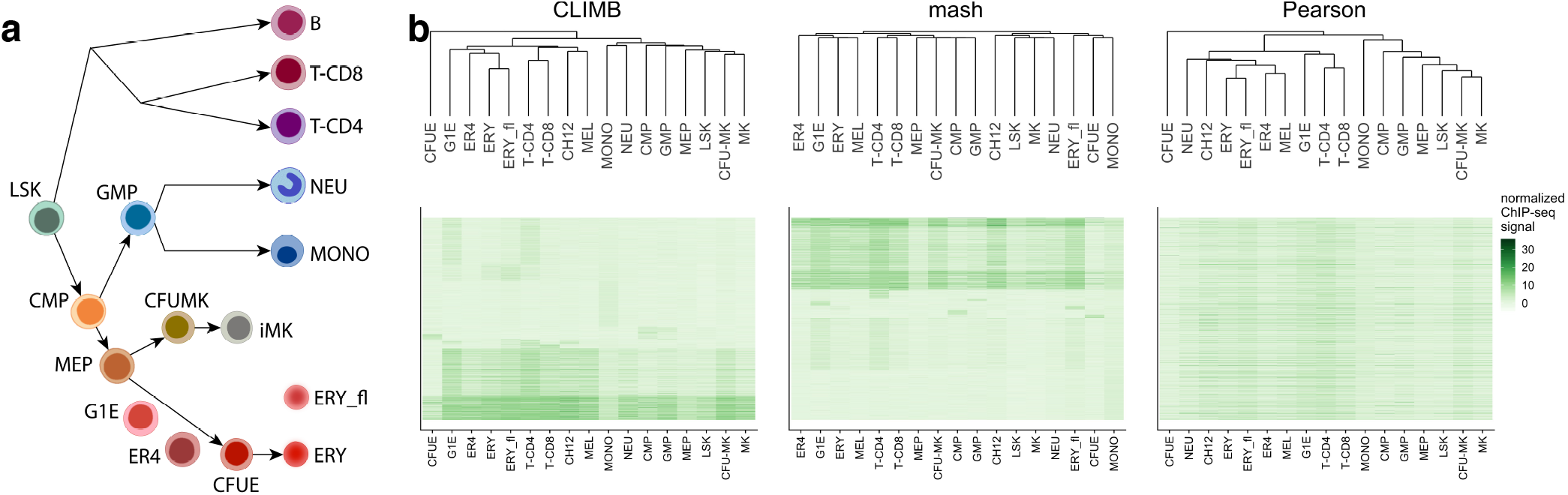
CLIMB uncovers interrelationships among hematopoietic cell populations based on CTCF binding patterns. **a**, Expected relationship among cell populations. **b**, Heatmaps displaying bi-clusterings of all ChIP-seq data for chromosome 11 based on CLIMB, mash, and Pearson correlation. The columns, corresponding to different cell populations, are ordered according to the dendrogram for each clustering method. The rows, corresponding to each loci, are ordered based on class membership (for CLIMB and mash) and Pearson correlation (for Pearson), respectively. (CH12 and MEL are murine lymphoma and erythroleukemia cell lines, respectively, and thus do not clearly occupy one space in the lineage, though CH12 is most related to B cells, and MEL is a mature erythroid cell type.)

#### CLIMB identifies succinct groupings of CTCF binding patterns

Visualization of binding sites assigned to different classes is important for identifying biologically meaningful patterns. To facilitate visual examination, CLIMB provides a means to merge similar classes based on model output (see supplementary *Implementation details, Obtaining parsimonious characterization* for details on the class merging procedure). From the VISION CTCF dataset, CLIMB clusters the binding sites into 15 non-empty classes. To simplify the visualization, we aggregated these classes into 5 parent groups, with sizes ranging from 254 to 5,462 binding sites. Supplementary Fig. S12a displays the average signal strength (Equation 30) associated with each of these groups. For example, group 1 includes constitutive binding sites, while group 4 contains progenitor-specific binding sites, and group 5 contains binding sites constituent to mature erythroid and T cells. Supplementary Fig. S12b displays the locations of the binding groups within the genomic region around murine gene *Bcl11a*, whose gene product is involved in gene regulation of multiple cell types.

#### CTCF binding patterns relate to epigenetic states during differentiation

We next examined how CLIMB’s classes of CTCF binding patterns relate to chromatin accessibility and various histone modifications. Interestingly, though we only supplied CTCF ChIP-seq data to each method, the classes estimated by CLIMB also displayed cell type-specific behavior of chromatin accessibility as measured using ATAC-seq and epigenetic histone modifications H3K4me1 and H3K4me3 (Fig. 3a– b). Further, using GREAT^36^ (Genomic Regions Enrichment of Annotations Tool), we identified that classes that exhibit erythroid- and immune cell-specific binding patterns are indeed enriched in erythroid- and T cell-specific functions (Fig. 3c). In contrast, the classes identified by mash do not appear to relate to epigenetic modifications (Supplementary Fig. S13–S16). In fact, there is not a large amount of overlap between CLIMB’s and mash’s estimated classes (Supplementary Fig. S17), altogether suggesting that CLIMB effectively captures biologically meaningful protein binding patterns.

**Figure 3:**
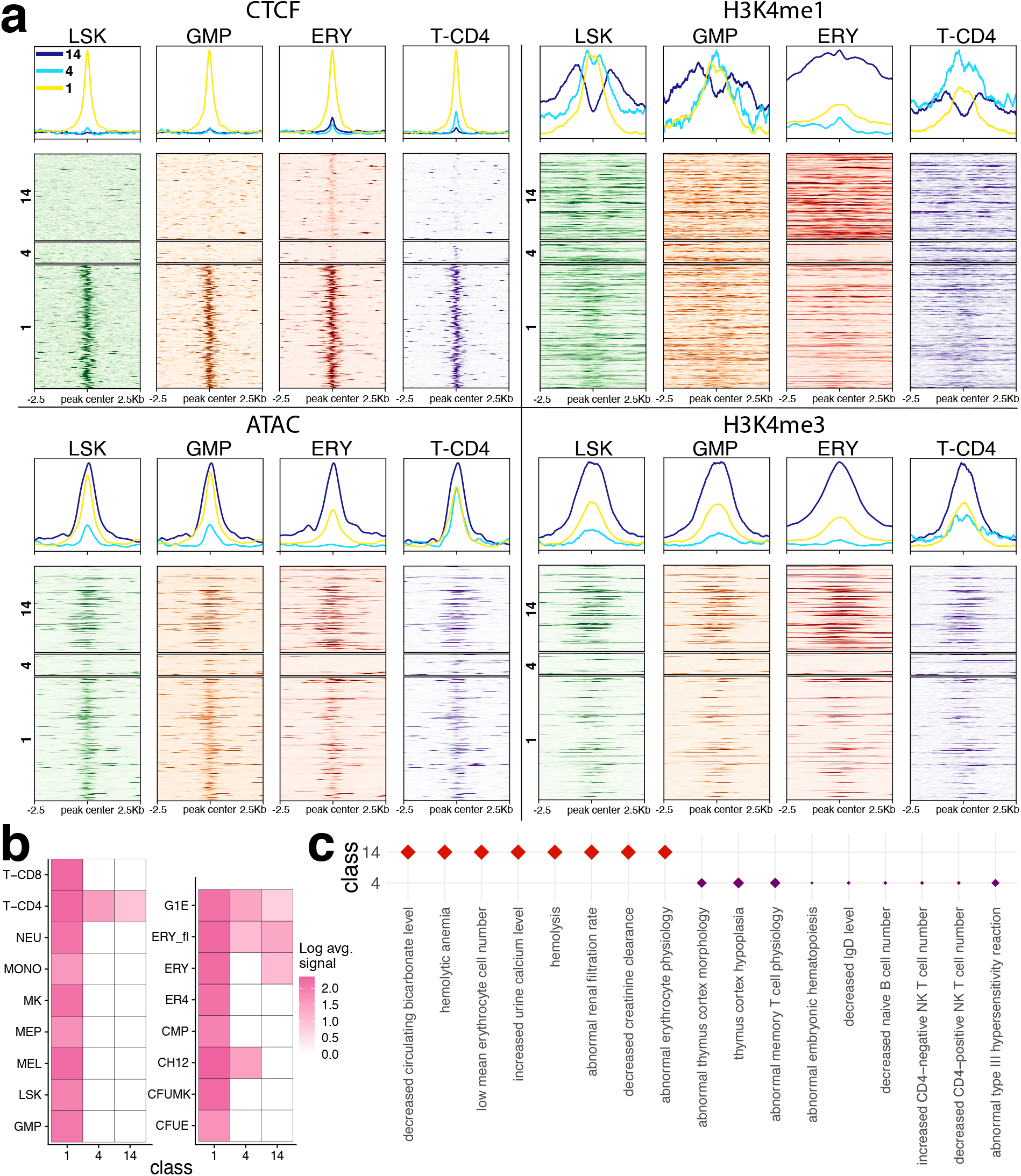
CTCF binding patterns uncovered by CLIMB capture different patterns of epigenetic modifications. **a**, Data from the loci on chromosome 11 that belong to classes of CTCF binding patterns (numbered 1, 4, and 14) identified by CLIMB are shown. The original CTCF ChIP-seq, alongside ATAC-seq and histone modification ChIP-seq data in 4 hematopoietic cell populations reveal differing patterns of epigenetic modifications across cell populations. **b**, Log class means based on CLIMB’s model of CTCF binding patterns for the 3 classes in **a. c**, Significantly enriched mouse phenotypes (FDR *<* 0.05 for all) associated with the plotted classes. Class 1, containing loci with CTCF bound in every cell type, is not significantly enriched in any mouse phenotypes. Class 4 is enriched with terms related to T and B cells and the thymus, while class 14 contains terms related to red blood cells and kidney function.

The classes learned by CLIMB also provide hypothesis-generating discoveries. For instance, though class 14 exhibits consistent but low signal for CTCF binding only in erythroid cells, these same sites are in open chromatin in all four cell populations, as assayed by ATAC-seq. Since transcription factor binding is often regulated by differentially open chromatin, this raises a question of what is driving the erythroid-specificity of this class. One possibility is that the sites could be bound by other transcription factors, occluding CTCF. The pattern of H3K4me1 as high surrounding peaks of H3K4me3 in these class 14 sites suggests that they may be promoters. Indeed, ~6% of the CTCF-bound sites in class 14 (as well as the constitutively bound classes 1 and 2) overlap with transcription start sites from GENCODE.v35, while this occurred on average ~2% for the remaining classes, which fits with the patterns of histone modifications and ATAC-seq data. This hypothesis is testable in further studies.

### VISION RNA-seq

We next used CLIMB to perform lineage-specific differential expression analysis. In the hematopoietic cell system, LSK, CMP and MEP are multipotent cells that differentiate into different terminal cells, such as ERY, MONO, NEU, and iMK cells (Fig. 2a). We considered three paths: the erythroid lineage (LSK → CMP → MEP → CFUE → ERY), the megakaryocytic lineage (LSK → CMP → MEP → CFUMK → iMK), and the myeloid lineage (LSK → CMP → GMP → MONO/NEU). The differentially expressed genes identified in each linage are expected to be related to the biological function of the specific differentiation path and cell fate commitment. The datasets for these lineages respectively contained 21,303, 20,995, and 22,940 expressed genes.

#### CLIMB identifies lineage-specific genes related to cell development and differentiation

We sought to identify genes that show varying gene expression levels across each differentiation path. We first fit a model with CLIMB to each lineage. We then pinpointed the genes that exhibit differential signals across each lineage based on model fit. To proceed, we first identified genes with consistent signals by performing a statistical test (see *Methods*). Briefly, a gene was considered “consistently expressed” across the lineage if its probability of belonging to a class that is interpreted as describing consistent expression behavior is sufficiently large. These classes are: (−1, −1, −1, −1, −1), (0, 0, 0, 0, 0), or (1, 1, 1, 1, 1), where *h*_[*d*]_ = 1 implies a gene is lowly expressed or off, *h*_[*d*]_ = 0 implies a gene is moderately expressed, and *h*_[*d*]_ = 1 implies a gene is highly expressed in cell population *d*. Otherwise, a gene was considered differentially expressed (DE) along the lineage.

As illustrated by the diagrams in Supplementary Fig. S18, one class of consistently expressed genes (1, 1, 1, 1, 1) contains about 10,000 genes that are highly expressed in all the cell types along each lineage. This observation is consistent with previous results showing that about half of human or mouse genes are expressed at similar levels in all cell types^37^; this set of constrained genes includes those encoding common cellular (“housekeeping”) functions. Another equally large class of consistently expressed genes (−1, −1, −1, −1, −1) was found on each lineage; these classes contain genes that are not expressed in blood cells. A rich set of distinct classes of differentially expressed genes were observed on each lineage. One class showed a dramatic increase in expression during erythroid maturation, which included erythroid marker genes *Alas2, Hba-a1, Hba-a2*, and *Gata1*. Similarly, three classes showed substantial induction during one or both of monocyte and neutrophil differentiation; these classes include myeloid marker genes *Cxcr2, C5ar1, Mpo, S100a8*, and *S100a9*. In contrast, no class of genes showed a dramatic induction to high expression levels during megakaryocyte differentiation, which is consistent with previous analyses showing similar gene expression patterns between multilineage progenitor cells and megakaryocytes^38^. In total, our results identified 2,242 DE genes along the erythroid lineage, 2,073 along the megakaryocytic lineage, and 2,376 along the myeloid lineage. Overlap of DE genes across lineages is diagrammed in Supplementary Fig. S19.

A common, alternative approach to this sort of analysis task is to apply a series of pairwise differential expression analyses along each lineage with standard software such as DESeq2^21^, then take the union of all DE genes across the analyses. We implemented this strategy using DESeq2 with FDR ≤ 0.01 and obtained 6,883 DE genes across the erythroid lineage, 7,458 across the megakaryocytic lineage, and 6,863 across the myeloid lineage. The number of DE genes called by DESeq2 was about one third of all input genes for each analysis, and about 3 times more than the number of DE genes identified by CLIMB. We also applied SCREEN to identify DE genes along each lineage, and found that SCREEN systematically reported lower precision in identifying lineage-related GO terms than both CLIMB and DESeq2 (Supplementary Fig. S20). All differential genes identified by CLIMB and DESeq2 are provided in Supplementary File 2.

The large number of DE genes returned by DESeq2 raises questions about the specificity of this approach in pinpointing genes relevant to differentiation. To probe whether DESeq2 is exhibiting low precision or CLIMB exhibiting low power, we first ran gene ontology (GO) enrichment analyses for each lineage^39,40^. Some enriched GO terms from the CLIMB analysis of each lineage are in Table 1. Meanwhile, with the exception of the myeloid analysis, the DESeq2 gene sets were not enriched in lineage-specific GO terms (Supplementary Files 3-8). The abundance of CLIMB’s enriched hematopoiesis-specific GO terms further suggests that, though CLIMB identifies far fewer DE genes than DESeq2, CLIMB is more precise in identifying key genes relevant to cell development and differentiation. See *Simulations and comparisons* to see further investigation of this claim.

**Table 1:**
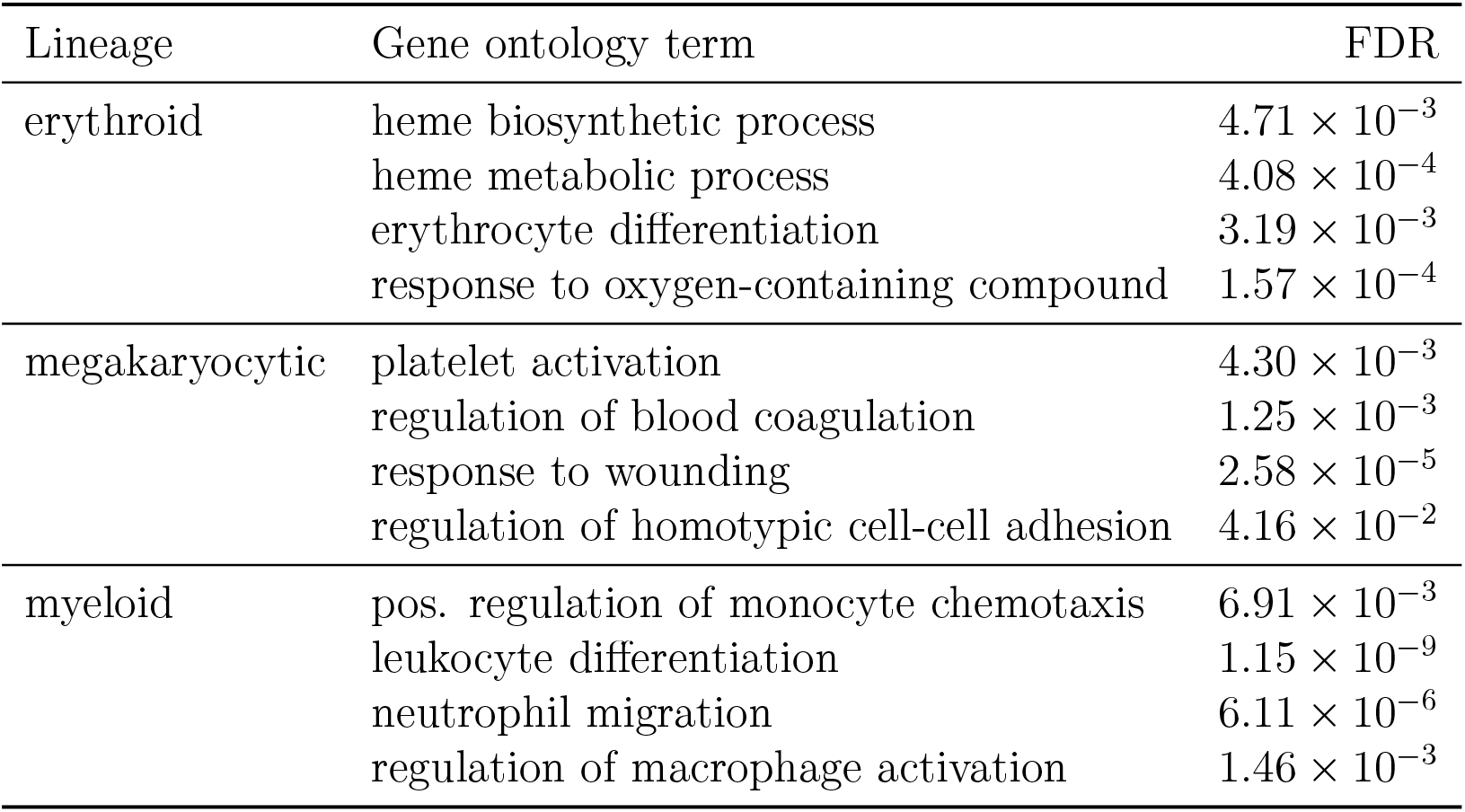
Lineage-specific differentially expressed genes identified by CLIMB are enriched in gene ontology terms related to terminal cell function.

To more directly compare CLIMB and DESeq2, we partitioned DE genes into three categories, namely, differentially expressed genes specific to CLIMB, DE genes specific to DESeq2, and DE genes in the intersection of both methods for each lineage (Fig. 4a), and ran GO analyses on these sets. We noticed that genes identified as DE by both CLIMB and DESeq2 are enriched in many hematopoietic-related terms, while DESeq2-specific genes are enriched for many terms related to general cell function. In each lineage, DESeq2-specific genes are highly enriched for functions that are not specific to hematopoietic cells; CLIMB-specific genes in general are not highly enriched for these same terms. Genes identified by both CLIMB and DESeq2 and CLIMB-specific genes are more frequently enriched for hematopoietic-specific functions (Fig. 4b). The result that DESeq2’s significant gene sets are only enriched in hematopoiesis-related GO terms after intersection with CLIMB’s significant gene sets demonstrates that CLIMB is a powerful and more precise approach to multi-condition differential gene expression analysis when compared to DESeq2 applied in a series across multiple conditions. CLIMB is also a sensitive tool for finding differentially expressed genes, even detecting low-level but differential expression during erythroid differentiation of some genes associated with functions in myeloid cells, in which they are expressed at substantially higher levels (Fig. 4b, Supplementary Fig. S21).

**Figure 4:**
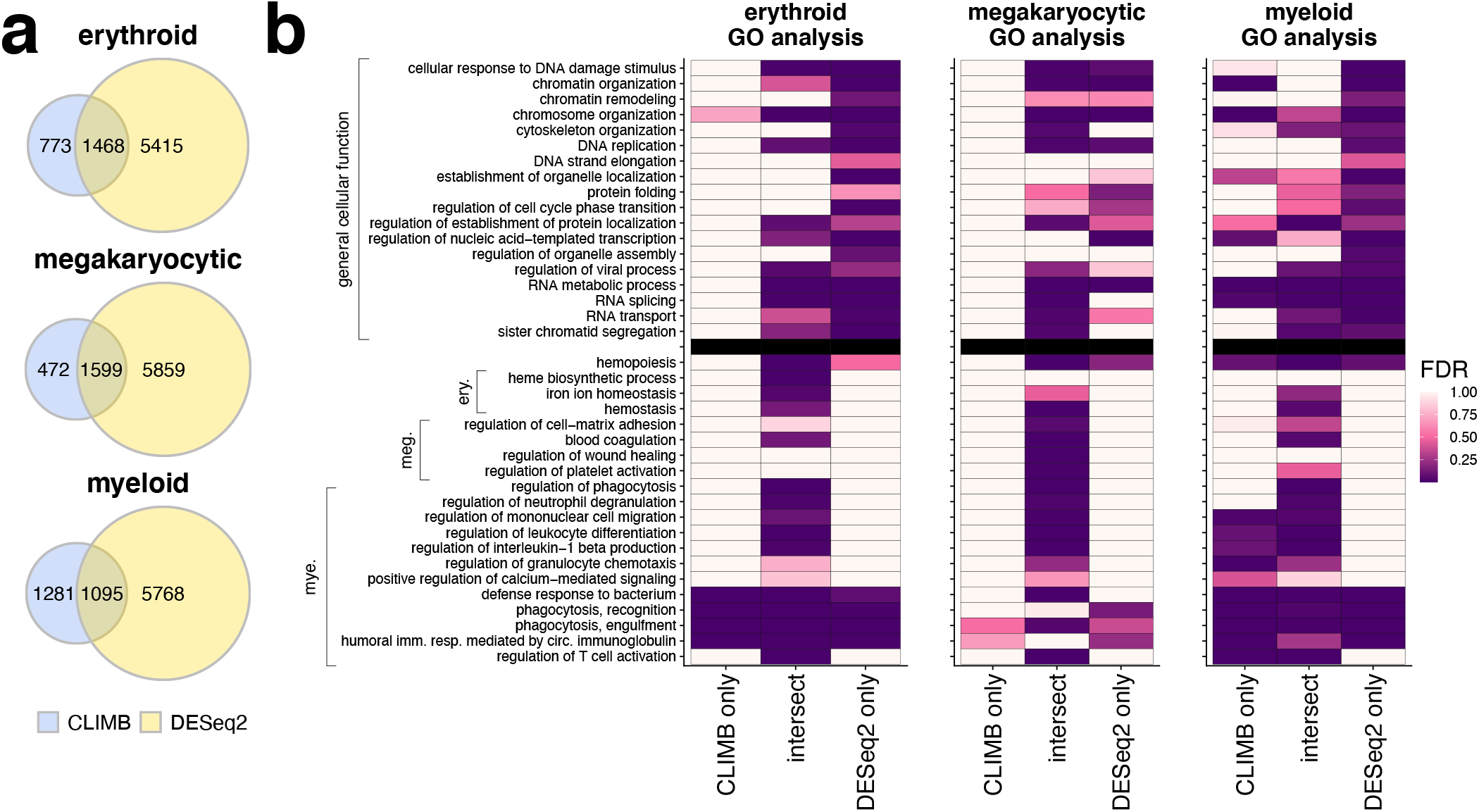
Comparison of differentially expressed genes identified by CLIMB and DESeq2. **a**, Venn diagrams displaying overlap of differentially expressed genes identified by both methods across all analyses. **b**, Significance of enrichment of GO terms in gene sets specific to CLIMB, specific to DESeq2, and in the intersection of both methods, for each studied lineage. Presented GO terms are organized according to knowledge-driven labels. Non-hematopoietic terms related to general cell function are above the black line. Hematopoietic-related terms, grouped according to lineage-specific function, are below the black line.

#### CLIMB latent association labels describe patterns of expression across cell differentiation

Next we used CLIMB to further probe specific gene expression patterns of interest. For example, in the erythroid analysis, 559 genes fell into the (−1, −1, −1, −1, −1) class. This class describes genes with little to no expression in the LSK, CMP, and MEP cell populations, but high expression in the CFUE and ERY cell populations. This gene set is enriched for GO terms such as erythrocyte development (FDR= 5.11 × 10^−7^), iron ion homeostasis (FDR= 9.46 × 10^−3^), and hydrogen peroxide metabolic process (FDR= 1.96 × 10^−2^). Cases of enrichment for terms related to other cell types may result from a process initially discovered in the other cell type being present also in the cell type of interest.

As another example, the 298 members of the (0, 0, 0, −1, −1) class from the myeloid lineage, corresponding to genes that are moderately expressed in LSK, CMP, and GMP cell populations, but lowly or not expressed in monocyte and neutrophil cell populations, are enriched for several GO terms concerning cell fate determination, such as microtubule cytoskeleton organization (FDR= 1.36 × 10^−5^) and mitotic cell cycle process (FDR= 4.42 × 10^−12^). Meanwhile, the 467 members of the (−1, −1, −1, −1, 0) class, corresponding to moderate gene expression specific to neutrophils, are enriched for GO terms immunoglobulin mediated immune response (FDR= 2.47 × 10^−20^), defense response to bacterium (FDR= 2.59 × 10^−20^), and immune response-activating signal transduction (FDR= 4.92 × 10^−25^). Moreover, the 777 members of the (−1, −1, −1, 0, −1) class, corresponding to genes exhibiting moderate expression specific to monocytes, are enriched for the GO terms for the production of tumor necrosis factor and interleukins 1, 6, and 12, as well as the regulation of mast cell activation (FDR= 1.24 × 10^−2^). Taken together, these results demonstrate that CLIMB’s utility goes beyond lineage-specific differential gene expression analysis; the individual latent classes also describe interpretable gene expression patterns.

### ENCODE DNase-seq

As part of the ENCODE project, Meuleman *et al*.^41^ studied DNase-seq in 733 human cell populations, partitioning accessible sites into 16 major groups of cellular accessibility patterns via non-negative matrix factorization (NMF). NMF extracts additive factors across all samples that, when combined, approximate primary signal patterns in the data. With a 38-sample subset of these data, we sought to examine how classes of chromatin accessibility patterns identified by CLIMB relate to differential transcription factor (TF) binding across cell populations, and how these results differ from those extracted via NMF. We applied NMF as before^41^ to a binarized version of this 38-sample subset, and selected an optimal number of 10 factors with NMF (Supplementary Fig. S22a). We merged classes identified with CLIMB into 10 parent groups to match NMF.

#### CLIMB extracts factors of cell type-specific accessibility patterns

We used the class mean and first two principal components (PCs) of the class covariance matrix to extract information from each CLIMB class. These quantities can be interpreted similarly to factors identified with NMF, capturing different cell type-specific accessibility patterns (Fig. 5a). For example, class 4 captures signals specific to K562 cells, while class 5 captures signals specific to T2 helper cells, GM12865, dendritic cells and classical monocytes. Class 7 contains accessible sites absent in differentiated erythroid, K562, HAP1, and fetal liver hepatic cells, yet present in all others. Classes 1 and 3 both correspond to loci broadly accessible across cell populations, although interestingly they bear striking differences in their PCs. Class 1 shares much with class 7, indicating sample-invariant trends in the first PC. The second PC splits CD34+ hematopoietic progenitors, classical monocytes, T helper cells, and regulatory T cells from CD4+ and CD8+ T cells and B cells. Meanwhile, the first PC of class 2 indicates nearly half of the variance in this class is explained by signals in lymphoid cells, while the second PC splits undifferentiated from differentiated CD34+ cells. Such differences suggest the possibility for functional differences inherent in these two different classes of accessible loci.

**Figure 5:**
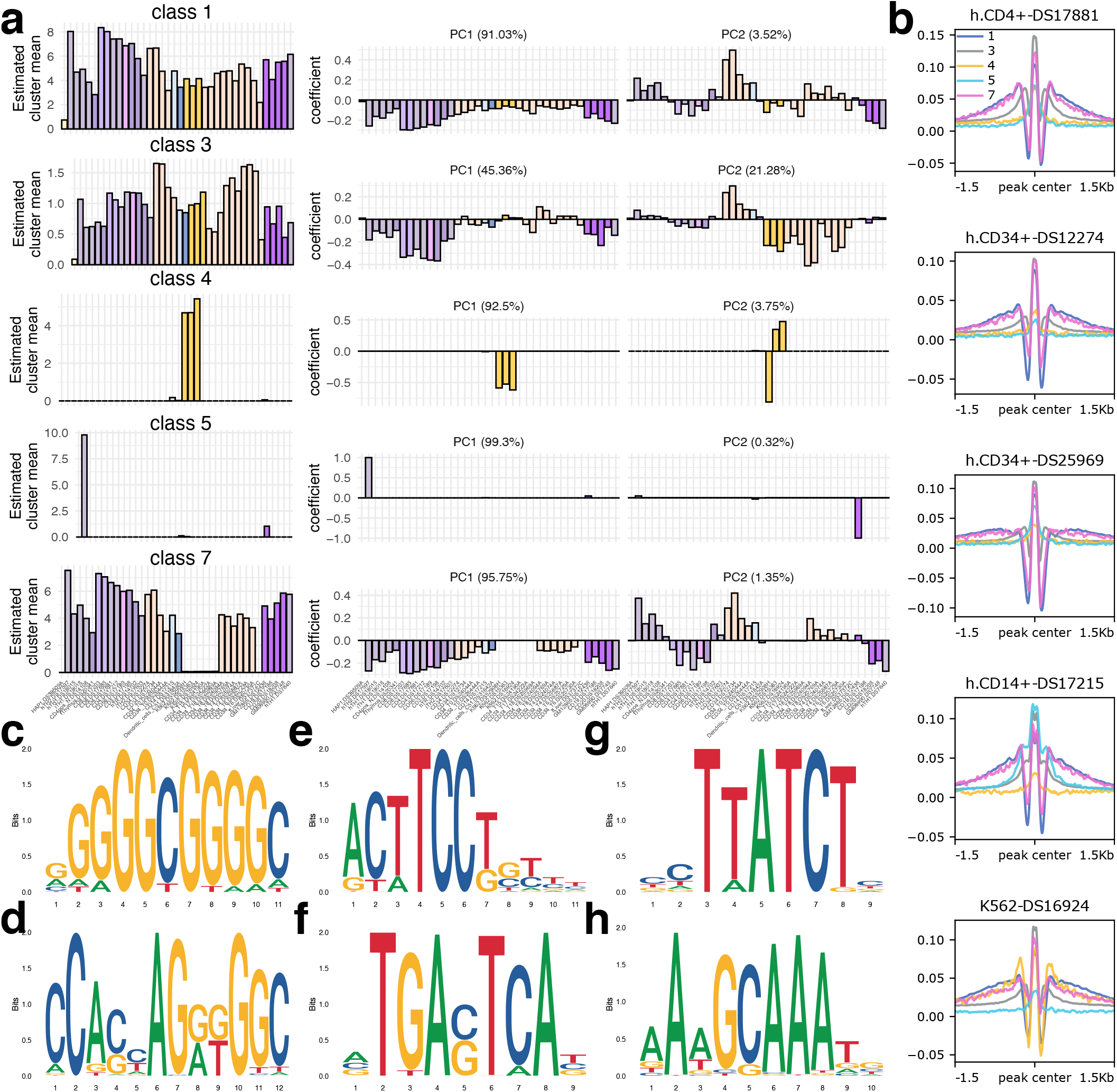
CLIMB identifies patterns of chromatin accessibility across hematopoietic cells relating to different transcription factor binding signatures. **a**, CLIMB’s estimated class means across all 38 cell populations are shown alongside the first two sets of eigenvector coefficients of the estimated class covariance matrices. Cell samples are ordered based on their similarity according to model output. **b**, Footprint signatures for the 5 shown classes in a subset of examined cell populations. **c-f**, Top 4 enriched motifs in class 1. **g**, Most enriched motif in class 4 **h**, Enriched motif specific to class 5.

Because class 3 appeared distinct from classes 1 and 7 based on the PCs, we investigated these loci further. We classified each locus into a PC1 or PC2 group using the PC scores based on the first two PCs, which assess how well each PC describes the signal patterns across all samples for each locus. These subgroups of class 3 contain 37,746 and 29,759 loci for PC1 and PC2, respectively. We used GREAT to identify significant biological processes associated with each set of loci. Interestingly, we found that all top terms in the PC1 group relate to either brain stem morphogenesis or male gamete function. Many of the top terms from the PC2 group relate to lymphoid cells, such as B cell adhesion (FDR=8.06 × 10^−7^), negative regulation of eosinophil migration (FDR=1.79 × 10^−5^) and T cell antigen processing and presentation (FDR=1.44 × 10^−4^). Additionally, the median signal among lymphoid cells in the PC2 group (1.06) is significantly higher than that in the PC1 group (0.286, two-sided Wilcoxon signed rank test, *P <* 2.2 × 10^−16^). The difference in median signal between these two groups is much less for the non-lymphoid cells (0.659 and 0.935 for PCs 1 and 2). This suggests that PC1 describes signals that are more variable in lymphoid cells, while PC2 captures signals that are stronger and more consistent in those same cells.

#### Classes of chromatin accessibility differentiate modes of TF occupancy

Vierstra *et al*.^42^ studied functional changes in regulation by TFs using TF footprinting data. They showed that footprint widths track closely with both the length of the contained canonical TF binding sequence(s) as well as the number of bound TFs, identifying sources of cell type-specific regulation. We interrogated whether classes of accessibility patterns identified by CLIMB and NMF relate to functional differences as captured by TF footprinting.

CLIMB classes bear striking TF footprinting patterns across different cell populations (Fig. 5b). For example, K562 shows a dramatic change in signal for class 4, aligning with the signal enrichment in Fig. 5a. As another example, class 5 has a relatively weak TF footprint signal in all shown cell types except the CD14+ cell; though the mean signal is dominated by a single T2 helper cell for this class, it is also specific to the myeloid CD14+ and dendritic cell populations. In contrast, though NMF identified 10 biologically interpretable classes, several of which have a counterpart class identified by CLIMB, differences between classes are not evident based on footprints (Supplementary Fig. S22). This suggests a greater sensitivity by CLIMB to separate weak patterns from strong, covarying ones.

We used STREME^43^ to interrogate enrichment for canonical TF recognition sequences in each of these classes (Fig. 5c–e). Given that classes 1, 3, and 7 each contain broadly accessible sites, we expected to find enrichment for sequences associated with TFs important for general cellular maintenance. As an example, the top 4 sequences from class 1 (Fig. 5c) include the recognition sequences for Sp1 and KLF families, CTCF, and the ETS and AP1 families (Fig. 5c-f, respectively), though these motifs are enriched in all 3 classes. Further, the most significantly enriched motif in class 4 is the recognition sequence for the GATA proteins (Fig. 5g), while class 5 is uniquely enriched in the non-canonical recognition sequence for the octamer TFs (Fig. 5h). The presence of class-specific motifs further suggests that classes of chromatin accessibility patterns identified by CLIMB relate to differentially regulated genomic regions.

## Discussion

We present a new method, CLIMB, for joint analysis of genomic data collected from multiple experimental conditions. CLIMB gains statistical power to uncover biologically relevant signals by providing a means to extend typical pairwise analyses to higher dimensions. Moreover, when compared against methods designed for a higher-dimensional setting, we demonstrated that CLIMB remains powerful, flexible, and interpretable in many contexts.

A major benefit of CLIMB is its ability to describe various patterns of condition-specificty in a mixture with corresponding association vectors that are estimated from the data. The model, aided by these association vectors, is scientifically interpretable. Estimated model parameters can elucidate similarity and interrelationships, and parsimoniously characterize representative association patterns present across experimental conditions. Importantly, the association vectors also serve as the basis for a novel and effective means of testing consistency of signals across several conditions or biological experiments.

Since CLIMB’s mixture modeling framework is quite flexible, it is effective on a wide range of input data, as long as the data can be reported as numerical scores that reflect strengths of association. Though we have focused on specific molecular traits, CLIMB has the potential to be effective in other applications, such as multi-omics molecular QTLs analysis^44^. The current implementation of CLIMB supports no more than a hundred conditions for genome-wide analyses of the size similar to our DNase-seq analysis. Algorithmically faster implementations, such as variational Bayes fitting for the final Bayesian mixture model, will be explored in future studies for supporting larger numbers of conditions.

## Methods

### Constrained mixture model for estimating association vectors

To estimate the association vectors, we consider the following mixture model. Define

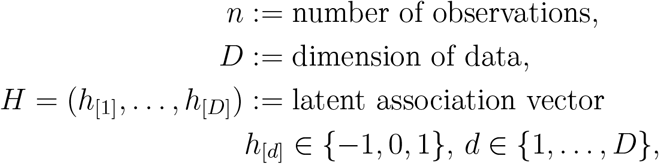

such that the observed data follow the constrained normal mixture model

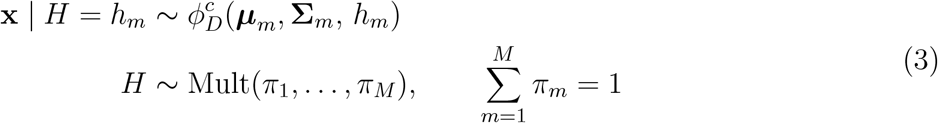

where *h*_*m*_ is the *m*^*th*^ latent class, *m* ∈ 1, …, *M*, and 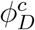 is a *D*-dimensional constrained normal distribution. Note that the number of candidate latent classes *M* changes as our methodology prunes unsupported classes (see *Pairwise fitting* and subsequent methodological steps).

If an observation has association label *h*_[*d*]_ = 1 (*h*_[*d*]_ = −1), this implies that it exhibits a significant positive (negative) association with condition *d*. Otherwise, if an observations has association label *h*_[*d*]_ = 0, this implied that it exhibits a null association with condition *d*. To capture this relationship described by the association vectors, we set the following constrains on 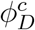:

1. Null associations in dimension *d* are assumed to follow the standard normal distribution (*µ*_*d*_ = 0, *σ*_*ds*_ = 1).
2. Non-nulls that have a positive (negative) association in dimension *d* have a strictly positive (negative) mean in dimension *d*.
3. Nulls in one dimension do not correlate with non-null associations in any other dimension (∑_*rt*_ = 0 ∀*t* ≠ *r* if either *h*_[*r*]_ = 0 or *h*_[*t*]_ = 0).
4. Non-nulls that show concordant (discordant) associations across dimensions—i.e., *h*_[*r*]_ = *h*_[*t*]_ (*h*_[*r*]_ = −*h*_[*t*]_) where *h*_[*r*]_ ∈ {−1, 1}—are positively (negatively) correlated, that is, ∑_*rt*_ > 0 (∑_*rt*_ < 0).

A 2-dimensional visualization of these constraints is in Fig. 1a. Though these constraints are desirable for interpretability, imposing them through latent association vectors leads to computational difficulties as the number of dimensions grows because there are 3^*D*^ possible configurations of the latent association vectors. We thus developed CLIMB, a modeling strategy designed to circumvent the computational intractability that arises under these circumstances. We now describe the steps of CLIMB in greater detail.

### Detailed CLIMB procedure

#### Pairwise fitting

Composite likelihood (CL) methods^45^, which have been reviewed extensively^46^, are computationally efficient modeling approaches that approximate the joint data model by making certain conditional independence assumptions. CL methods are frequently utilized in statistical literature. For instance, they can simplify a genetic model of recombination rates by assuming conditional independence given nearest neighbors along the genome^47^, or sidestep specifying a complex joint likelihood in favor of a product of bivariate models^48^. CL estimators are consistent, though they exhibit some loss in efficiency.

We are seeking to reduce model complexity in the number of latent classes by limiting the dimension of the data through pairwise CL. Let Ω = {(**X**_*·*1_, **X**_*·*2_), …, (**X**_·D−1_, **X**_·D_)} be the set of all pairs of dimensions of **X**_*n*×*D*_, giving 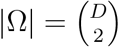. The pairwise CL is

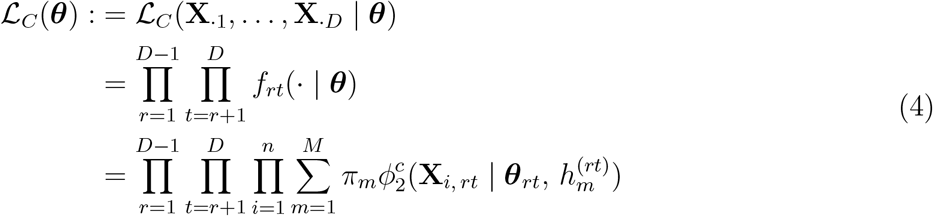

where **X**_·*rt*_ is the *n* × 2 matrix of observations from dimensions *r* and *t*, 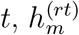 is the *m*^th^ class in the set of all possible 2-dimensional latent association vectors *h*_*rt*_ between dimensions *r* and *t*, and ***θ***_*rt*_ := {***µ***_*rt*_, ∑_*rt*_}is the parameter vector describing the normal mixture between dimensions *r* and *t*. The signs of all elements of ***θ***_*rt*_ are governed by *h*_*rt*_, as in Equation 2. Note that for each pair in D, each pairwise model, *f*_*rt*_, is computationally tractable. This style of pairwise CL, termed “pairwise fitting”, has been utilized most frequently to alleviate computational difficulty when analyzing survey data with multivariate responses^49,50,51,52,53^. Because each dimension appears in *D* 1 different pairwise fits, the mean and variance of each class are estimated *D* 1 times, leading to *D* 1 not necessarily equal estimates for the same mean and variance. It has been shown that, though these pairwise estimates are redundant and not necessarily concordant, they carry useful information about the true parameters^53^. Thus we will recycle these estimates to inform the priors in the final step of our procedure (see *An empirical Bayesian model*).

Fitting each pairwise model *f*_*rt*_ amounts to fitting a finite normal mixture model arising from 9 classes described by latent association vectors *h* ∈ **ℋ**_*rt*_ where

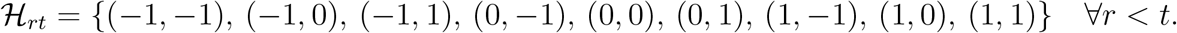

However, since the total number of latent classes in the full model is less than 3^*D*^, we expect that the true number of latent classes in some, if not all of the pairwise fits, is less than 9. Accordingly, for each pairwise fit, we perform model selection to filter out unsupported classes at the pairwise level using a previously described penalized maximum likelihood approach^19^. This method provides an automated model selection procedure for normal mixture models with theoretical guarantees of consistency in selecting the correct number of clusters (see *Model selection details*).

#### Construction of *D*-dimensional association labels

Next, we assemble the list of candidate *D*-dimensional latent association vectors by concatenating all the pairwise association vectors of adjacent dimensions estimated in the previous step. Only association vectors that are on this candidate list are retained for downstream analyses. Example 1 shows a simple example for a 3-dimensional dataset.

*Example 1:* Let ℋ_*rt*_ ⊆ **ℋ**_*rt*_ be the set of 2-dimensional latent association vectors present in a model of dimensions *r* and *t*. Now, consider a three-dimensional dataset, where latent association vectors (−1, 0) ∈ ℋ_12_ and (0, 1) ∈ ℋ_23_. These two association vectors suggest that some observations belong to the null class in dimension 2, and that some of these observations exhibit negative signals in dimension 1 [since (−1, 0) ∈ ℋ_12_], and positive signals in dimension 3 [because (0, 1) ∈ ℋ_23_]. Thus, the data support that (−1, 0, 1) remains a candidate *D*-dimensional latent association vector.

To perform this task computationally efficiently, we construct a directed acyclic graphical representation of the pairwise classification results, designed in the spirit of a de Bruijn graph^54,55^. This novel representation allows one to efficiently enumerate all plausible candidate *D*—dimensional latent association vectors in the concatenation by applying a standard graph search algorithm.

Specifically, we denote a vertex in the graph as (*d, a*), representing a possible association, *a*, at a given dimension, *d*. For a model with *D* dimensions, the graph has *D* layers and 3 possible associations at each layer: −1, 0, and 1. A pictorial view is in Supplementary Fig. S23. We write the vertex set as the collection of all ordered pairs

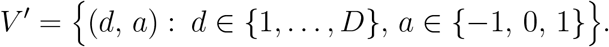

The edge set is defined as

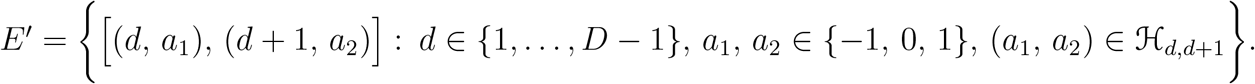

The final graph also contains dummy source and target nodes *S* and *T*, such that the final vertex set *V* = *V*′ ⋃ {*S, T*}. The source node has edges pointing to all nodes in layer 1, while each node in layer *D* has an edge pointing to the target node. The final edge set is then defined as

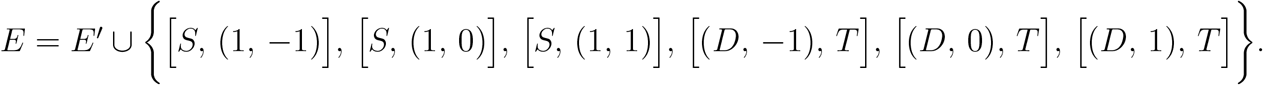

Once the graph is constructed, depth-first search with backtracking^56^, a graph search algorithm that enumerates all paths in a graph from a given source node to a given target node, is used to enumerate all paths from *S* to *T*. Each path contains one node from each of the *D* layers plus the source and target nodes, and has *D* +1 edges of the form

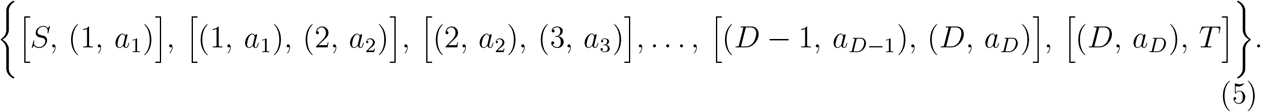

This path corresponds to the latent association vector (*a*_1_, …, *a*_*D*_).

#### Pairwise fit-based pruning

The initial construction of the graph in *Construction of D-dimensional association labels* only uses output from the *D*−1 pairwise fits between dimensions and +1 for *d* ∈ {1, …, *D*−1}. Certain paths may be incompatible with the remaining 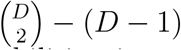 fits. We next remove these paths from the candidate list by checking for incompatabilities, in a manner similar to the continuation of Example 1 below.

As shown previously, (−1, 0, 1) was identified as a candidate *D*-dimensional latent association vector. If (−1, 1) ∉ ℋ_13_, then the latent class (−1, 0, 1) is discarded from down-stream analysis. This is because ℋ_13_ shows that (−1, 0, 1) is incompatible with the pairwise findings.

The graph-based enumeration and pruning algorithm is a deterministic procedure that is guaranteed to produce a list of candidate latent classes that includes all true underlying classes with the possibility of additional empty classes, assuming the correct pairwise classes were estimated (Proposition 1). Further, the results are not affected by reordering of the dimensions (Proposition 2, see Supplementary Section 1 for formal proofs).

### Mixing weight-based class pruning

Since the pairwise fit-based class pruning procedure is *conservative*, some remaining candidate classes still may not be present in the data (e.g, the (0, 0, 0) latent association label in the toy example in Fig. 1). To prune these classes, we estimate the weights of the remaining classes based on the pairwise fitting, and remove those whose weights are near zero. To elucidate which classes are unsupported, we devise an estimator that measures the concordance between the candidate list of *D*-dimensional association labels against the pairwise labels for each observation. Intuitively, our estimator is motivated by the assertion that if observation **x** belongs to a given class *h*, then **x**’s pairwise latent class assignment *h*^(*rt*)^ should equal (*h*_[*r*]_, *h*_[*t*]_) for most pairs *r* and *t, r < t*. Then, the weight for a *D*-dimensional class can be estimated by computing the proportion of observations that follow the pairwise labels of the D-dimensional association vector closely.

To construct such an estimator, let 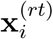 be the sub-vector of the *i*^*th*^ observation vector corresponding to the pairwise fit between dimensions *r* and *t*. Then, let 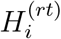 be the pairwise association vector assigned to observation 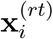. Assuming there are *M* remaining candidate *D*—dimensional latent classes *h*_*m*_, *m* ∈ *{*1, …, *M}*, let 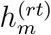 be the sub-vector of *h*_*m*_ corresponding to dimensions *r* and *t*. Then, for a given *D*—dimensional latent class *h*_*m*_, define

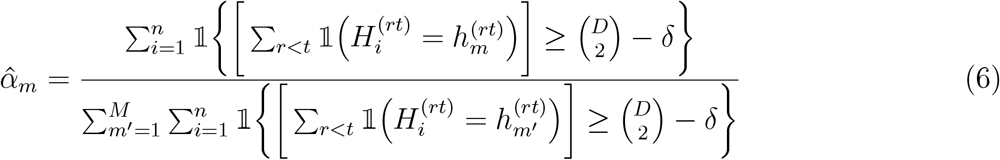

as the normalized proportion of observations whose pairwise class labels are concordant, up to tolerance *δ*, with *h*_*m*_, where 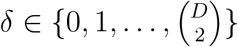, which controls the permitted level of discordance between an observation’s pairwise class labels and its *D*—dimensional latent class. We show that 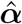 is a reasonable estimator of the proportion of observations belonging to each class *h*_*m*_ given the data (see *Proofs*, Proposition 3).

When the list of remaining candidate latent classes is still large, even after the pruning steps in previous section, 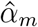 may be very close or exactly equal to 0 for many *m* resulting in a degenerated distribution for these classes in the mixture. This step will remove these classes, guaranteeing that the number of remaining classes *M* is bounded above by the sample size *n*. In practice, we find that this procedure often can reduce *M* to be less than 0.01*n*.

To estimate 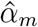, we first obtain each 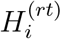 by sampling the pairwise labels of the **x**_*i*_’s according to their posterior probabilities of belonging to each class estimated from the pairwise fits:

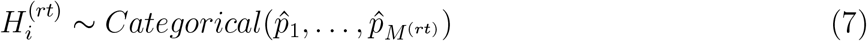

Where 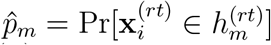, the estimated posterior probability that observation 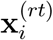 belongs to class 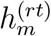 for *m* ∈ {1, …*M*}, and *M* is the number of pairwise latent classes estimated to be present in pairwise fit between dimensions *r* and *t*. Because 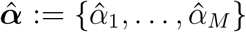 estimates the proportion of observations belonging to each class *h*_*m*_,*m* = 1, …, *M*, we treat 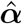 as the prior probabilities for the class mixing weights in the *D*—dimensional model in the next and final step of CLIMB (see next section).

The number of observations needed to obtain a good estimate 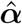 is affected both by the dimension of the data and the accuracy of estimates made during pairwise fitting. For datasets with well-separated clusters, a more stringent *δ* (i.e. 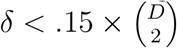) is recommended, whereas a relaxed *δ* (i.e. 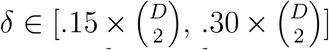) is more suited for datasets with less separated clusters to avoid removing true classes that are small in size. This heuristic guide may be refined by then selecting *δ* within this range where *M* remains constant for *δ′* ∈ {*δ δ*+ 1, …, *δ* + *c*} for some *c* ≥ 1. While this step of our methodology requires user input, it requires similar levels of user input as in existing methods.

### An empirical Bayesian model

With the steps described thus far, we are able to pare down the number of latent classes to a more computationally manageable size for regular mixture modeling. Next we reestimate the parameters in the *D*-dimensional model (1) using an empirical Bayesian approach, recycling the pairwise estimates as prior hyperparameters. We employ the following hierarchical structure to represent the constrained mixture model:

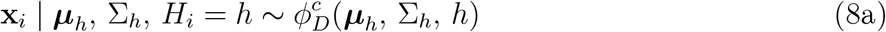

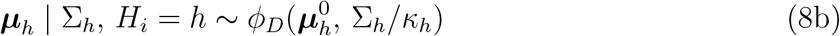

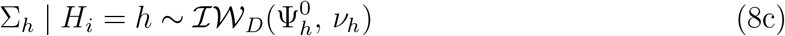

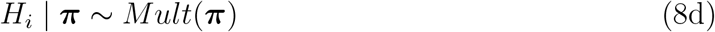

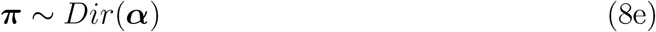

Quantities ***µ***_***h***_, ∑_*h*_∀*h* and ***π*** are estimated using MCMC. The remaining terms *κ*_*h*_, 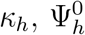, and *ν*_*h*_∀*h* and ***α*** are hyperparameters.

This sort of representation incorporates typical prior distributions and a constrained likelihood model, and has been exploited frequently^57,58,59^ for its desirable posterior structure which is suitable for Gibbs sampling. Similarly here, by applying the necessary parameter constraints, defined by the latent association vectors, into the data model (Equation 8a), the parameters (***µ***_*h*_, ∑_*h*_) possess the correct constraints in the posterior. That is, ***µ***_*h*_ follows a multivariate truncated normal distribution with truncation points dictated by the constraints defined in (8a), while ∑_*h*_ follows the constrained inverse-Wishart distribution defined presently.

Let be distributed according to a *D*—dimensional constrained inverse-Wishart 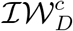 with constraints imposed by latent class *h*, and let ℐ𝒲_*D*_ be an unconstrained *D*—dimensional inverse-Wishart density. Then

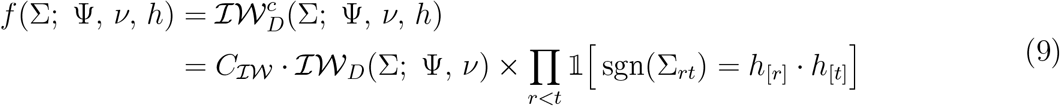

where *C*_ℐ𝒲_ is a normalizing constant.

We do inference on this model using a Metropolis Hastings within Gibbs algorithm, the details of which are in Supplementary File 1. With this procedure, we estimate ***π*** and ***µ***_*h*_ and _*h*_ *h*. An important feature of the mixture model used by CLIMB is that, since the labels *h* explicitly define constraints on the parameters for each class, label switching is not a concern during the inference process. Output from the pairwise fits are used to calculate hyperparameters ***α***, 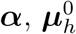, and 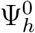 : computation of ***α*** was described in Equation 6, while 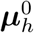, and 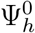 are aggregations of pairwise parameter estimates constructed using a tactic described in *MCMC details*. Parameters *κ*_*h*_ and *ν*_*h*_ ≈ *nα*_*h*_, where *α*_*h*_ is the prior mixing weight for class *h*. We remove classes that satisfy *n–*_*h*_ *D*, since such classes are unlikely to have members, and an inverse-Wishart distribution is singular for these classes.

## Testing consistency of effects

The model fit output from CLIMB can be used to conduct hypothesis tests; in particular, we are interested in identifying consistency of signals across conditions. We propose a new test that generalizes the partial conjunction hypothesis test^60^, a standard hypothesis used for testing consistency, defined as

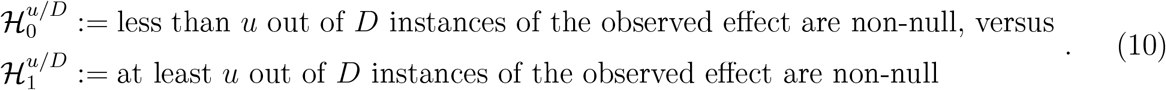

When seeking consistent signals, one may care not only about the significance of the signals, but also the *sign* of the effect. That is, if an observation is significantly positive in one experiment but significantly negative in another, then the observation should not be considered as consistent. Therefore, we propose a simple statistic for assessing the consistency of the sign of the effect across dimensions that generalizes the partial conjunction hypothesis to consider sign:

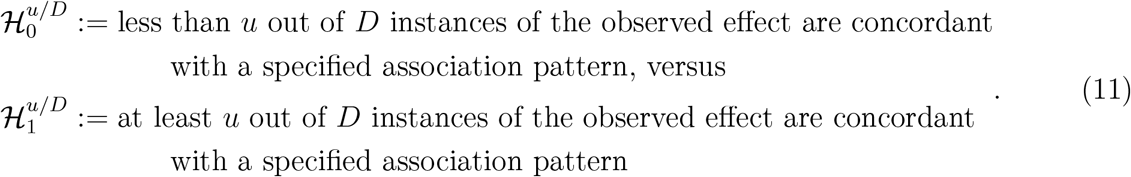

To describe the rejection region (*RR*) for this hypothesis, first define 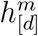 as the *d*^*th*^ element of latent association vector *h*_*m*_. Then,

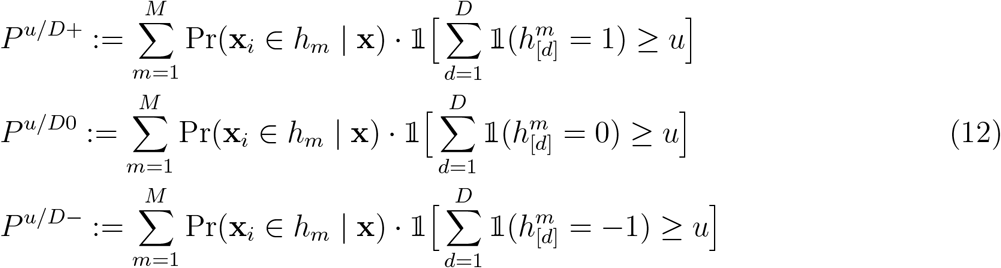

where Pr(**x**_*i*_ ∈ *h*_*m*_|**x**) is the posterior probability of belonging to the class described by association vector *h*_*m*_. We define *P*^*u/D*^ = max {*P*^*u/D*+^, *P*^*u/D*0^, *P*^*u/D*−^}, and *RR* := {**x** : *P*^*u/D*^ *> b*}, where *b* is the confidence threshold of at least 0.5. For each observation, this calculation sums over its posterior probabilities of belonging to classes with association vectors indicating sufficient consistency.

Letting *T* be the number of MCMC iterations retained after burn-in, the quantities in (12) are estimated as

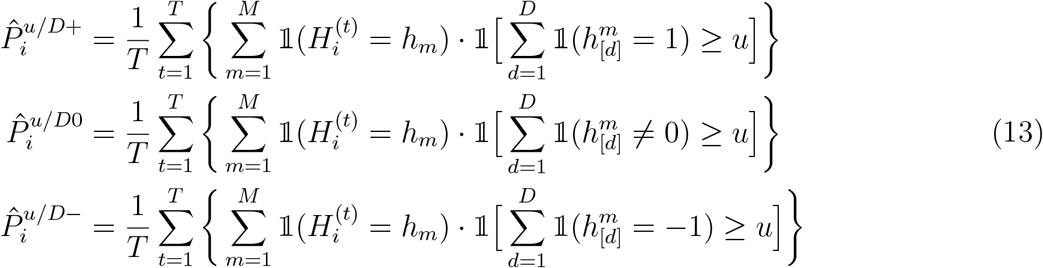

for each observation *i*, leading to 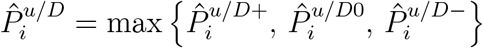, and we reject those **x**_*i*_ with 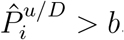. Large values of 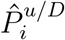 correspond to consistent effects.

This test is flexible, and can be adapted to several purposes. For example, to test the typical partial conjunction hypothesis, one could modify the quantities in Equation 13 to

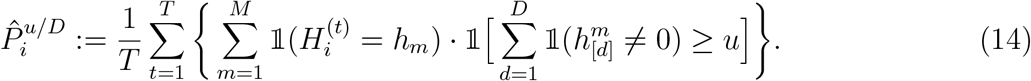

In the analysis of VISION RNA-seq data, we tested for consistency in all −1, 0, and 1 groups. Thus, we applied our statistical test using all quantities in Equation 12 and letting *u* = 5, such that *P* ^5*/*5^ = max {*P*^5*/*5+^,*P*^5*/*50^,*P*^5*/*5−^}. Then, a consistently expressed gene is one that falls within the *RR* := {**x** : *P*^5*/*5^ *>* 0.5}, and all others were called differentially expressed.

## Simulations and comparisons

We used simulations to compare CLIMB to SCREEN^16^ and mash^14^, two methods designed for a similar purpose as CLIMB, as well as DESeq2^21^, a popular method for pairwise differential expression analysis. SCREEN was designed specifically to test for consistent signals across many experiments. Like CLIMB, SCREEN employs a mixture model with classes governed by latent association vectors. SCREEN tackles the issue of computational intractability associated with these classes in two ways. First, it assumes the association vectors to be binary, rather than ternary. This reduces the growth rate of candidate latent classes to 2^*D*^, but comes at the cost of eliminating the method’s ability to detect inverse associations and signs of effects. Second, SCREEN partitions the data’s original conditions into clusters using a network community detection algorithm as an initial step, fitting separate models to each cluster. SCREEN next uses a heuristic to test for consistent signals across all conditions.

Mash, on the other hand, captures the relationship between observations across conditions through the covariances of each cluster in the mixture. Mash assumes the data come from a multivariate normal mixture, restricting each cluster to have zero mean. It sidesteps computational issues by not explicitly specifying the latent association vectors; instead, it models different clusters by specifying a list of candidate covariances which are generated *a priori*. Since the assumed distribution is symmetric and unimodal, model fitting is simplified to a convex optimization problem that can be computed efficiently. Unlike CLIMB, SCREEN, and mash, DESeq2 was not designed for joint testing of conditions, but for testing differential expression pairwise between conditions. In order to simulate data that mimic empirical data, we first fit CLIMB to real datasets (ChIP-seq, differential analysis of RNA-seq, and erythroid lineage RNA-seq data described in *VISION CTCF ChIP-seq*, Shukla *et al*.^61^, and *VISION RNA-seq*, respectively). Parameter estimates similar to those obtained from these model fits were used to simulate *n* = 15, 000, 15, 000, and 21, 303 observations with 18, 11, and 5 dimensions, respectively, according to the constrained normal mixture model in Equation 2 (see Supplementary Tables S4 – S12 and Supplementary Figs. S24 – S26 for specific parameter settings for all simulations). Since DESeq2 requires replicates for each experimental condition, for Simulation 3 we simulated 2 replicates per condition under the same model, but with a correlation of 0.96 between replicates. Since CLIMB is more appropriate for log-transformed RNA-seq data, while DESeq2 is used on counts, i.e. untransformed data, we inputted a rounded 2^*X*^, where *X* is the simulated data, to DESeq2 for analysis. The simulated replicates were averaged before passing to CLIMB.

Like the real datasets, all simulated data contain shared effects that are positively or negatively correlated across dimensions and effects that are unique to one dimension. We applied CLIMB, SCREEN, and mash to Simulations 1 and 2, since these analyses focus on identifying signal patterns across all conditions. We applied CLIMB and DESeq2 to Simulation 3, since the goal of this analysis is specifically to detect differential expression.

Though a usual goal of analyzing these types of data is to uncover the true association patterns of observations across conditions, of all methods, only CLIMB can report the full latent association vectors. To provide a fair comparison among CLIMB, mash, and SCREEN, we test the partial conjunction hypothesis across a series of levels *u*. We do this as SCREEN’s sole functionality is to test this hypothesis, while CLIMB and mash can be utilized for this purpose. By evaluating a range of *u*, we can obtain a comprehensive assessment of each method’s ability to identify consistent signals at different levels of condition-specificity. To compare against DESeq2 in the case of multi-condition differential expression, we identified genes that were differentially expressed along the lineage using the same procedure as in the section *VISION RNA-seq*.

We assessed the performance of each method by comparing the identified consistent signals with the truth and computing the precision and recall at these thresholds (Supplementary Fig. S3 – S5). Precision and recall were computed as

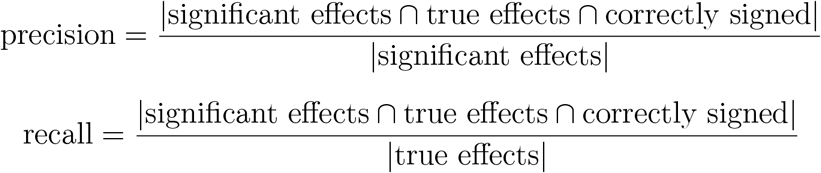

where significant effects are observations that have been estimated to be consistent, true effects are observations that truly are consistent, and correctly signed effects are observations whose true and estimated associations have the same sign. This computation is designed such that an effect correctly identified by an algorithm as significant, but whose effect was missigned, is considered a false positive. The sign requirement was omitted for DESeq2. Analogous precision-recall curves for simulations 1 and 2 that do not incorporate sign information are in Supplementary Fig. S6 and S7.

Separately, we sought to evaluate how accurate CLIMB is at the pairwise fitting step. While the pairwise modeling need not be perfect, it should retain true classes at the pairwise level and have reasonable classification accuracy, such that true classes are likely to be retained in the final model. We assessed CLIMB’s performance during pairwise fitting by calculating classification accuracy and counting the number of missed classes and superfluous classes for each pairwise fit and each simulation (Supplementary Fig. S8). Indeed, CLIMB’s pairwise fitting was more likely to retain extra classes than it was to remove true classes from the model.

## Data availability

The data are available at NCBI’s Gene Expression Omnibus (https://www.ncbi.nlm.nih.gov/geo/)^62^ under accession code GSE156074.

## Code availability

CLIMB is implemented in an R package, freely available on GitHub under an Artistic-2.0 license (https://github.com/hillarykoch/CLIMB).

## Contributions

R.C.H. and Q.L. supervised the project. H.K., G.X, F.Z., Y.W., R.C.H., and Q.L. designed analytical strategies. H.K., C.A.K., G.X., B.G., and R.C.H. analyzed data. H.K. developed analytical tools. C.A.K. performed experiments. G.X. and B.G. administered infrastructure for data storage, quality control, and normalization. H.K., R.C.H., and Q.L. wrote the paper with input from all authors.

## Acknowledgments

The authors are grateful for the support from their funding agencies: NIGMS training grant T32GM102057 (CBIOS training program to The Pennsylvania State University, to H.K.); NHGRI pre-doctoral fellowship 1F31HG010574 (to H.K.); NIGMS grant R01GM109453 (to Q.L.); NIDDK grant R24DK106766 (to R.C.H.). The authors thank Marit Vermunt and Gerd Blobel for generating and sharing the G1E CTCF ChIP-seq data used in this work.

