## Supplementary File 1 for "High-dimensional association detection in large scale genomic data"

### 1 Supplementary Material

#### Model selection details

A finite normal mixture model is estimated with a modified EM algorithm which maximizes a likelihood with a penalty applied to the mixing proportions. For finite bivariate normal mixture model $f(\mathbf{x};\ \boldsymbol{\theta}) = \sum_{m=1}^{9} \pi_m \phi_2(\mathbf{x} \mid \boldsymbol{\mu}_m,\ \boldsymbol{\Sigma}_m,\ h_m)$, denoting by $y_{im}$ the indicator that observation $i$ arises from class $m$, the conditional expected complete data log-likelihood function[63] given parameters $\boldsymbol{\theta} = \{\pi_1,\ \boldsymbol{\mu}_1,\ \boldsymbol{\Sigma}_1, \ldots, \pi_9,\ \boldsymbol{\mu}_9,\ \boldsymbol{\Sigma}_9\}$ is

$$\begin{aligned}
\ell(\mathbf{x} \mid \boldsymbol{\theta}) &= \mathrm{E}\left[\log \prod_{i=1}^{n} f(\mathbf{x}_i;\ \boldsymbol{\theta}) \,\middle|\, \mathbf{x}\right] \\
&= \mathrm{E}\left[\sum_{i=1}^{n} \sum_{m=1}^{9} y_{im} \left\{ \log \pi_m + \log \phi_2(\mathbf{x}_i;\ \boldsymbol{\mu}_m,\ \boldsymbol{\Sigma}_m,\ h_m) \right\} \,\middle|\, \mathbf{x}\right] \\
&= \sum_{i=1}^{n} \sum_{m=1}^{9} \tilde{y}_{im} \log \pi_m + \sum_{i=1}^{n} \sum_{m=1}^{9} \tilde{y}_{im} \log \phi_2(\mathbf{x}_i;\ \boldsymbol{\mu}_m,\ \boldsymbol{\Sigma}_m,\ h_m)
\end{aligned} \tag{15}$$

where $\tilde{y}_{im}$ is the posterior probability that observation $i$ belongs to class $m$ given the observations. We follow Huang et al. (2017)[19], applying a penalty (in our case, SCAD[64]) to the term $\log \pi_m$, to obtain the penalized log-likelihood function

$$\ell_P(\mathbf{x} \mid \boldsymbol{\theta}) = \ell(\mathbf{x} \mid \boldsymbol{\theta}) - n\lambda \sum_{m=1}^{9} \gamma_m \Big[ \log \Big( \epsilon + p_\lambda(\pi_m) \Big) - \log \epsilon \Big] \tag{16}$$

such that the mixing proportions of unlikely clusters are shrunk to zero during the estimation process. Here, $\lambda$ is the tuning parameter, $\gamma_m$ is the number of free parameters in cluster $m$, $p_\lambda$ is the SCAD penalty function applied with parameter $\lambda$, and $\epsilon$ is a small positive number introduced for numerical stability. The SCAD penalty is well-suited for our application since in many genomic applications the classes are not of homogeneous size. While $L_p$ style penalties will over-penalize large values of $\pi_m$, the SCAD penalty yields an unbiased result and will not penalize values of $\pi_m$ that are large enough.

A slight modification to the usual EM algorithm is used to iteratively maximize the penalized likelihood (Equation 16). In the E-step, given current estimate $\hat{\boldsymbol{\theta}} = \{\hat{\pi}_1^0,\ \hat{\boldsymbol{\mu}}_1^0,\ \hat{\boldsymbol{\Sigma}}_1^0, \ldots, \hat{\pi}_9^0,\ \hat{\boldsymbol{\mu}}_9^0,\ \hat{\boldsymbol{\Sigma}}_9^0\}$, we compute the posterior probability that each observation belongs to every class given the data as

$$\tilde{y}_{im} = \frac{\hat{\pi}_m^0 \phi_2(\mathbf{x}_i;\ \hat{\boldsymbol{\mu}}_m^0,\ \hat{\boldsymbol{\Sigma}}_m^0,\ h_m)}{\sum_{m=1}^{M} \hat{\pi}_m^0 \phi_2(\mathbf{x}_i;\ \hat{\boldsymbol{\mu}}_m^0,\ \hat{\boldsymbol{\Sigma}}_m^0,\ h_m)} \tag{17}$$

by some abuse of notation for $M$, which is allowed to shrink during the algorithm's progression. In the M-step, we update (16), which is separable into a component concerning the mixing proportions, and a component concerning the normal parameters. For the former, straightforward calculus with a Lagrange multiplier used to impose a sum-to-one constraint yields a closed-form solution to the maximization as

$$\hat{\pi}_m^\star = \frac{1}{T_m} \sum_{i=1}^{n} \tilde{y}_{im}, \tag{18}$$

where

$$T_m = n + n\lambda\gamma_m \left[ \frac{p'_\lambda(\hat{\pi}_m^0)}{\epsilon + p_\lambda(\hat{\pi}_m^0)} - \sum_{m=1}^{M} \frac{p'_\lambda(\hat{\pi}_m^0)\hat{\pi}_m^0}{\epsilon + p_\lambda(\hat{\pi}_m^0)} \right] \tag{19}$$

and $p'_\lambda$ is the first derivative of the SCAD penalty. For the latter, since we impose application-specific constraints on the model parameters, maximization must be done numerically.

### MCMC details

#### Computing prior hyperparameters from pairwise fits

We recycle information from the $\binom{D}{2}$ pairwise models on order to obtain more informative priors in our model. Here, we detail how we compute the hyperparameters $\{\boldsymbol{\mu}_1, \ldots, \boldsymbol{\mu}_M, \Sigma_1, \ldots, \Sigma_M\}$. For simplicity, we describe only the computation of $\{\mu_1, \ldots, \mu_M, \sigma_1^2, \ldots, \sigma_M^2\}$, which denote the means and variances in just the *first* dimension of each of the $M$ clusters. The computation for other dimensions follows similarly.

In order to begin, we first obtain the pairwise association labels $h_m^{(1d)}, d \in \{2, \ldots, D\}$ for each observation. As discussed in *Mixing weight-based class pruning*, these labels are assigned via multinomial sampling with weights equal to the observation's posterior probability of belonging to each class. Then, for a given cluster $m$ and pairwise fit between dimension 1 and $d$, compute the mean of the subset of data belonging to class $m$ in the first dimension. Then $\mu_1$ equals the average of this value across all $D - 1$ pairwise fits that contain the first dimension. Variances are handled analogously; variance estimates for a fixed class $m$ are combined across pairwise fits not by averaging, but by taking the 75th quantile of the collection of estimates across fits, as averaging often results in covariance estimates that are not positive definite. The off-diagonals of the covariance matrices are computed on the relevant data subsets as well, but do not need to be aggregated. This is because we do not obtain redundant estimates of the off-diagonal covariance terms.

#### Metropolis-Hastings algorithm

Note that typically, since all full conditionals (Equation 26, below) are available for the model in Equation 8, we could do inference using a Gibbs sampler. However, this requires making draws from the truncated multivariate normal and constrained inverse-Wishart distributions. While efficient methods for sampling truncated multivariate normal random variables are available[65,66], unfortunately it is not clear how to efficiently sample constrained inverse-Wishart random variables. As an alternative, we introduce a fast algorithm that is useful for generating positive-definite matrices with constraints on the off-diagonal elements. This algorithm enables us to implement an efficient Metropolis-Hastings step in our MCMC to propose a new covariance drawn with the correct support at each iteration.

Our constrained matrix sampling algorithm modifies the procedure for generating Wishart-distributed random matrices. That is, Wijsman[67] proposed a method for sampling Wishart random matrices based on the Bartlett decomposition of a positive-definite matrix. Consider $\Sigma \sim \mathcal{W}_D(\nu,\, \Psi)$, a Wishart distribution parameterized by $D \times D$ positive-definite scale matrix $\Psi$ and degrees of freedom $\nu$, $\nu > D - 1$. To draw a random sample from this distribution, one computes $U$, the upper-triangular Cholesky factor of $\Psi$, and simulates a $D \times D$ lower-triangular matrix $A$ as

$$A = \begin{pmatrix} c_1 & 0 & \dots & 0 \\ z_{21} & c_2 & \dots & 0 \\ \vdots & \vdots & \ddots & \vdots \\ z_{D1} & z_{D2} & \dots & c_D \end{pmatrix}$$

where $c_i^2 \sim \chi^2_{\nu-i+1}$ and $z_{ij} \sim \phi_1(0,1)$. Then $\Sigma = U^T A A^T U$ is a Wishart-distributed random matrix.

We claim that we can modify Wijsman's simulation procedure to produce positive-definite matrices with the desired support. We notice that we cannot make changes to $U$ without destroying the desired covariance structure of $\Sigma$, nor can we change the diagonal elements of $A$ without risking simulating a matrix that is singular. However, we note that each off-diagonal element of $\Sigma = U^T A A^T U$ can be expanded as

$$\Sigma_{pm} = \Sigma_{mp} = \begin{cases} U_{11}A_{11}\Big(\sum_{i=1}^{p} U_{ip}A_{i1}\Big), & \text{for } m = 1 \\ \sum_{k=1}^{m} U_{km}\left\{\sum_{j=1}^{k} A_{kj}\left[\sum_{i=j}^{p}(A_{ij}U_{ip})\right]\right\}, & \text{for } m > 1 \end{cases}. \tag{20}$$

This representation shows that, due to the triangular structure of both $U$ and $A$, each off-diagonal element $\Sigma_{pm}$ computed using Equation 20 depends only on *certain* off-diagonal elements of $A$ (that is, $A_{21}$ depends on no other off-diagonal element. $A_{31}$ depends only on $A_{21}$, $A_{32}$ on $A_{21}$ and $A_{31}$, and so forth, proceeding in a top-down, row-wise order). Further, each $\Sigma_{pm}$ is a monotonic function of $A_{pm}$.

The monotonicity of $\Sigma_{pm}$ in $A_{pm}$ guarantees that, when $A_{pm}$ is small (large) enough, then $\Sigma_{pm}$ will be negative (positive). Therefore, we can find a upper (lower) bound on $z_{pm}$ such that $\Sigma_{pm}$ will satisfy the given sign constraints. Moreover, given the dependence structure of the off-diagonal elements of $A$, we observe that, conditional on obtaining a satisfactory value of $A_{21}$, we can find a satisfactory value for $A_{31}$, then $A_{32}$, and so on.

For $m < p$, solving Equation 20 for $A_{pm}$ gives

$$A_{pm} = \begin{cases} \frac{\Sigma_{p1} - U_{11}A_{11}(\sum_{i=1}^{p-1} U_{ip}A_{i1})}{U_{pp}U_{11}A_{11}} & \text{for } m = 1 \\ \frac{\Sigma_{pm} - \sum_{k=1}^{m-1} U_{km}\left\{\sum_{j=1}^{k} A_{kj}\left[\sum_{i=j}^{p}(A_{ij}U_{ip})\right]\right\}}{U_{pp}U_{mm}A_{mm}} \\ \quad - \frac{\sum_{j=1}^{m-1} A_{mj}\left[\sum_{i=j}^{p}(A_{ij}U_{ip})\right]}{U_{pp}A_{mm}} \\ \quad - \frac{\sum_{i=m}^{p-1} A_{im}U_{ip}}{U_{pp}} & \text{for } m > 1 \end{cases}. \tag{21}$$

This formula for $A_{pm}$ is used to find upper and lower truncation points for the univariate standard normal random variables that comprise the off-diagonal elements of $A$ such that $\Sigma$ satisfies the desired constraints. The constraint $\Sigma_{pm} < 0$ ($\Sigma_{pm} > 0$) is satisfied when $A_{pm}$ is greater (less) than the result of Equation 21, setting $\Sigma_{pm}$ equal to 0. The full simulation procedure is made explicit in Algorithm 1. Fig. S27 shows some example data simulated using this Algorithm 1 with $\nu = 25$,

$$\Psi = \begin{pmatrix} 2 & 0.3 & -0.6 & -0.8 \\ 0.3 & 1.5 & -0.75 & -0.1 \\ -0.6 & -0.75 & 1.5 & 0.4 \\ -0.8 & -0.1 & 0.4 & 2 \end{pmatrix} \tag{22}$$

,

$$R_{mp} = R_{pm} = \begin{cases} 0, & \text{for } \Sigma_{mp} > 0 \\ -\infty, & \text{for } \Sigma_{mp} < 0 \end{cases} \tag{23}$$

and

$$S_{mp} = S_{pm} = \begin{cases} \infty, & \text{for } \Sigma_{mp} > 0 \\ 0, & \text{for } \Sigma_{mp} < 0 \end{cases}. \tag{24}$$

**Algorithm 1** Sampling $D \times D$ positive-definite matrix $\Sigma$ with desired support

```
Define fixed parameters Ψ, ν, R, and S, where R and S are symmetric D × D matrices of lower
    and upper truncation points for each of the off-diagonal elements of Σ
Compute U, the upper-triangular Cholesky factor of Ψ
for i in 1 to D do
    A_ii ~ sqrt(χ²_{ν−i+1})
for p in 2 to D do
    for m in 1 to p − 1 do
        a_pm ← solution to Equation 21, setting Σ_pm = R_pm
        b_pm ← solution to Equation 21, setting Σ_pm = S_pm
        A_pm ~ TN(0, 1, a_pm, b_pm)
return U^T A A^T U/ν
```

The MCMC proceeds as follows. Initialize all parameters $\boldsymbol{\theta} = \{\boldsymbol{\mu}_1, \ldots, \boldsymbol{\mu}_M, \Sigma_1, \ldots, \Sigma_M, \boldsymbol{\pi}\}$ in the model. At iteration $t$ of the MCMC algorithm, we begin by proposing a constrained covariance matrix cluster-at-a-time. For each cluster $m$, we propose a new covariance, drawn according to $\Sigma_m^\star \sim$ Algorithm $1(\Sigma_m^t, \nu_m^t, h)$, since covariances that do not meet the given constraints will have zero likelihood. $\Sigma_m^t$ is our current estimate of $\Sigma$ at iteration $t$ and $\nu_m^t$ is our current proposal degrees of freedom, log adaptively tuned for each cluster[68]. The acceptance ratio for this proposal is

$$\frac{\prod_{i=1}^{n_m} \phi_D^c(\mathbf{X}_{\cdot i};\ \boldsymbol{\mu}_m^\star,\ \Sigma_m^\star,\ h)^{\mathbb{1}(z_i=m)}}{\prod_{i=1}^{n_m} \phi_D^c(\mathbf{X}_{\cdot i};\ \boldsymbol{\mu}_m^t,\ \Sigma_m^t,\ h)^{\mathbb{1}(z_i=m)}} \cdot \frac{\prod_{i=1}^{n} g(z_i;\ \mathbf{X},\ \boldsymbol{\theta}^\star)}{\prod_{i=1}^{n} g(z_i;\ \mathbf{X},\ \boldsymbol{\theta})}$$
$$\cdot \frac{\phi_D(\boldsymbol{\mu}_m^t;\ \boldsymbol{\mu}_m^0,\ \Sigma_m^\star)}{\phi_D(\boldsymbol{\mu}_m^t;\ \boldsymbol{\mu}_m^0,\ \Sigma_m^t)} \cdot \frac{q(\Sigma_m^\star/(\kappa_m - D - 1);\ \Psi_m^0,\ \kappa_m)}{q(\Sigma_m^t/(\kappa_m - D - 1);\ \Psi_m^0,\ \kappa_m)} \tag{25}$$
$$\cdot \frac{\eta(\Sigma_m^\star \mid \Sigma_m^t)}{\eta(\Sigma_m^t \mid \Sigma_m^\star)}$$

where $g$ is a multinomial density with weights

$$\left\{ \frac{\pi_1 \phi_{(1)}^c(\mathbf{x};\ \boldsymbol{\mu}_1, \Sigma_1)}{\sum_{m=1}^M \pi_m \phi_{(m)}^c(\mathbf{x};\ \boldsymbol{\mu}_m, \Sigma_m)}, \ldots, \frac{\pi_M \phi_{(M)}^c(\mathbf{x};\ \boldsymbol{\mu}_M, \Sigma_M)}{\sum_{m=1}^M \pi_m \phi_{(m)}^c(\mathbf{x};\ \boldsymbol{\mu}_m, \Sigma_m)} \right\},$$

$q$ is an inverse-Wishart density, and $\eta$ accounts for the asymmetric proposal, maintaining detailed balance. The first line of Equation 25 captures the likelihood of the data given the estimated cluster mean vector, covariance matrix, and cluster indicator and the likelihood of the cluster indicators given the data and all estimated means, covariances and mixing weights. The second line captures the priors on the mean and covariance. The final term is proportional the the univariate $\chi^2$ and truncated normal densities that make up the samples from Algorithm 1. This procedure conveniently sidesteps any expensive computation of intractable normalizing constants.

Following this Metropolis step, cluster indicators $\mathbf{z}$, means $\boldsymbol{\mu}$, and mixing weights $\boldsymbol{\pi}$ are sampled using standard Gibbs updates. The full conditionals are written

$$\begin{aligned}
&\boldsymbol{\pi} \mid \mathbf{X},\ \mathbf{h} \sim Dir(\boldsymbol{\alpha} + \boldsymbol{n}), \\
&\boldsymbol{n} \text{ an } M\text{-vector with entries } n_m = \sum_{i=1}^n \mathbb{1}(h_{[i]} = m) \\
(\boldsymbol{\mu}_h,\ \Sigma_h) \mid \mathbf{X},\ H = h \sim \mathcal{NIW}_D^c\Bigg( &\frac{\kappa_h \boldsymbol{\mu}_h^0 + n_h \bar{\boldsymbol{x}}_h}{\kappa_h + n_h},\ \kappa_h + n_h,\ \nu_h + n_h, \\
&\Psi_h^0 + \sum_{i=1}^n (\mathbf{X}_{i\cdot} - \boldsymbol{\mu}_i)(\mathbf{X}_{i\cdot} - \boldsymbol{\mu}_i)^T \mathbb{1}(H_i = h) \\
&+ \frac{\kappa_h n_h}{\kappa_h + n_h} (\bar{\boldsymbol{x}}_h - \boldsymbol{\mu}_h)(\bar{\boldsymbol{x}}_h - \boldsymbol{\mu}_h)^T \Bigg) \\
&H \mid \mathbf{X},\ \boldsymbol{\pi},\ (\boldsymbol{\mu}_1,\ \Sigma_1), \ldots, (\boldsymbol{\mu}_M,\ \Sigma_M)\ \sim \\
&Mult\left( \frac{\pi_1 \phi_{(1)}^c(\mathbf{x};\ \boldsymbol{\mu}_1, \Sigma_1)}{\sum_{m=1}^M \pi_m \phi_{(m)}^c(\mathbf{x};\ \boldsymbol{\mu}_m, \Sigma_m)}, \ldots, \frac{\pi_M \phi_{(M)}^c(\mathbf{x};\ \boldsymbol{\mu}_M, \Sigma_M)}{\sum_{m=1}^M \pi_m \phi_{(m)}^c(\mathbf{x};\ \boldsymbol{\mu}_m, \Sigma_m)} \right).
\end{aligned} \tag{26}$$

### Simulation results

#### Simulation 1: ChIP-seq data

In simulation 1, based on ChIP-seq data, we assess the performance of each method by comparing the identified consistent signals with the truth and computing the precision and recall at a series of identification thresholds. As shown in Fig. S3, CLIMB outperforms mash and SCREEN across

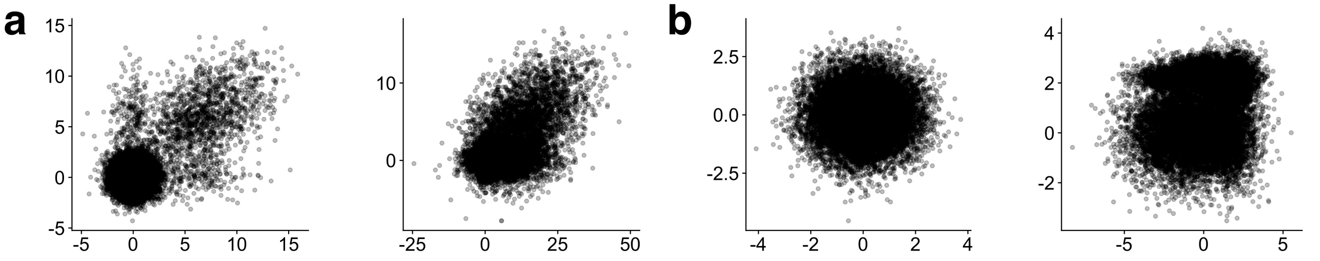


Figure S1: Pairwise plots of data used in simulation studies. **a,** Simulated data between two different pairs of dimensions from simulation 1, resembling ChIP-seq data and **b,** simulation 2, resembling differential analysis of RNA-seq data.

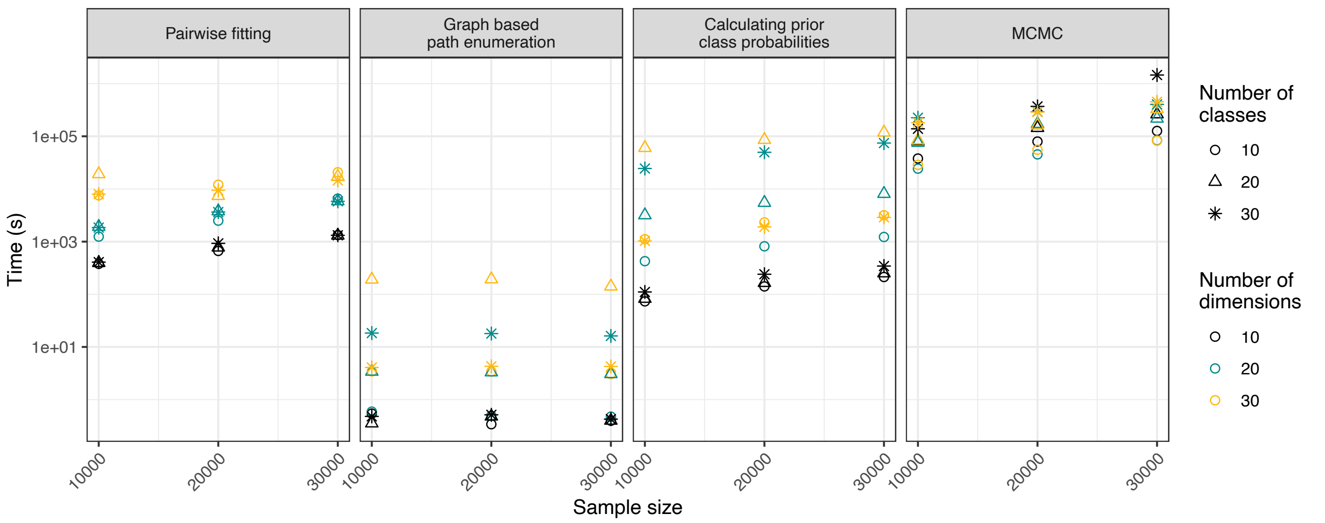


Figure S2: 27 simulations were conducted to investigate the computational demand of each step of CLIMB across different sample sizes, dimensions, and numbers of latent classes. Parameter settings and association vectors were randomly generated in each simulation. The MCMC sampler was run for 15,000 iterations in each simulation. The dimension of the dataset is the major source of increased computational cost. Technically, the MCMC sampler could have a worst-case time complexity of $O(D^5)$. However, the time complexity is also largely driven by how dense the latent association vectors $h$ are with non-zero elements, as this drives the number of parameters needing estimation. In the context we study using this method, there are a large number of zeros in each class (e.g. Supplementary Figs. S10, S18, and S29). Analyses were done on a machine (2.8 GHz Intel Xeon Processor with 256 GB RAM) with 20 cores for parallel processing.

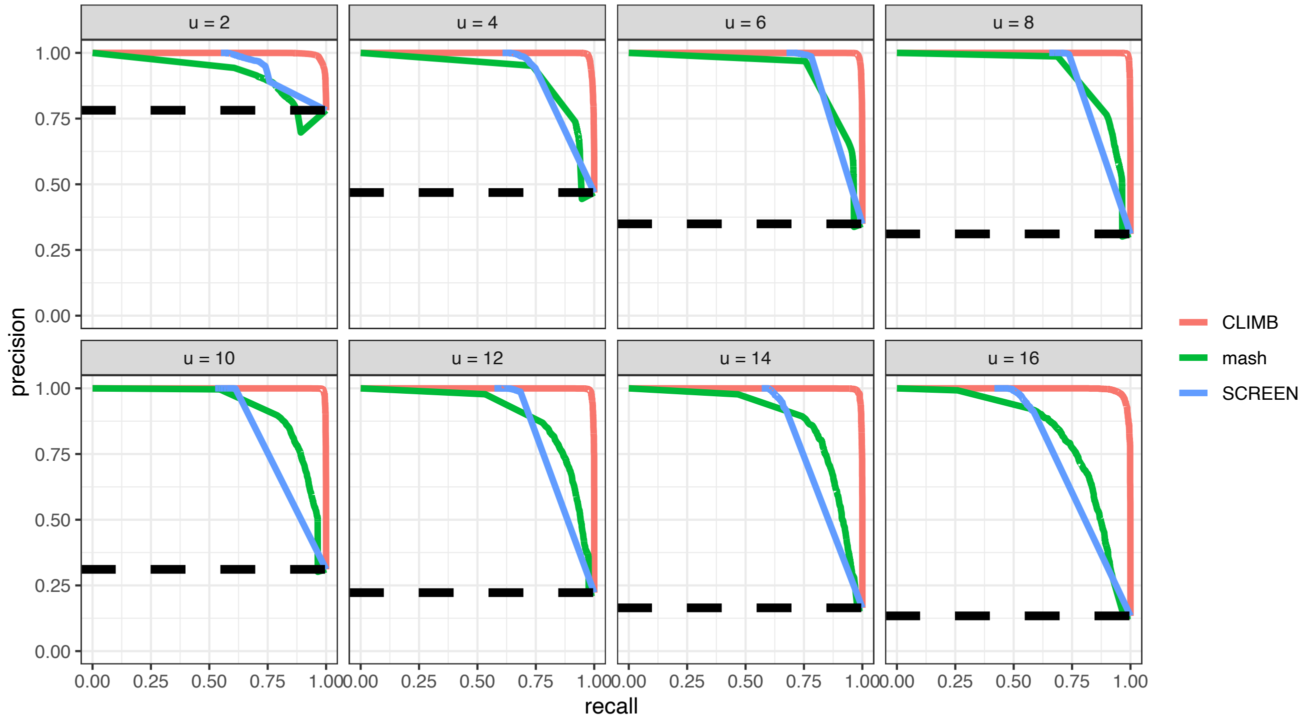


Figure S3: Precision-recall curves for simulation 1, resembling a ChIP-seq dataset. The curves assess CLIMB, mash, and SCREEN's ability to identify genomic loci that are bound by the protein of interest in at least $u$ out of 18 cell types. The dashed black line is the baseline for a random classifier.

all tested $u$, where $u$ is the consistency threshold of the partial conjunction hypothesis (Section *Testing consistency of effects*). We also examined the frequency of missigned signals (Table S1), and found that CLIMB and SCREEN do not missign signals, while mash does. In fact, since SCREEN assumes all signals have positive sign–an assumption that these data satisfy–SCREEN benefits in this comparison. Mash, on the other hand, sometimes missigns effects as negative, and is especially likely to do so for lower $u$. This is likely due in part to the fact that mash expects data to be symmetric and unimodal, but ChIP-seq data do not in general display such structure. CLIMB, on the other hand, can adapt to the ChIP-seq data structure. Indeed a closer examination of the results after pairwise fitting shows that all candidate latent classes with a -1 label are removed during the pairwise fitting step. These results collectively suggest that CLIMB's pairwise fitting step is well-suited to identify pairwise associations in the data, and that the final joint modeling step boosts statistical power to identify consistent signals across $u$.

#### Simulation 2: differential RNA-seq data

Simulation 2, designed to resemble a differential RNA-seq dataset, reveals several contrasting results. Here again, CLIMB outperforms mash and SCREEN (Supplementary Fig. S4). For these data, we noticed that as $u$ increases, accurate classification appears to become more difficult. This is likely due to the fact that few observations are consistent at high thresholds, and the most

| $u$ | 2 | 4 | 6 | 8 | 10 | 12 | 14 | 16 |
|---|---|---|---|---|---|---|---|---|
| CLIMB | 0/10314 | 0/6735 | 0/5055 | 0/4539 | 0/4539 | 0/3211 | 0/2356 | 0/1844 |
| mash | 858/9300 | 326/9088 | 181/8917 | 123/7959 | 39/5752 | 9/4166 | 3/2814 | 0/1718 |
| SCREEN | 0/6602 | 0/4396 | 0/3656 | 0/3129 | 0/2519 | 0/1984 | 0/1468 | 0/930 |

Table S1: Proportion of effects identified as significant at level 0.05 that are true effects, yet incorrectly signed by CLIMB, mash, and SCREEN, from simulation 1.

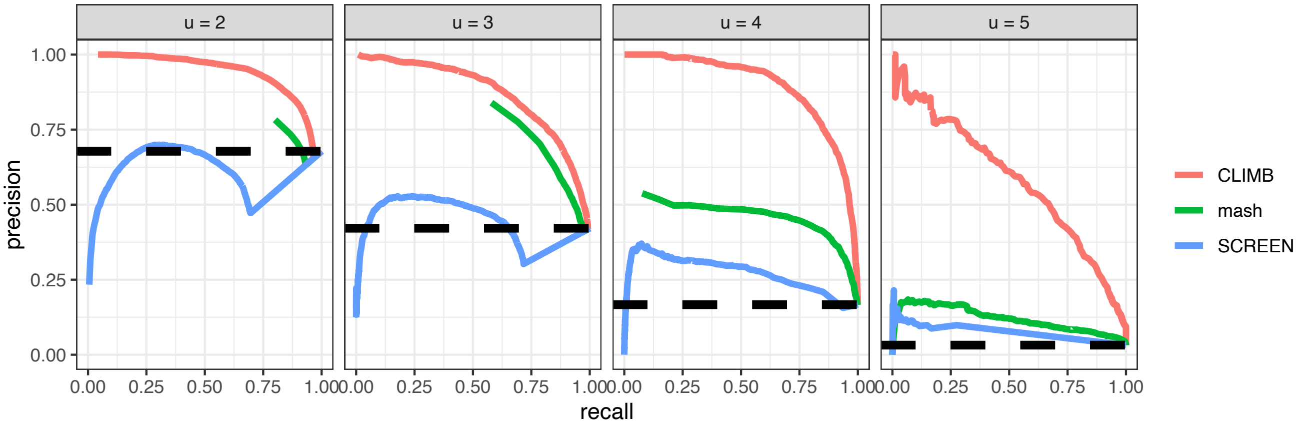


Figure S4: Precision-recall curves for simulation 2, resembling a differential analysis of RNA-seq dataset. The curves assess CLIMB, mash, and SCREEN's ability to identify effects that are consistent in at least $u$ out of 11 tissues. The dashed black line is the baseline for a random classifier.

consistent observations in this dataset are neither well-separated from inconsistent observations nor strongly correlated across dimensions. For example, we find that a mere 480 observations (3.2% of total sample size) are consistent at $u = 5$. The average signal overall is 2.05, and the average signal for the observations that are consistent at $u = 5$ is only 1.61. These features of the data would challenge any method, explaining the precipitous drop in performance of mash and SCREEN at higher thresholds ($u \geq 4$). Due to the stochastic nature of class pruning and modeling employed by CLIMB, this challenging setting can affect CLIMB's performance as well, yielding somewhat differing results for $u \geq 4$ across multiple runs. Still, CLIMB outperforms the other two methods and all methods perform better than a random classifier.

We next investigated the frequency with which each method missigns signals (Supplementary Table S2), and found that CLIMB performs the best according to this metric. Most notably, SCREEN struggles to make accurate inference in this setting. This is because SCREEN does not differentiate signs, thus it identifies many inconsistent effects that appear significant in both the positive and negative directions, as consistent. Since CLIMB and mash model signals in both the positive and negative directions, these methods missign signals far less frequently.

On the other hand, mash reports a very small estimated mixing weight of $2.02 \times 10^{-3}$ for the null class, resulting in almost all observations being called significant, a trend that CLIMB and SCREEN do not exhibit (see Supplementary Fig. S9 for a comparison of null observations across all methods). This indicates that mash is sensitive compared to other methods, rendering it

| $u$ | 2 | 3 | 4 | 5 |
|---|---|---|---|---|
| CLIMB | 9/6059 | 0/2771 | 0/786 | 0/2 |
| mash | 686/15000 | 188/15000 | 4/12425 | 0/5056 |
| SCREEN | 1289/5207 | 119/285 | 0/31 | 0/0 |

Table S2: Proportion of effects identified as significant at level 0.05 that are true effects, yet incorrectly signed by CLIMB, mash, and SCREEN, from simulation 2. There are no truly consistent effects at more than 5 ($u > 5$) conditions.

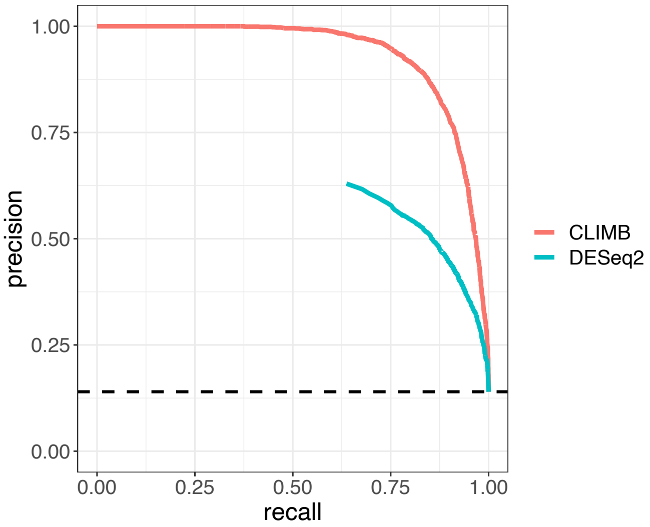


Figure S5: Precision-recall curve for simulation 3, resembling a lineage-specific RNA-seq dataset. The curves compare the performance of CLIMB and DESeq2 at identifying genes that are differentially expressed along the lineage. The dashed black line is the baseline for a random classifier.

inappropriate for testing consistency of signals. CLIMB, meanwhile, demonstrates the strongest performance, based on both the precision-recall curves (Fig. S4) and the proportion of missigned effects (Table S2). Taken together, these results affirm that CLIMB's flexible modeling framework is essential to adapt to complex datasets, and the inclusion of latent association labels reduce oversensitivity when identifying signals.

#### Simulation 3: RNA-seq data across cell differentiation

We set up simulation 3 to mimic an analysis whose goal is to understand how gene expression levels change across cell developmental stages. It supports a claim that CLIMB shows better precision than DESeq2 in the specific setting of multi-condition differential analysis. In particular, though DESeq2 is known to be an effective tool for differential expression analysis between a pair of conditions, the modeling and testing approach, which aggregates pairwise tests across conditions, may weaken the analysis when there are additional conditions. This is a setting where CLIMB can excel by cohesively capturing expression patterns across differentiation.

### Simulation details

In simulations 1 and 2, testing for consistency was carried out for each method using comparable, but not exactly the same, procedures. For CLIMB, we tested for consistency using the procedure described in the Methods (see *Testing significance and replicability of effects*). For SCREEN, we used the test provided by Amar et al.[16] in the SCREEN software. Both tests are similar, in that they use binary (for SCREEN) and ternary (for CLIMB) vectors as class labels of association patterns to determine the probability of a given effect being significant in a given dimension. They are different in that SCREEN clusters the dimensions and thus does not produce $D-$dimensional cluster labels, and that CLIMB requires the additional step of selecting the sign of each effect (Equation 12). Mash, on the other hand, does not define its classes by association patterns dictating the significance and direction of effects. Therefore, we designed a consistency test for mash to be as close to the test used for CLIMB as possible.

The consistency test for mash begins with the quantitites $p_{id}^{+}$ and $p_{id}^{-}$, $i \in \{1,\ldots,n\}$, $d \in \{1,\ldots,D\}$, the probability that effect $i$ in dimension $d$ is non-null positive or non-null negative, respectively. These quantities are readily output by mash. We used these terms to compute $\tilde{P}_i^{u/D+}$ and $\tilde{P}_i^{u/D-}$, the probability that effect $i$ is non-null positive or non-null negative, respectively, in at least $u$ out of $D$ dimensions. These terms are analogous to those used by CLIMB (see Equation 13), and are calculated with the equations

$$\begin{aligned}
\tilde{P}_i^{u/D+} &:= \sum_{d=u}^{D} \sum_{j \in \binom{\{1,\ldots,D\}}{d}} \prod p_{ij}^{+} \\
\tilde{P}_i^{u/D-} &:= \sum_{d=u}^{D} \sum_{j \in \binom{\{1,\ldots,D\}}{d}} \prod p_{ij}^{-}
\end{aligned} \tag{27}$$

by minor abuse of notation as index $j$ in fact refers to a whole set of dimension indices. Again, analogous to the replicability test for CLIMB, we define $\tilde{P}_i^{u/D} = \max\left\{\tilde{P}_i^{u/D+}, \tilde{P}_i^{u/D-}\right\}$ as the probability that effect $i$ is replicable at level $u$. This test comes with the caveat that it assumes the $p_{ij}$'s are independent.

When testing with SCREEN, the sign of the association is always positive. For both CLIMB and mash, the sign of the association must match the direction of association that maximizes $P_i^{u/D}$ and $\tilde{P}_i^{u/D}$, respectively. Precision-recall curves for simulations 1 and 2 constructed without considering the sign of association are presented in Supplementary Fig. S6 and S7.

### Implementation details

#### Running CLIMB

*Parameter constraints during pairwise fitting.* During the pairwise fitting step of CLIMB, we impose additional constraints on $\boldsymbol{\theta}_{rt}$ which are relaxed in the final, $D-$dimensional model. For any pairwise fit between dimensions $r$ and $t$, these constraints are

1. all non-null elements of $\boldsymbol{\mu}_h$ are equal in magnitude $\forall h \in h_{rt}$,

2. all non-null variance terms of $\Sigma_h$ are equal $\forall h \in h_{rt}$, and

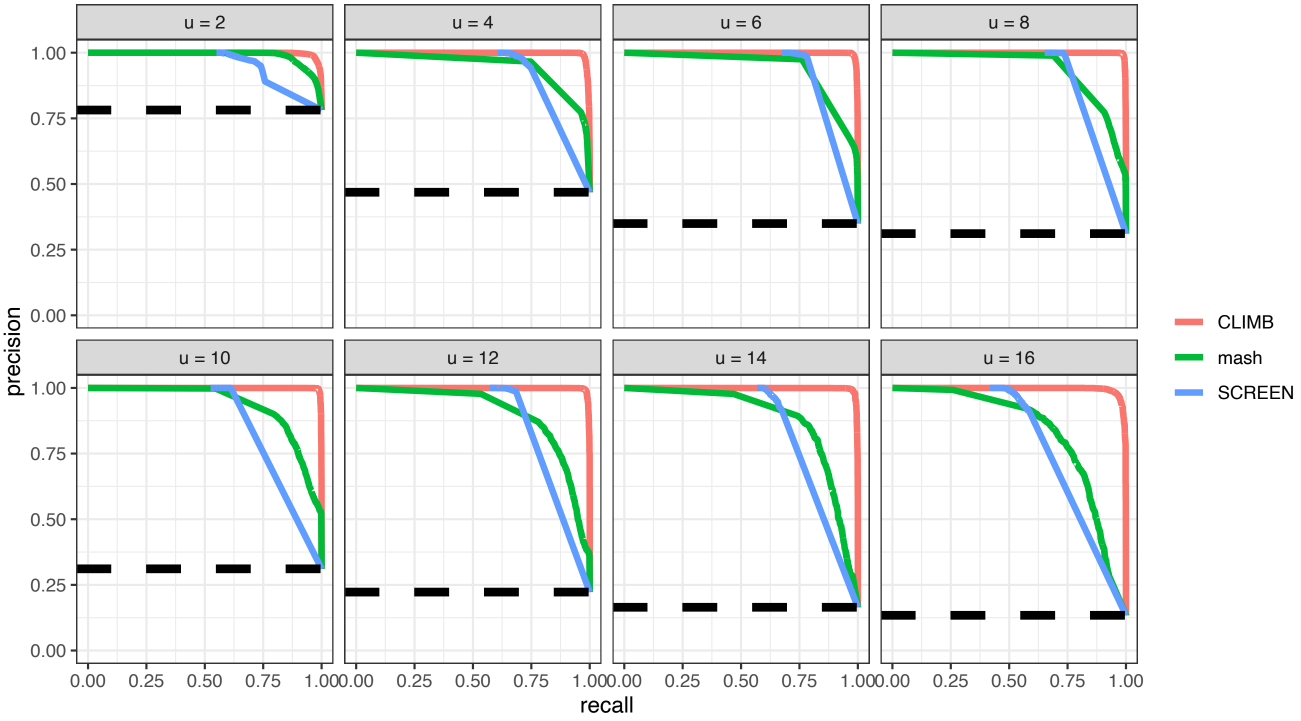


Figure S6: Precision-recall curves analogous to those in Fig. S3, but which use the alternative definitions of FPR and TPR that do not account for missigned effects. Here, precision = $\frac{|\text{significant effects} \cap \text{true effects}|}{|\text{significant effects}|}$ and recall = $\frac{|\text{significant effects} \cap \text{true effects}|}{|\text{true effects}|}$.

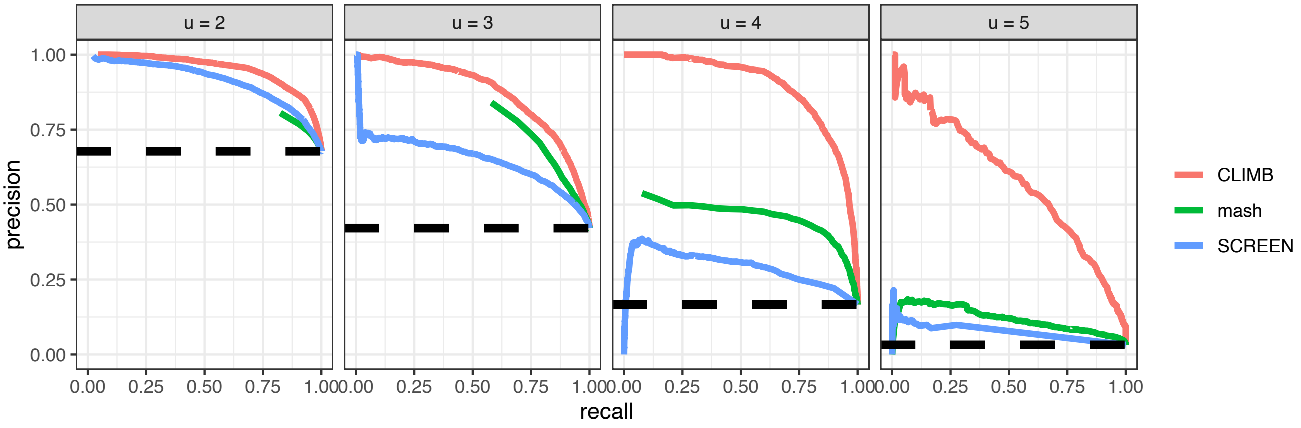


Figure S7: Precision-recall curves analogous to those in Fig. S4, but which use the alternative definitions of FPR and TPR that do not account for missigned effects. Here, precision = $\frac{|\text{significant effects} \cap \text{true effects}|}{|\text{significant effects}|}$ and recall = $\frac{|\text{significant effects} \cap \text{true effects}|}{|\text{true effects}|}$.

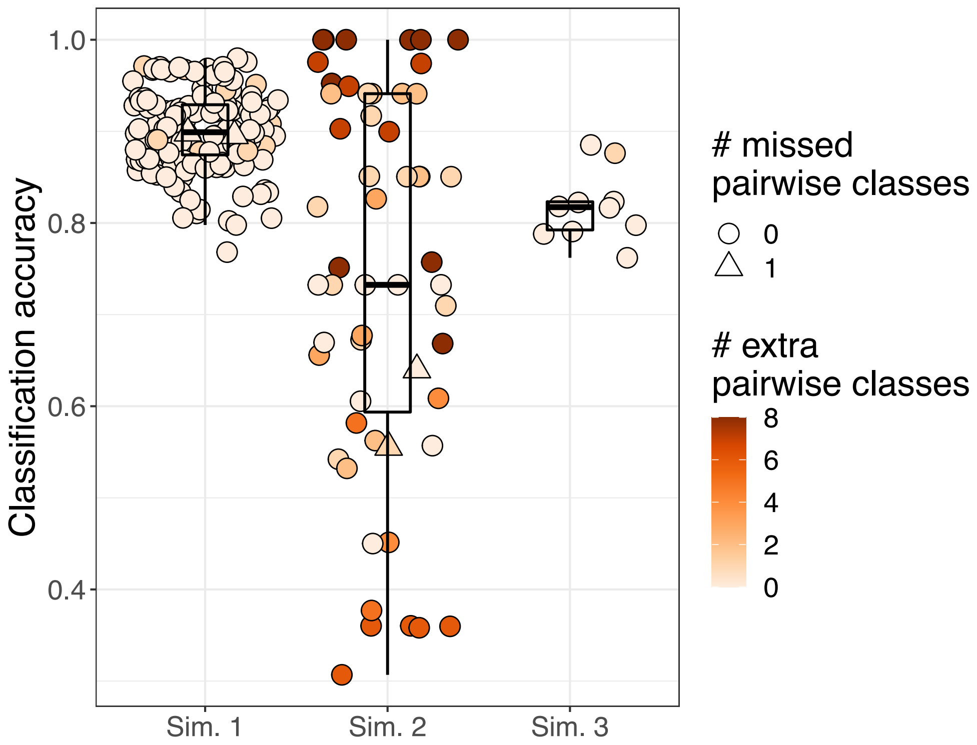


Figure S8: **Classification accuracy of the pairwise fitting step of CLIMB in simulation studies.** Each point in the figure represents the outcome of a pairwise fit. For each pairwise fit, CLIMB estimates which latent classes are present in the model, and assigns each observation a probability of belonging to each class. We assessed CLIMB's classification accuracy by calculating the proportion of times, for a given pairwise fit, the observations' maximum *a posteriori* class estimate matches the true class label. CLIMB was overall more likely to retain extra classes at the pairwise level (# extra pairwise classes) than it was to remove classes from the model that truly belonged (# missed pairwise classes). No more than one class was missed across all pairwise fits in simulation. All triangles (that is, cases where 1 class was erroneously removed from the model) are plotted in the top layer and are thus visible in the figure. Boxplot shows the 25th, 50th, and 75th quantiles.

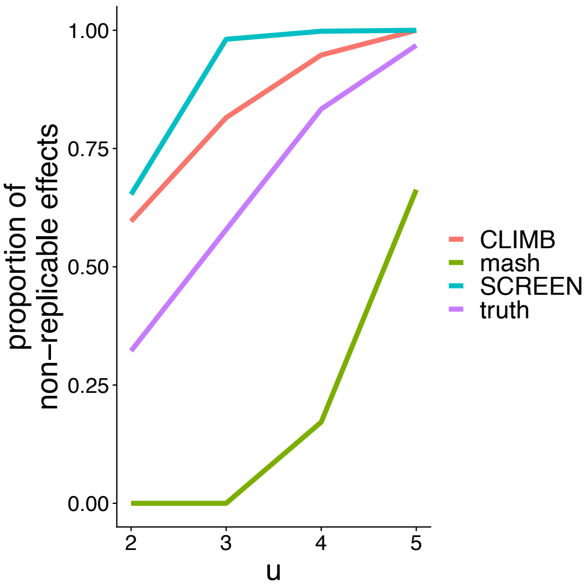


Figure S9: The proportion of effects that are deemed null by CLIMB, mash, and SCREEN at significance level 0.05, as compared to the true proportion of null effects across various thresholds $u$.

3. all off-diagonal covariance terms are equal in magnitude for $h \in \{(-1,-1), (-1,1), (1,-1), (1,1)\}$.

Simulations indicated that, without these constraints, estimation is rendered inconsistent; simultaneous model selection and flexible parameter estimation tended to result in an underestimation of the true number of classes. We maintain that in the pairwise fitting step of CLIMB, accurate estimation of the number of latent classes is the most salient task. While imposing the above parameter constraints results in some estimation bias, we find that the method is still able to consistently estimate the number of latent classes. Thus, we imposed these constraints in all analyses with CLIMB. It is important to note that the $D-$dimensional Bayesian normal mixture model implemented in the last step of the CLIMB methodology does not impose these constraints.

The *Overview of CLIMB* section describes how CLIMB assumes the data model to follow a constrained normal distribution $\phi^c$ (see Equation 2) such that for any class $h$, elements of $\boldsymbol{\mu}_h$ corresponding to non-null dimensions are non-zero. This assumption on the non-null elements of $\boldsymbol{\mu}_h$ can be modified to accomodate a more strict definition of the non-null dimensions of each class. That is, we can assume that a non-null dimension must have a mean whose magnitude is greater than or equal to some positive value, for example, the 95% quantile of a standard normal distribution. Indeed, for all analyses presenting here we assumed the non-null elements of $\boldsymbol{\mu}_h$ to be non-zero.

*Informativity of the prior.* In *An empirical Bayesian model*, we state that the degree of freedom parameters in the Bayesian mixture model, $\boldsymbol{\nu}$ and $\boldsymbol{\kappa}$, can be set approximately equal to $n\hat{\boldsymbol{\alpha}}$, where $\alpha$ is computed from the pairwise fitting results with Equation 6. For each class $h$, these parameters control the concentration of the prior on the covariance around the $\Psi_h^0$ and the concentration of the prior on the mean around $\boldsymbol{\mu}_h^0$, respectively. These terms can therefore be adjusted to tune the informativity of the priors. That is, for smaller values, the priors will become more diffuse, whereas for larger values, the priors will become more concentrated. In all simulations and VISION ChIP-seq analyses, we left $\boldsymbol{\nu}$ and $\boldsymbol{\kappa}$ equal to $n\hat{\boldsymbol{\alpha}}$. In each VISION RNA-seq analysis, we set these hyperparameters equal to 50 for all classes. We chose this path because the clusters were less well-separated, and since $n$ was large for these datasets, setting $\boldsymbol{\kappa} = n\hat{\boldsymbol{\alpha}}$ would possibly place undue confidence in the priors. Deliberately making priors more diffuse is a good strategy for one who is concerned with sensitivity to the prior.

*MCMC implementation details.* Setting up the MCMC for CLIMB requires some user intervention. This is mainly through the selection of threshold $\delta$ (Equation 6), which determines the number $M$ of classes included in the final model. As with any MCMC, the user must also choose the number of iterations for which to run the MCMC and discard initial iterations for a chain-specific burn-in period. These values are reported in Table S3. Parameters for the MCMC were initialized by drawing from the priors. Visual inspection of traceplots for a subset of model parameters was used to assess convergence.

*Inferring similarity between conditions with CLIMB.* Output from CLIMB can be used to determine a relationship among the dimensions in a dataset. To do this, we first compute pairwise distances between dimensions based on the estimated mixture model by extending correlation-based distances[34]. Letting $\hat{C}_m$ be the correlation matrix derived from the estimated covariance matrix $\hat{\Sigma}_m$ for cluster $m$, and $(Y)_{ij}$ be the $(i, j)$th element of matrix $Y$, the distance between dimensions $d$ and $d'$ is

$$\text{dist}(d, d') = \sqrt{1 - \Big[\sum_{m=1}^{M} \pi_m (\hat{C}_m)_{dd'}\Big]^2}. \tag{28}$$

Hierarchical clustering is used on the matrix of pairwise distances found via Equation 28 to extract the association between dimensions.

*Obtaining parsimonious characterization of condition-specificity by merging classes with CLIMB.* To merge classes with CLIMB's output, we compute all pairwise distances between them. Noting that each class is described parametrically by a multivariate normal distribution, we can compute the distance between any two classes as

$$\text{dist}(m, m') = \frac{KL(p||q) + KL(q||p)}{2} \tag{29}$$

where $q$ and $p$ are estimated multivariate normal densities corresponding to classes $m$ and $m'$, and $KL$ is the Kullback-Leibler divergence. Equation 29 can be computed analytically. Then, one can perform hierarchical clustering based on this distance, and cut the tree to produce the desired number of groups. A group of merged classes inherits a new class mean as a weighted average of the mean estimates of the member classes. That is, the mean estimate $\hat{\boldsymbol{\mu}}_G$ of group $G$, comprised of classes $g \in G$, is

$$\hat{\boldsymbol{\mu}}_G = \sum_{g \in G} \frac{\pi_g \hat{\boldsymbol{\mu}}_g}{\sum_{g' \in G} \pi_{g'}}. \tag{30}$$

##### Running SCREEN, mash, and NMF

SCREEN asks the user to select several input settings before running the model. These are

1. `nH`: the maximum number of binary latent class configurations to store in memory
2. `lfdr_method`: the algorithm used to estimate the two-groups model
3. `use_power`: whether or not to weight each dimension by the estimated quality of data in that dimension

In all analyses, we ran SCREEN at the default settings, setting `nH` = 10,000, `lfdr_method` = *znormix*, which is based on the EM algorithm for normal mixtures[69], and `use_power` = True.

Mash asks the user to supply a matrix of standard errors for each observation if available, and also to pre-specify a list of candidate covariance matrices. In all analyses, we used built-in functions of the `mashr` R package to generate the list of candidate covariances. These include

1. canonical covariance matrices (e.g., the identity matrix or matrices that correspond to signals specific to a single dimension) obtained using the `cov_canonical` function
2. data-driven covariances obtained from the subset of the data with the strongest signals
   - strongest signals are selected by first running dimension-specific analyses separately for each dimension, the applying the `get_significant_results` function on the results
   - from these strongest signals, data-driven covariances based principal components analysis with 5 components are found using the `cov_pca` function
   - also from the strongest signals, data-driven covariances based on the Extreme Deconvolution method[70] are computed using the `cov_ed` function

We did not provide mash with standard error estimates as they were not available. All other parameter settings were left at their defaults.

In implementing NMF, we followed the procedure used by Meuleman *et al.*[41]. To avoid sensitivity to random initial conditions, we seeded the NMF algorithm with a singular value decomposition. We examined the performance of the NMF from ranks 5 to 30, and selected the optimal rank as the one to maximize the silhouette coefficient before that same metric began a sharp decline. This NMF analysis completed in 36 hours.

#### Processing of empirical data

For the VISION ChIP-seq analysis, reads were aligned to reference genome mm10 and peaks were called using MACS[71] according to the ENCODE pipeline[72]. The peaks called across various cell types were aligned using BEDTools[73]. CTCF peaks, reported from MACS using $P$-values, were normalized with S3norm across different samples to adjust for variation in both signal and background regions[74]. Normalized values were converted to $Z$-scores by applying the standard normal quantile transformation, then taking the negative such that strong peaks correspond to positive $Z$-scores. For any locus for which a peak was called in at least one cell type but not all others, there are not $P$-values available for the cell types without a called peak. We imputed the $Z$-scores for these absent peaks by imputing a null signal that is randomly simulated from a standard normal distribution. In doing so, this imputation introduces null signals for each cell type at places where we do not have strong evidence for the presence of CTCF and avoids producing ties in the data. The proportion of loci with imputed signals in a given cell type ranged from 0.25% to 33.16%. The final data set is comprised of 10,141 binding sites.

For the VISION RNA-seq data, RNA abundances were quantified with RSEM[75]. For a given lineage, genes whose estimated transcripts per million (TPMs) were equal to zero across all experiments were removed from the analysis. Estimated TPMs were then averaged across two replicates for each cell population. The data were then shifted by an offset to prevent infinities, $\log_2$ transformed, and quantile normalized. Lastly, we applied a location shift to position the data's positive mode over the origin. The corresponding datasets for these lineages respectively contained 21,303, 20,995, and 22,940 genes.

For the DNase-seq data, we downloaded the peak significance data made available via Zenodo (https://doi.org/10.5281/zenodo.3838751, file name dat_FDR01_hg38.RData). We subsetted the samples to the 38 hematopoietic cell populations, and filtered out loci such that a peak was present in two or more samples. As with the ChIP-seq data, we converted the $P$-values to $Z$-scores with a standard normal quantile transformation, and imputed $Z$-scores in the case of absent peaks. A small number of accessible signals were so large as to cause numerical issues for CLIMB. Thus, we applied an upper threshold to the signals, thresholding any value that exceeded the 99.9th quantile of all signals at that quantile.

### Enrichment analyses

Each gene ontology genomic regions enrichment analysis was conducted with an appropriate reference set. For the GREAT analysis described in the analysis of the VISION CTCF ChIP-seq data, the set of all detected CTCF binding sites on chromosome 11, across all cell populations, was used. For the VISION RNA-seq analyses, the set of input genes for each lineage was used as the reference set for each lineage-specific gene ontology enrichment analysis. For the GREAT analyses described in the analysis of the DNase-seq data, the set of all analyzed accessible sites across all autosomes was used as the reference set.

Gene sets obtained by CLIMB and DESeq2 for the differential expression analysis of each of the three lineages examined are presented in Supplementary File 2. To select gene ontology terms displayed in Fig. 4b, we searched for terms that were significantly enriched in at least one gene set (i.e., CLIMB only, DESeq2 only, or their intersect for some lineage), and reported the terms with distinct meanings.

### Analysis of CTCF ChIP-seq on chromosome 7

In order to understand the stability of CLIMB, we complemented our original VISION CTCF ChIP-seq analysis with an examination of CTCF ChIP-seq data on chromosome 7. We prepared these data in the same manner as with chromosome 11. The proportion of loci with imputed signals in a given cell type ranged from 1.00% to 31.18%. The final data set is comprised of 8,983 loci. We identified 3 classes corresponding to constitutive binding behavior: the class of all ones, the class of all ones except for in the CFUE cell population, and the class of all ones except for in the CFUE and MONO cell populations. As in the previous analysis, these 3 classes make up $\sim 36\%$ of all analyzed loci, and the average signal of the classes of constitutive loci is larger than that of the loci that are not constitutive (one-sided $t$-test, $P = 4.52 \times 10^{-4}$). In fact, similar to the analysis of chromosome 11 which contained 15 non-empty classes, we estimated 14 non-empty classes to be in the data from chromosome 7. Indeed, many of the classes are the same as those previously identified (see Supplementary Fig. S29 for a full illustration of non-empty classes). Further, CLIMB again returns a clustering of cell types that more closely resembles the expected relationship among the cell populations when compared against mash and Pearson correlations (Fig. S28). This clustering is quite similar to the one obtained from the analysis of chromosome 11, suggesting that if CTCF binding patterns are similar across chromosomes, CLIMB's inference is fairly robust.

### Proofs

**Proposition 1.** *Assume the estimated pairwise association vectors are equal to or a superset of the true pairwise association vectors. Then the graph-based enumeration and pruning algorithm is* conservative*, in that it results in a collection of all the true latent labels, with the possibility of additional, unsupported classes. That is, letting $\mathcal{H}_{true}$ be the collection of true latent classes and $\mathcal{H}_{alg}$ be the collection of latent classes obtained from the enumeration and pruning algorithm, then $\mathcal{H}_{true} \subseteq \mathcal{H}_{alg}$.*

*Proof.* Assume to the contrary that $\exists h_m \in \mathcal{H}$ but $h_m \notin \mathcal{H}_{alg}$. This implies $\exists (i, j) \in \{1, \ldots, D\}$, $i < j$, such that $(a_i, a_j) \notin h_{ij}$. We obtain an immediate contradiction because $\mathcal{H}_{alg}$ contains all paths $\tilde{h}'$ such that $\tilde{h}' = \{h_m : (a_i, a_j) \in h_{ij} \, \forall i, j \text{ where } i < j\}$. To show that $\mathcal{H}_{alg}$ is not necessarily equal to $\mathcal{H}$, an explicit example of this, when pairwise labels are obtained exactly, is worked out in Fig. 1. □

**Proposition 2.** *The graph-based enumeration and pruning algorithm provides a unique list of latent classes up to permutation of dimension labeling.*

*Proof.* Consider a list of valid paths produced by executing the path enumeration and pruning algorithm, defined by edge set of the form

$$\Big\{\Big[S, (1, a_1)\Big], \Big[(1, a_1), (2, a_2)\Big], \ldots, \Big[(D-1, a_{D-1}), (D, a_D)\Big], \Big[(D, a_D), T\Big]\Big\}$$

as in Equation 5, which produces latent classes of the form $\{a_1, \ldots, a_D\}$. Then, $\forall r, t \in \{1, \ldots, D\}$, $r < t$, we have that $\{a_r, a_t\} \in h_{rt}$. Define a bijection $\varphi : A \to A$ for some set $A$

that forms the symmetric group $S_D$ under composition of permutations. Applying $\varphi$ to each $\{a_1, \ldots, a_D\}$ thus gives a reordering of the latent classes $\{a'_1, \ldots, a'_d\}$, such that $\exists\, h'_{rt} \ni \{a'_r, a'_t\}$. This $h'_{rt}$ is clearly obtained by applying $\varphi$ to the set of ordered dimensions $\{1, 2, \ldots, D\}$ such that the new edge set is of the form

$$\Bigg\{\Big[S,\, (\varphi(1),\, \varphi(a_1))\Big],\, \Big[(\varphi(1),\, \varphi(a_1)),\, (\varphi(2),\, \varphi(a_2))\Big], \ldots,\, \Big[(\varphi(D),\, \varphi(a_D)),\, T\Big]\Bigg\}$$

corresponding to valid latent classes $\{\varphi(a_1), \ldots, \varphi(a_D)\}$. □

**Proposition 3.** *For pairwise labels sampled according to Equation 7, and for some $D$ and $M$, $\hat{\alpha}_m$ is a consistent estimator for the expected posterior mixing proportion of class $m$ that could be obtained from a $D$-dimensional mixture model of the same $M$ classes.*

*Proof.* We will prove the consistensy of $\hat{\alpha}_m$ inductively. For simplicity of notation, we replace previous notation $\mathbf{x}_i^{(rt)}$ with $\mathbf{x}_i^{(p)}$ and $h_m^{(rt)}$ with $h_m^{(p)}$ for pair $p$. First, consider the base case, when $D = 2$. We will show that for $\epsilon > 0$,

$$\lim_{n\to\infty} \lim_{\delta \downarrow 0} \Pr(|\hat{\alpha}_m - \theta_2| > \epsilon) \to 0$$

where $\theta_D$ here is the true expected posterior probability of class membership output from a $D$-dimensional mixture model. That is, for some class class $m \in \{1, \ldots, M\}$, $\theta_D = \frac{1}{n}\sum_{i=1}^n P(\mathbf{x}_i \in h_m)$.

$$\begin{aligned}
&\lim_{n\to\infty} \lim_{\delta \downarrow 0} \Pr(|\hat{\alpha}_m - \theta_2| > \epsilon) \\
&\quad= \lim_{n\to\infty} \lim_{\delta \downarrow 0} \Bigg[\Pr(\hat{\alpha}_m - \theta_2 > \epsilon)\mathbb{1}(\hat{\alpha}_m > \theta_2) + \Pr(\hat{\alpha}_m - \theta_2 < -\epsilon)\mathbb{1}(\hat{\alpha}_m < \theta_2)\Bigg] \\
&\quad= \lim_{n\to\infty} \lim_{\delta \downarrow 0} \Bigg[\mathrm{P}\Bigg(\frac{\sum_{i=1}^n \Big\{\mathbb{1}\Big[\sum_{p=1}^{\binom{D}{2}} \mathbb{1}(\mathbf{x}_i^{(p)} \in h_m^{(p)}) \geq \binom{D}{2} - \delta\Big]\Big\}}{\sum_{m'=1}^M \sum_{i=1}^n \Big\{\mathbb{1}\Big[\sum_{p=1}^{\binom{D}{2}} \mathbb{1}\Big(\mathbf{x}_i^{(p)} \in h_{m'}^{(p)}\Big)\Big] \geq \binom{D}{2} - \delta\Big\}} - \theta_2 > \epsilon\Bigg)\mathbb{1}(\hat{\alpha}_m > \theta_2) \\
&\qquad\quad + \mathrm{P}\Bigg(\frac{\sum_{i=1}^n \Big\{\mathbb{1}\Big[\sum_{p=1}^{\binom{D}{2}} \mathbb{1}(\mathbf{x}_i^{(p)} \in h_m^{(p)}) \geq \binom{D}{2} - \delta\Big]\Big\}}{\sum_{m'=1}^M \sum_{i=1}^n \Big\{\mathbb{1}\Big[\sum_{p=1}^{\binom{D}{2}} \mathbb{1}\Big(\mathbf{x}_i^{(p)} \in h_{m'}^{(p)}\Big)\Big] \geq \binom{D}{2} - \delta\Big\}} - \theta_2 < -\epsilon\Bigg)\mathbb{1}(\hat{\alpha}_m < \theta_2)\Bigg] \\
&\quad= \lim_{n\to\infty} \Bigg[\mathrm{P}\Bigg(\frac{\sum_{i=1}^n \mathbb{1}(\mathbf{x}_i \in h_m)}{\sum_{m'=1}^M \sum_{i=1}^n \mathbb{1}\Big(\mathbf{x}_i \in h_{m'}\Big)} > \theta_2 + \epsilon\Bigg)\mathbb{1}(\hat{\alpha}_m > \theta_2) \\
&\qquad\quad + P\Bigg(\frac{\sum_{i=1}^n \mathbb{1}(\mathbf{x}_i \in h_m)}{\sum_{m'=1}^M \sum_{i=1}^n \mathbb{1}\Big(\mathbf{x}_i \in h_{m'}\Big)} < \theta_2 - \epsilon\Bigg)\mathbb{1}(\hat{\alpha}_m < \theta_2) \qquad \Bigg(\binom{D}{2} = 1\Bigg) \\
&\quad= \lim_{n\to\infty} \Bigg[\mathrm{P}\Bigg(\frac{1}{n}\sum_{i=1}^n \mathbb{1}(\mathbf{x}_i \in h_m) > \theta_2 + \epsilon\Bigg)\mathbb{1}(\hat{\alpha}_m > \theta_2) + P\Bigg(\frac{1}{n}\sum_{i=1}^n \mathbb{1}(\mathbf{x}_i \in h_m) < \theta_2 - \epsilon\Bigg)\mathbb{1}(\hat{\alpha}_m < \theta_2)\Bigg] \\
&\quad= \mathrm{P}\Bigg(\mathbb{E}\Big\{\mathbb{1}(\mathbf{x}_i \in h_m)\Big\} > \theta_2 + \epsilon\Bigg)\mathbb{1}(\hat{\alpha}_m > \theta_2)
\end{aligned}$$

$$+ P\Bigg(\mathbb{E}\Big\{\mathbb{1}(\mathbf{x}_i \in h_m)\Big\} < \theta_2 - \epsilon\Bigg)\mathbb{1}(\hat{\alpha}_m < \theta_2) \qquad \text{(Dominated Convergence Theorem)}$$

$$= \mathrm{P}\Bigg(\mathrm{P}(\mathbf{x}_i \in h_m) > \theta_2 + \epsilon\Bigg)\mathbb{1}(\hat{\alpha}_m > \theta_2) + P\Bigg(\mathrm{P}(\mathbf{x}_i \in h_m) < \theta_2 - \epsilon\Bigg)\mathbb{1}(\hat{\alpha}_m < \theta_2)$$

$$= 0$$

The inductive hypothesis for generic $D$ is

$$\lim_{n\to\infty}\lim_{\delta\downarrow 0}\Pr(|\hat{\alpha}_m - \theta_D| > \epsilon) =$$

$$= \lim_{n\to\infty}\lim_{\delta\downarrow 0}\Bigg[\mathrm{P}\Bigg(\frac{\sum_{i=1}^{n}\Big\{\mathbb{1}\Big[\sum_{p=1}^{\binom{D}{2}}\mathbb{1}(\mathbf{x}_i^{(p)} \in h_m^{(p)}) \geq \binom{D}{2} - \delta\Big]\Big\}}{\sum_{m'=1}^{M}\sum_{i=1}^{n}\Big\{\mathbb{1}\Big[\sum_{p=1}^{\binom{D}{2}}\mathbb{1}\Big(\mathbf{x}_i^{(p)} \in h_{m'}^{(p)}\Big)\Big] \geq \binom{D}{2} - \delta\Big\}} - \theta_D > \epsilon\Bigg)\mathbb{1}(\hat{\alpha}_m > \theta_D)$$

$$+ \mathrm{P}\Bigg(\frac{\sum_{i=1}^{n}\Big\{\mathbb{1}\Big[\sum_{p=1}^{\binom{D}{2}}\mathbb{1}(\mathbf{x}_i^{(p)} \in h_m^{(p)}) \geq \binom{D}{2} - \delta\Big]\Big\}}{\sum_{m'=1}^{M}\sum_{i=1}^{n}\Big\{\mathbb{1}\Big[\sum_{p=1}^{\binom{D}{2}}\mathbb{1}\Big(\mathbf{x}_i^{(p)} \in h_{m'}^{(p)}\Big)\Big] \geq \binom{D}{2} - \delta\Big\}} - \theta_D < -\epsilon\Bigg)\mathbb{1}(\hat{\alpha}_m < \theta_D)\Bigg]$$

$$\to 0$$

and we want to show that, for $D+1$, we still have

$$\lim_{n\to\infty}\lim_{\delta\downarrow 0}\Pr(|\hat{\alpha}_m - \theta_{D+1}| > \epsilon) =$$

$$= \lim_{n\to\infty}\lim_{\delta\downarrow 0}\Bigg[\mathrm{P}\Bigg(\frac{\sum_{i=1}^{n}\Big\{\mathbb{1}\Big[\sum_{p=1}^{\binom{D+1}{2}}\mathbb{1}(\mathbf{x}_i^{(p)} \in h_m^{(p)}) \geq \binom{D+1}{2} - \delta\Big]\Big\}}{\sum_{m'=1}^{3M}\sum_{i=1}^{n}\Big\{\mathbb{1}\Big[\sum_{p=1}^{\binom{D+1}{2}}\mathbb{1}\Big(\mathbf{x}_i^{(p)} \in h_{m'}^{(p)}\Big)\Big] \geq \binom{D+1}{2} - \delta\Big\}} - \theta_{D+1} > \epsilon\Bigg)\mathbb{1}(\hat{\alpha}_m > \theta_{D+1})$$

$$+ \mathrm{P}\Bigg(\frac{\sum_{i=1}^{n}\Big\{\mathbb{1}\Big[\sum_{p=1}^{\binom{D+1}{2}}\mathbb{1}(\mathbf{x}_i^{(p)} \in h_m^{(p)}) \geq \binom{D+1}{2} - \delta\Big]\Big\}}{\sum_{m'=1}^{3M}\sum_{i=1}^{n}\Big\{\mathbb{1}\Big[\sum_{p=1}^{\binom{D+1}{2}}\mathbb{1}\Big(\mathbf{x}_i^{(p)} \in h_{m'}^{(p)}\Big)\Big] \geq \binom{D+1}{2} - \delta\Big\}} - \theta_{D+1} < -\epsilon\Bigg)\mathbb{1}(\hat{\alpha}_m < \theta_{D+1})\Bigg]$$

$$\to 0$$

where $3M$ comes from the fact the if there are $M$ classes in $D$ dimensions, there can be no more than $3M$ classes in $D+1$ dimensions. Simply letting $D^\star = D+1$ and $M^\star = 3M$, we get

$$\lim_{n\to\infty}\lim_{\delta\downarrow 0}\Pr(|\hat{\alpha}_m - \theta_{D^\star}| > \epsilon) =$$

$$
\begin{aligned}
&= \lim_{n\to\infty}\lim_{\delta\downarrow 0}\Bigg[\mathrm{P}\Bigg(\frac{\sum_{i=1}^{n}\Bigg\{\mathbb{1}\Bigg[\sum_{p=1}^{\binom{D^\star}{2}}\mathbb{1}(\mathbf{x}_i^{(p)}\in h_m^{(p)})\geq\binom{D^\star}{2}-\delta\Bigg]\Bigg\}}{\sum_{m'=1}^{M^\star}\sum_{i=1}^{n}\Bigg\{\mathbb{1}\Bigg[\sum_{p=1}^{\binom{D^\star}{2}}\mathbb{1}\Big(\mathbf{x}_i^{(p)}\in h_{m'}^{(p)}\Big)\Bigg]\geq\binom{D^\star}{2}-\delta\Bigg\}}-\theta_{D^\star}>\epsilon\Bigg)\mathbb{1}(\hat{\alpha}_m>\theta_{D^\star})\\
&\qquad+\mathrm{P}\Bigg(\frac{\sum_{i=1}^{n}\Bigg\{\mathbb{1}\Bigg[\sum_{p=1}^{\binom{D^\star}{2}}\mathbb{1}(\mathbf{x}_i^{(p)}\in h_m^{(p)})\geq\binom{D^\star}{2}-\delta\Bigg]\Bigg\}}{\sum_{m'=1}^{M^\star}\sum_{i=1}^{n}\Bigg\{\mathbb{1}\Bigg[\sum_{p=1}^{\binom{D^\star}{2}}\mathbb{1}\Big(\mathbf{x}_i^{(p)}\in h_{m'}^{(p)}\Big)\Bigg]\geq\binom{D^\star}{2}-\delta\Bigg\}}-\theta_{D^\star}<-\epsilon\Bigg)\mathbb{1}(\hat{\alpha}_m<\theta_{D^\star})\Bigg]\\
&\to 0 \qquad\qquad \text{(inductive hypothesis)}
\end{aligned}
$$

□

### Supplementary figures and tables

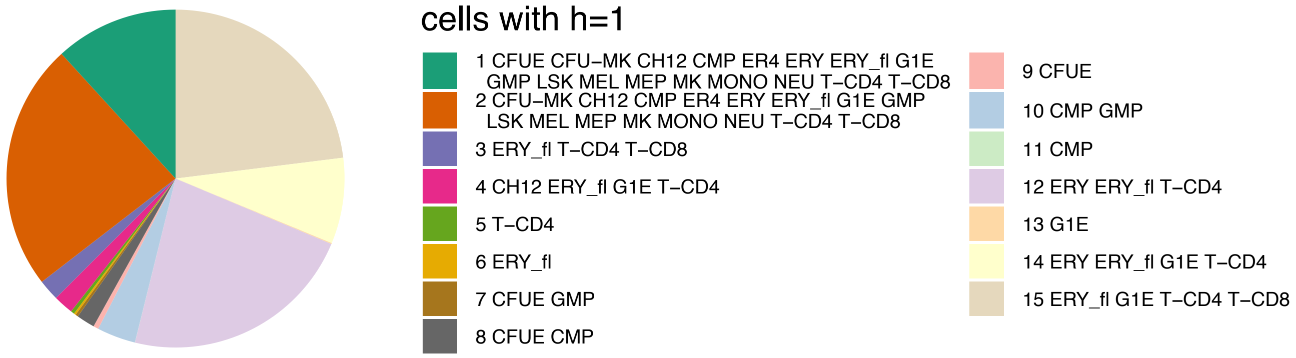


Figure S10: Class sizes in the CTCF ChIP-seq data analysis of chromosome 11. The final model estimated 15 non-empty classes, each described by association vectors with binary elements. The legend shows which cell types in each class were assigned a 1, indicating presence of CTCF.

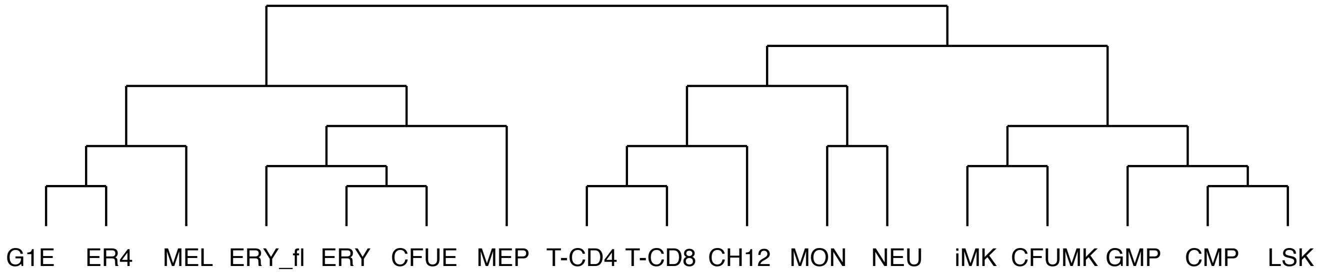


Figure S11: Assumed ground truth hierarchical tree among 17 hematopoietic cell populations. A hierarchical tree capturing the ground truth is needed for quantification of a method's performance with respect to cell population clustering. This clustering is largely derived from the hierarchical clustering of ATAC-seq data on mouse blood cell populations from previous work[24].

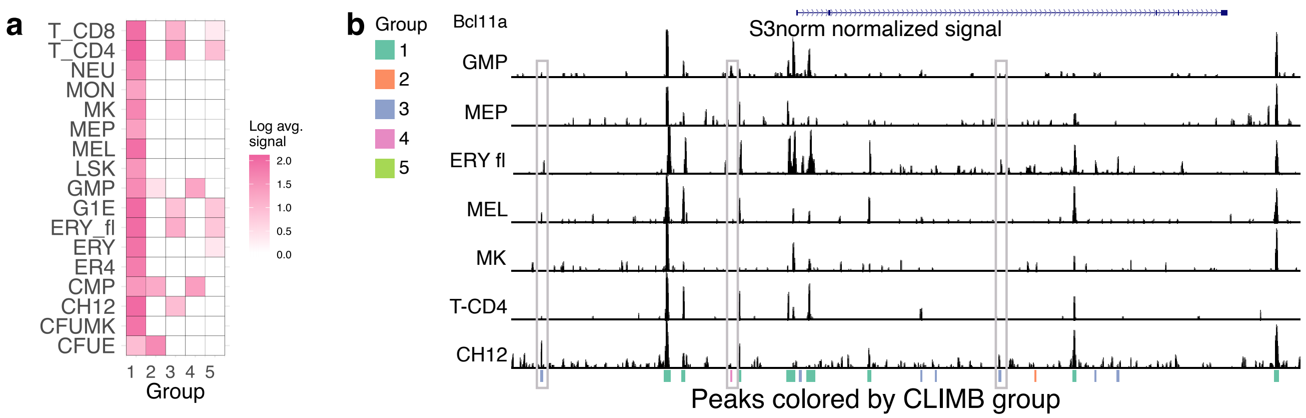


Figure S12: **CLIMB uncovers interrelationships among hematopoietic cell populations based on CTCF binding patterns. a,** For ease of visualization, CLIMB facilitates merging similar clusters. After merging 15 classes into 5 parent groups, the new group means show the most salient CTCF binding patterns in the data. **b,** Genome browser track shot of 7 hematopoietic cell types around the murine gene *Bcl11a* (chr11:24,014,641–24,190,463). Normalized peaks are marked by group membership in **a**; exemplifying non-constitutively bound sites are circumscribed by gray rectangles.

| Data | $M$ after step 2 | $M$ | $\delta$ | burn-in | total iterations |
|---|---|---|---|---|---|
| Simulation 1 | 128 | 48 | 5 | 2,500 | 10,000 |
| Simulation 2 | 57 | 38 | 14 | 2,500 | 10,000 |
| Simulation 3 | 34 | 34 | 5 | 2,500 | 10,000 |
| VISION–ChIP chr. 7 | 388 | 35 | 25 | 20,000 | 40,000 |
| VISION–ChIP chr. 11 | 548 | 37 | 28 | 10,000 | 25,000 |
| VISION–RNA: | | | | | |
| Erythroid lineage | 34 | 34 | 10 | 5,000 | 15,000 |
| Megakaryocytic lineage | 33 | 33 | 10 | 5,000 | 15,000 |
| Myeloid lineage | 34 | 34 | 10 | 5,000 | 15,000 |
| ENCODE DNase-seq: | | | | | |
| chr. 1 | 56 | 33 | 150 | 8,000 | 20,000 |
| chr. 2 | 80 | 22 | 150 | 8,000 | 20,000 |
| chr. 3 | 42 | 14 | 150 | 8,000 | 20,000 |
| chr. 4 | 31 | 12 | 108 | 8,000 | 20,000 |
| chr. 5 | 62 | 15 | 79 | 8,000 | 20,000 |
| chr. 6 | 230 | 22 | 114 | 8,000 | 20,000 |
| chr. 7 | 41 | 16 | 114 | 8,000 | 20,000 |
| chr. 8 | 134 | 13 | 79 | 8,000 | 20,000 |
| chr. 9 | 131 | 14 | 144 | 8,000 | 20,000 |
| chr. 10 | 215 | 14 | 114 | 8,000 | 20,000 |
| chr. 11 | 38 | 19 | 150 | 8,000 | 20,000 |
| chr. 12 | 38 | 14 | 150 | 8,000 | 20,000 |
| chr. 13 | 14 | 14 | 703 | 8,000 | 20,000 |
| chr. 14 | 77 | 13 | 110 | 8,000 | 20,000 |
| chr. 15 | 51 | 13 | 110 | 8,000 | 20,000 |
| chr. 16 | 56 | 23 | 143 | 8,000 | 20,000 |
| chr. 17 | 128 | 29 | 145 | 8,000 | 20,000 |
| chr. 18 | 26 | 7 | 41 | 8,000 | 20,000 |
| chr. 19 | 173 | 23 | 150 | 8,000 | 20,000 |
| chr. 20 | 46 | 12 | 86 | 8,000 | 20,000 |
| chr. 21 | 157 | 7 | 39 | 8,000 | 20,000 |
| chr. 22 | 140 | 11 | 78 | 8,000 | 20,000 |

Table S3: MCMC details for all CLIMB analyses. $M$ after step 2 refers to the number of candidate latent classes after pruning by concordance with the non-adjacent pairs (step 2) and before pruning by mixing weights (step 3). $M$ is the number of classes included in the final model as determined by threshold $\delta$. Burn-in periods were determined based on each chain's time to enter its stationary distribution.

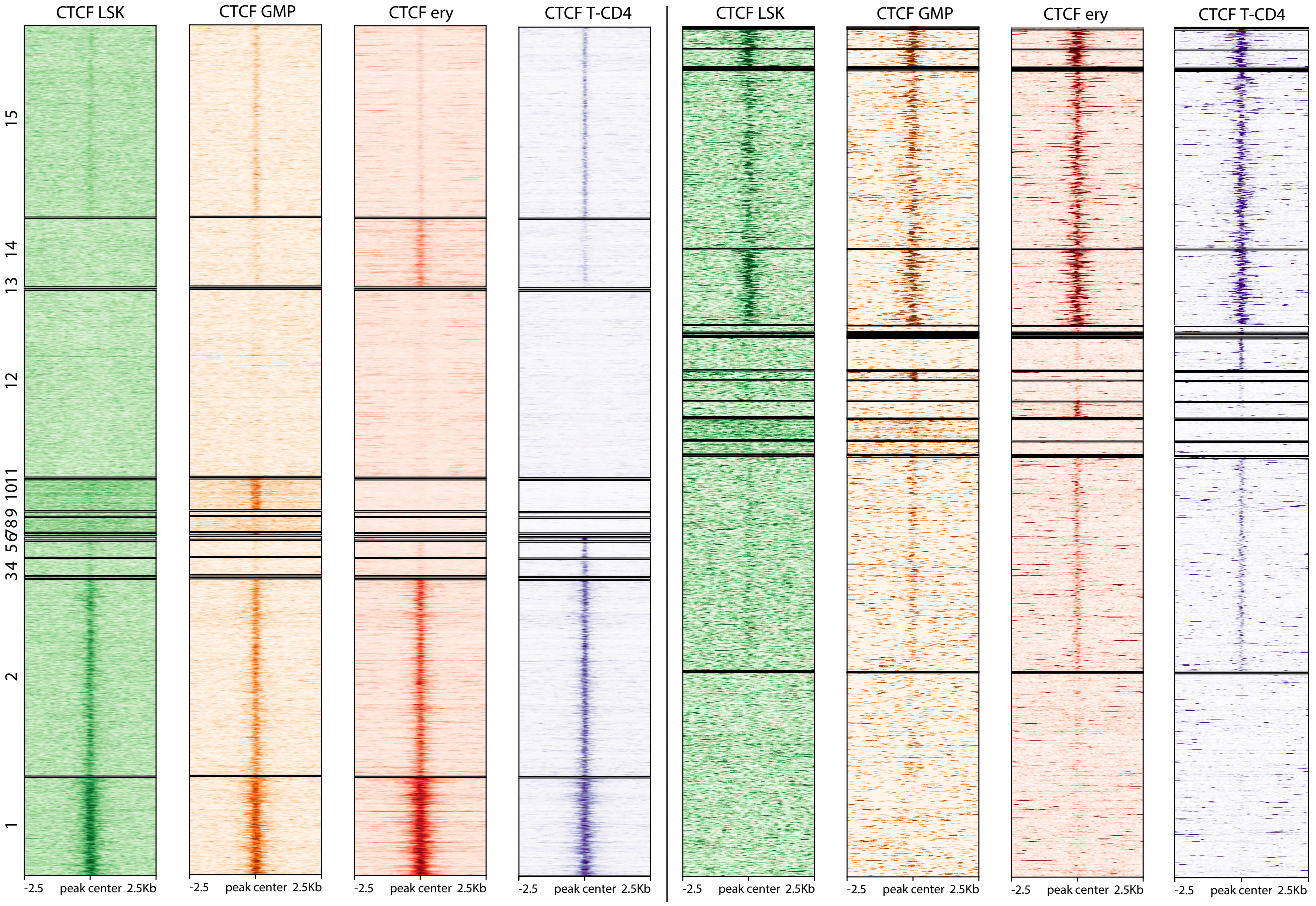


Figure S13: CTCF ChIP-seq signals for binding sites on chromosome 11 in LSK, GMP, ERY, and T-CD4 cells. The same data is clustered by CLIMB (left) and mash (right).

| cluster num. | 1 | 2 | 3 | 4 | 5 | 6 | 7 | 8 | 9 | 10 | 11 |
|---|---|---|---|---|---|---|---|---|---|---|---|
| $\pi$ | 4.46E-02 | 2.37E-03 | 6.19E-03 | 1.38E-02 | 3.42E-03 | 6.44E-07 | 4.12E-07 | 9.00E-03 | 2.74E-02 | 1.77E-02 | 5.34E-03 |
| cluster num. | 12 | 13 | 14 | 15 | 16 | 17 | 18 | 19 | 20 | 21 | 22 |
| $\pi$ | 6.12E-07 | 1.41E-06 | 7.04E-07 | 8.10E-03 | 7.22E-07 | 4.63E-03 | 6.46E-03 | 3.90E-03 | 8.69E-03 | 6.59E-03 | 2.33E-02 |
| cluster num. | 23 | 24 | 25 | 26 | 27 | 28 | 29 | 30 | 31 | 32 | 33 |
| $\pi$ | 2.16E-07 | 4.76E-03 | 1.95E-02 | 4.78E-07 | 3.23E-03 | 8.91E-02 | 3.96E-02 | 3.90E-02 | 3.72E-02 | 2.24E-02 | 1.97E-02 |
| cluster num. | 34 | 35 | 36 | 37 | 38 | 39 | 40 | 41 | 42 | 43 | |
| $\pi$ | 3.48E-02 | 3.74E-02 | 3.83E-02 | 2.17E-02 | 7.74E-03 | 5.30E-02 | 1.39E-01 | 4.05E-02 | 1.46E-02 | 1.47E-01 | |

Table S4: Mixing weights for each cluster used in simulation 1.

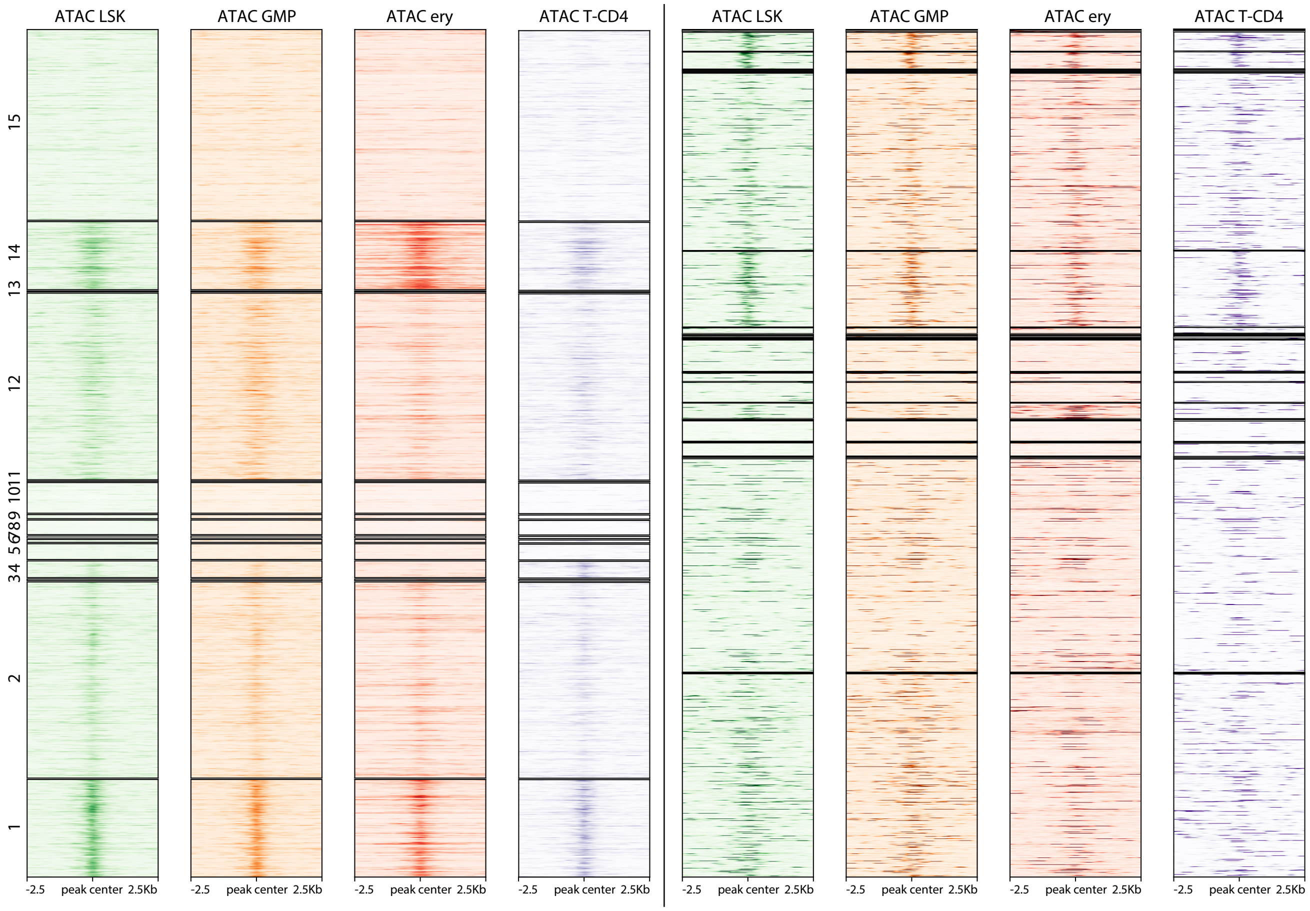


Figure S14: ATAC-seq signals for CTCF binding sites on chromosome 11 in LSK, GMP, ERY, and T-CD4 cells. The same data is clustered by CLIMB (left) and mash (right).

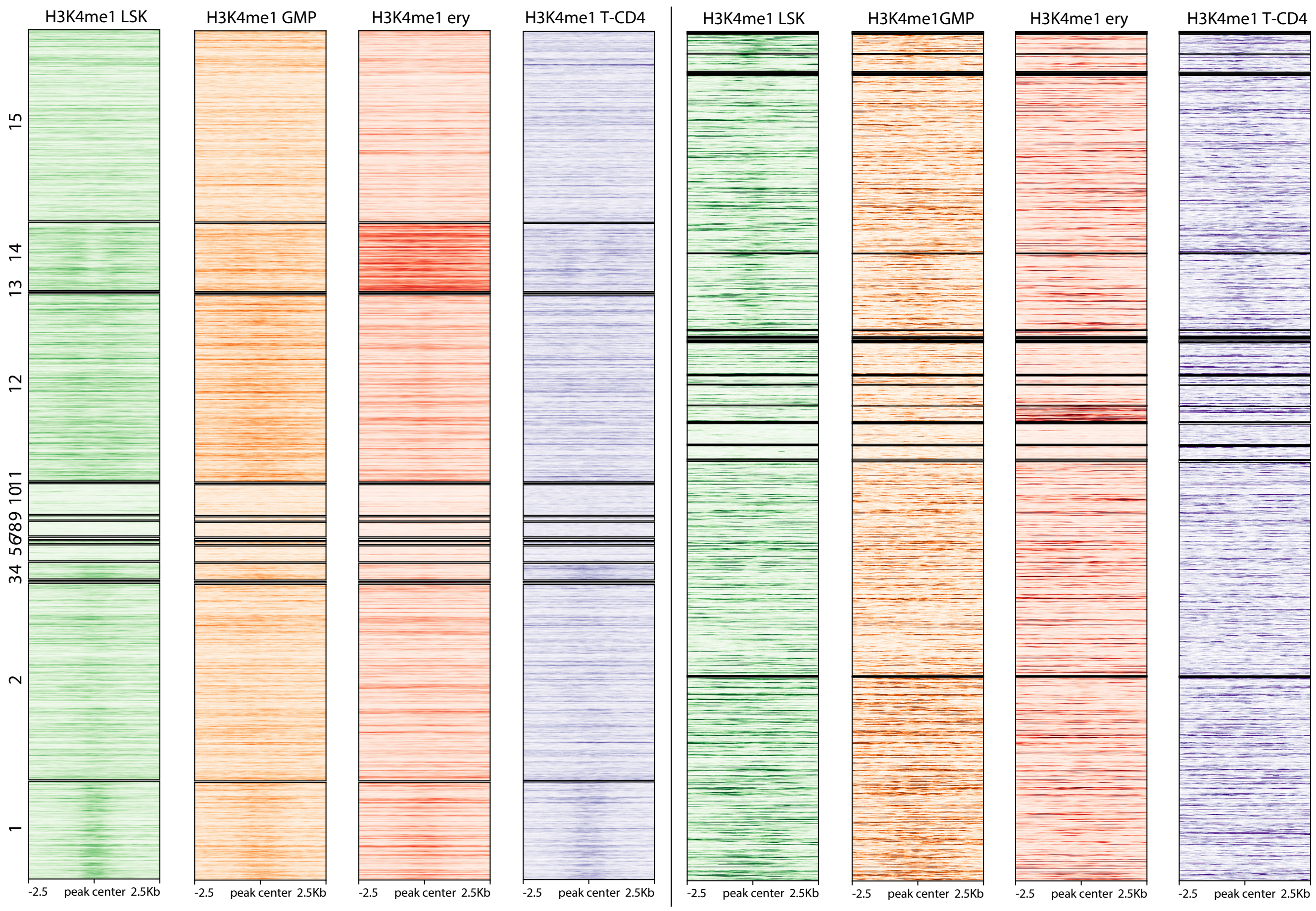


Figure S15: H3K4me1 ChIP-seq signals for CTCF binding sites on chromosome 11 in LSK, GMP, ERY, and T-CD4 cells. The same data is clustered by CLIMB (left) and mash (right).

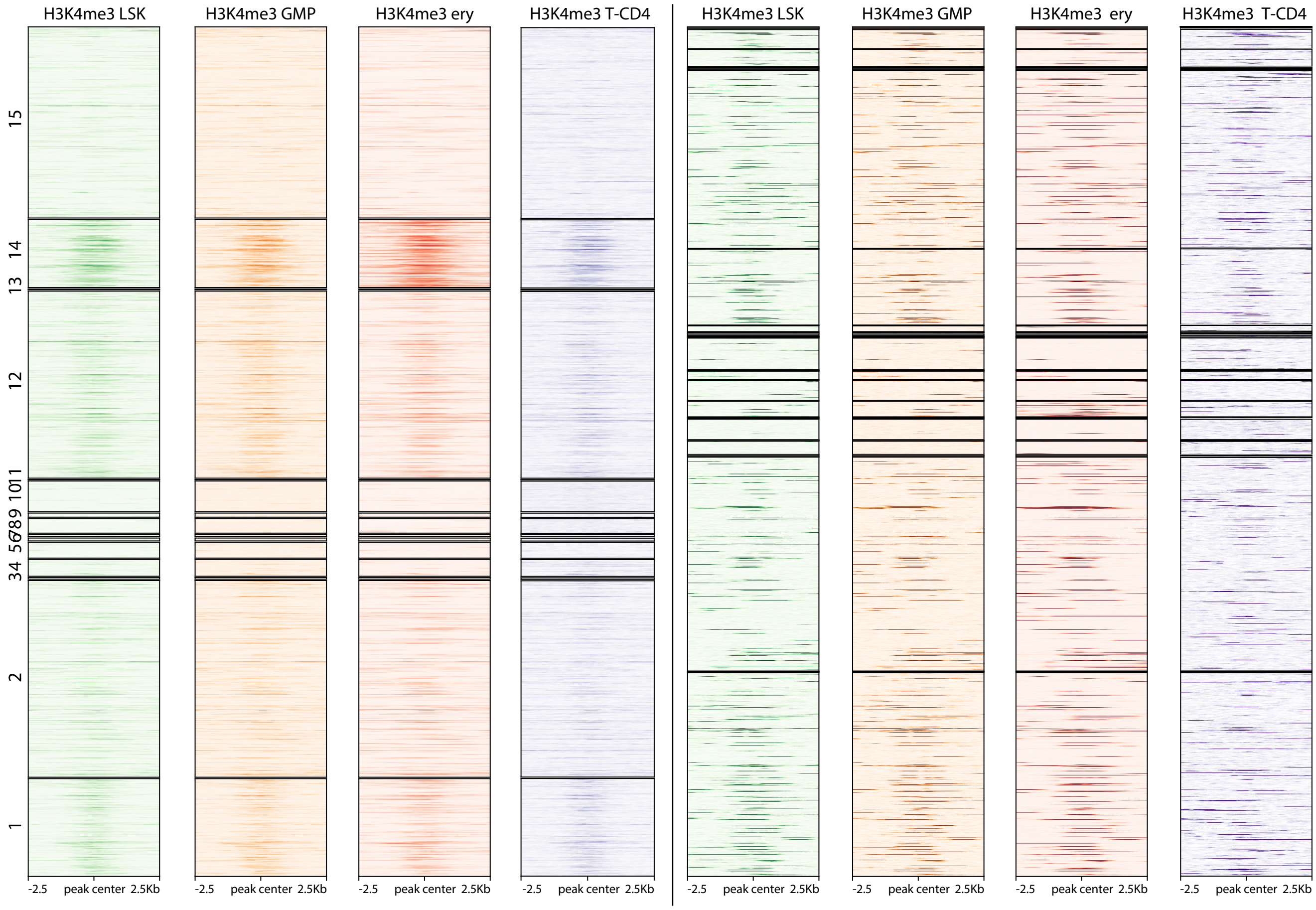


Figure S16: H3K4me3 ChIP-seq signals for CTCF binding sites on chromosome 11 in LSK, GMP, ERY, and T-CD4 cells. The same data is clustered by CLIMB (left) and mash (right).

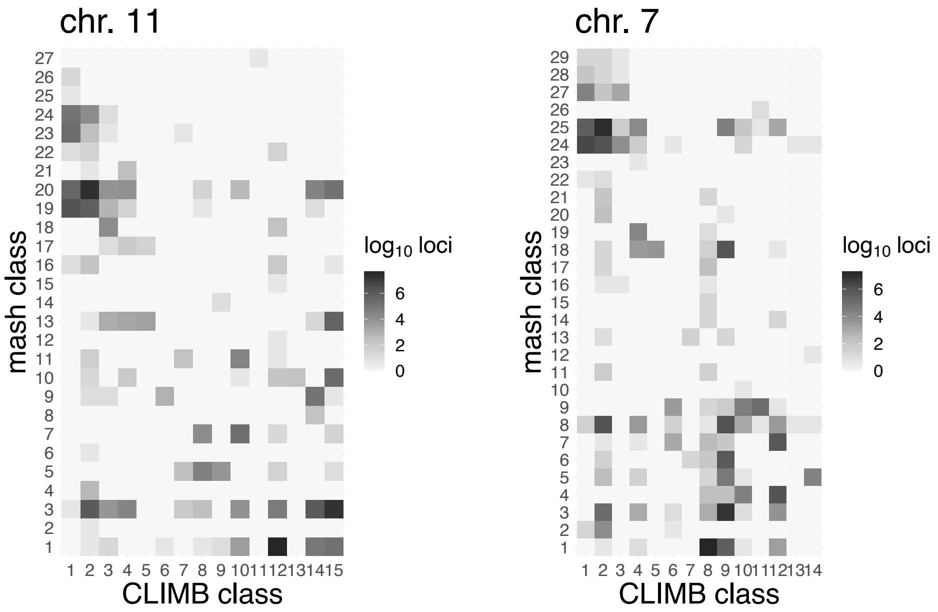


Figure S17: Comparison of classifications of CTCF binding sites using CLIMB and mash for the chromosome 11 and chromosome 7 CTCF ChIP-seq data.

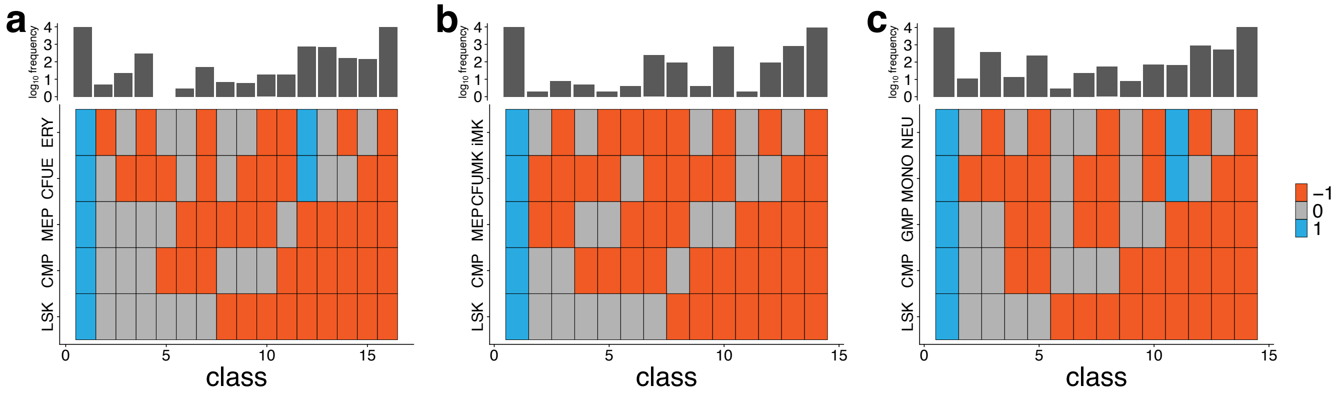


Figure S18: **Class labels and sizes in each of the VISION RNA-seq data analyses.** The gene expression classes obtained from CLIMB for the **a,** erythroid, **b,** megakaryocytic, and **c,** myeloid lineages are represented as a series of blocks in each column, with one block for the genes in that expession class for each cell types. The blocks are colored by the expression category, with -1 for genes that are lowly expressed or off, 0 for moderately expressed genes, and 1 for highly expressed genes. All genes were assigned to a class based on their maximum *a posteriori* class estimate. The log (base 10) of the number of genes found in each class is plotted above the matrix of blocks.

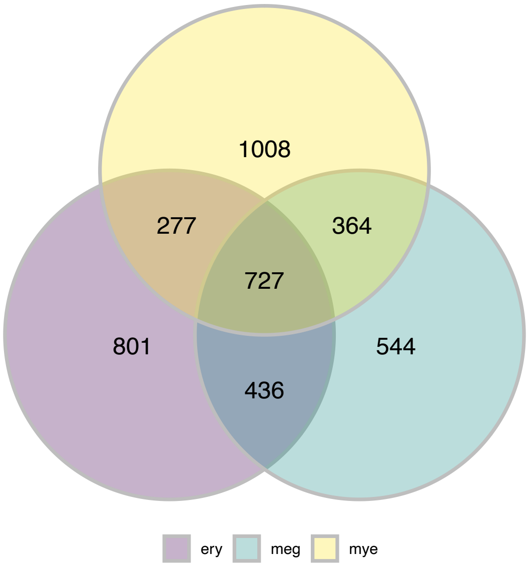


Figure S19: **Overlap of CLIMB's differentially expressed genes across 3 lineages.** Venn diagram of the overlap in genes that CLIMB identified as differentially expressed across three studied hematopoietic lineages.

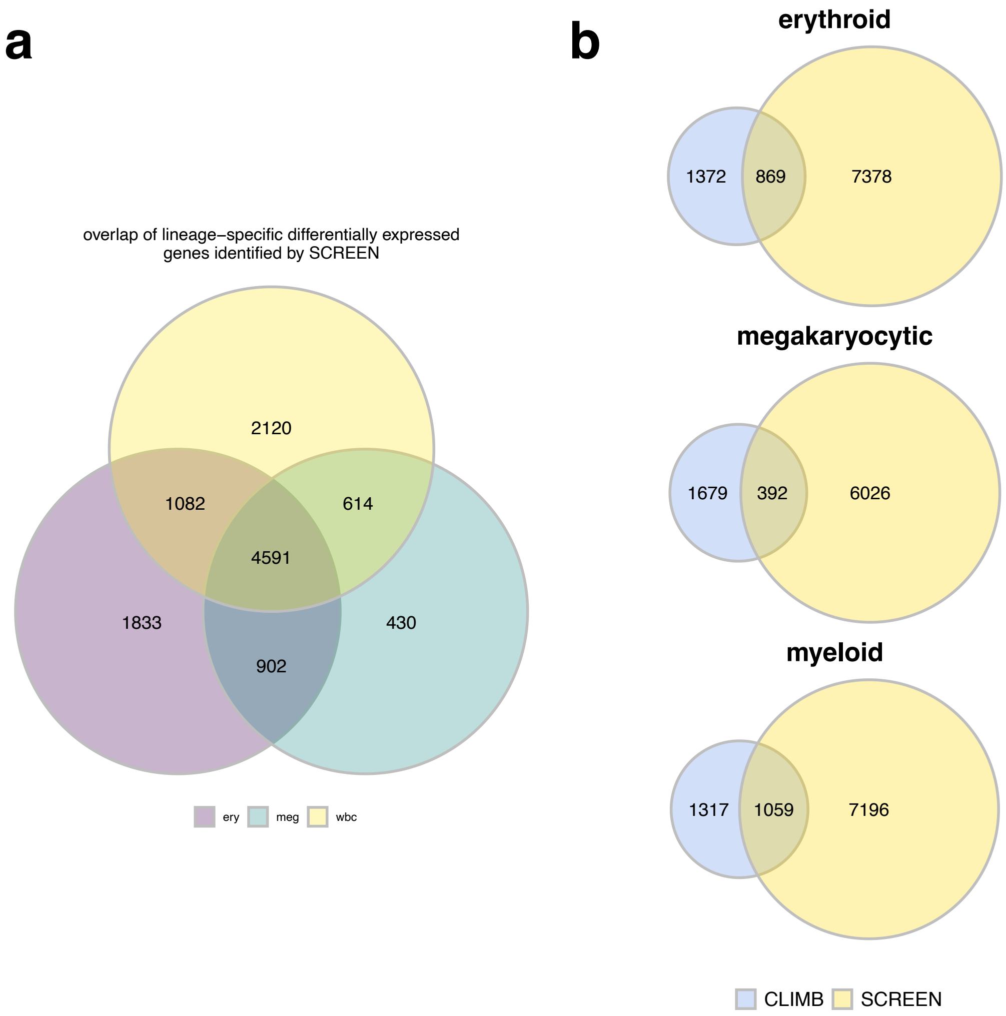


Figure S20: **SCREEN's differentially expressed genes across 3 lineages differ from CLIMB's.** **a,** Venn diagram of the overlap in genes that SCREEN identified as differentially expressed across three studied hematopoietic lineages. **b,** Overlap of differentially expressed genes found by CLIMB and SCREEN across all three hematopoietic lineages.

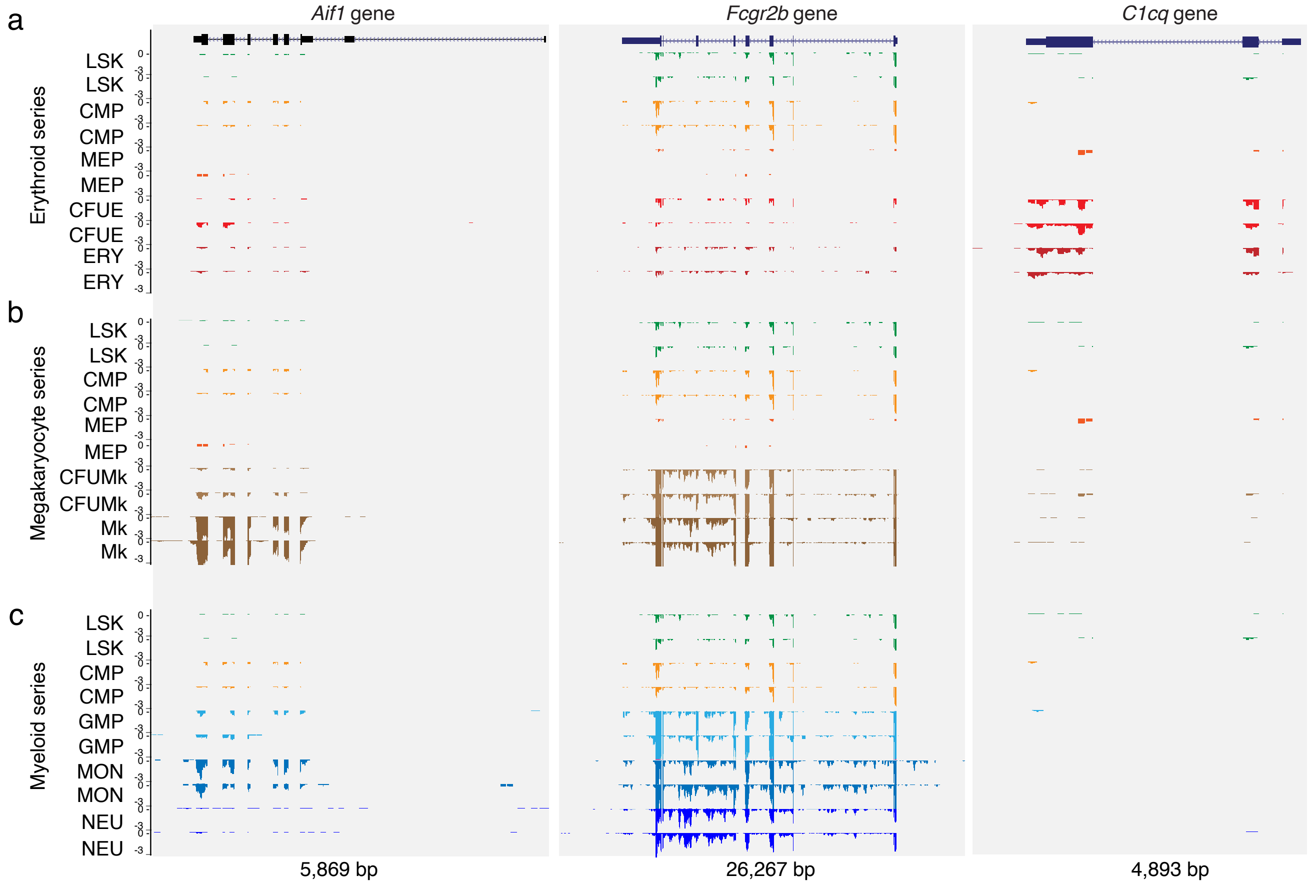


Figure S21: **Lineage-specific differential gene expression.** Some myeloid-specific gene ontology (GO) terms were significantly enriched in the differentially expressed gene sets for the non-myeloid lineages (Fig. 4b). To more closely examine this result, we collected the genes whose expression patterns led to the myeloid-associated GO term enrichments in non-myeloid lineages. Three genes driving these GO term enrichments were *Aif1*, *C1qc*, and *Fcgr2b*. We visualized the RNA-seq patterns at these representative loci *Aif1* (chr17: 35,169, 740 – 35,177,254), *C1cq* (chr4: 136,888,989 – 136, 893,881) and *Fcgr2b* (chr1: 170,954,239 – 170,980,505) for the **a,** erythroid series, **b,** megakaryocyte series, and **c,** myeloid series. The RNA-seq data are shown only for the minus strand (with respect to the reference genome) because the genes are oriented right to left in the mouse genome assembly (mm10). In the conventions for this display of stranded RNA-seq data, negative values indicate the density of reads mapping to the minus strand. Indeed, the raw data support that these genes are differentially expressed across megakaryocyte and erythroid lineages, though they are generally expressed at much lower levels than for the myeloid lineage.

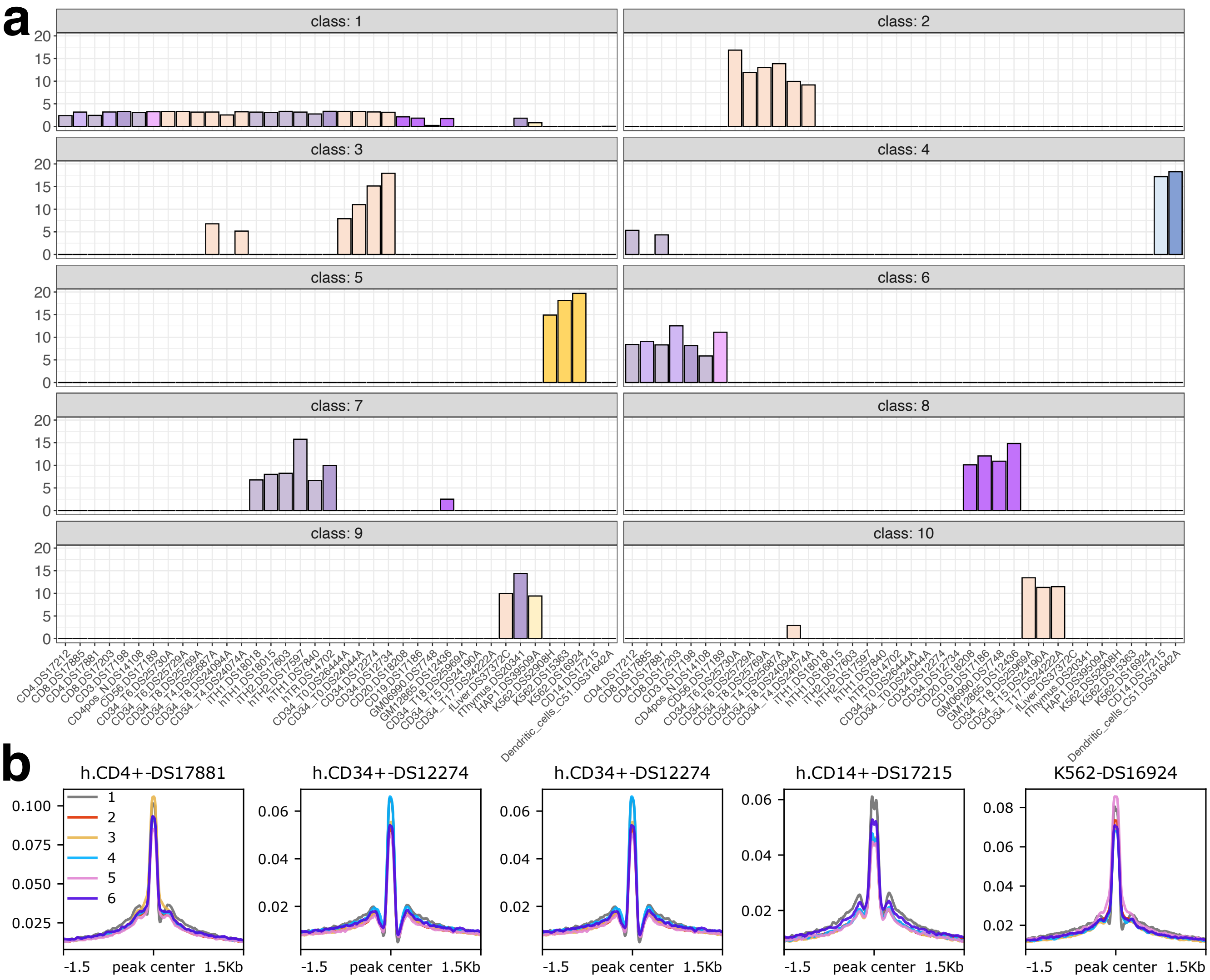


Figure S22: **NMF applied to ENCODE DNase-seq data in 38 cell populations. a,** Signals for each of the 10 classes described by the coefficient matrix estimated via NMF. Each class captures different patterns of chromatin accessibility across cell populations. For example, class 4 captures loci accessible in CD4+ T cells, classical monocytes, and dendritic cells, while class 5 describes loci that are accessible in K562 cells. Samples are ordered based on their similarity according to NMF output. **b,** Transcription factor footprint signatures ($\log_2$[observed/expected cleavage] for DNase-seq) for classes 1–6 in a subset of the examined cell populations. Within a given cell population, footprint signatures are very similar across all classes.

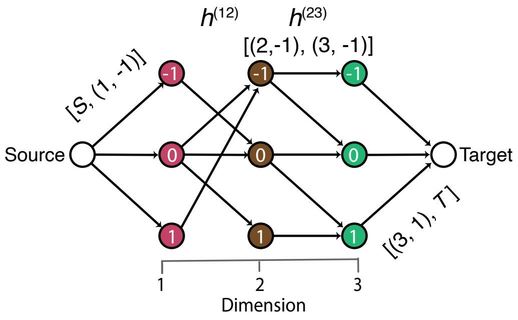


Figure S23: Graphical illustration of the construction of full association vectors from pairwise vectors for the toy example in Fig. 1.

| class num. \ dim. | 1 | 2 | 3 | 4 | 5 | 6 | 7 | 8 | 9 | 10 | 11 | 12 | 13 | 14 | 15 | 16 | 17 | 18 |
|---|---|---|---|---|---|---|---|---|---|---|---|---|---|---|---|---|---|---|
| 1 | 13.41 | 13.94 | 5.57 | 11.02 | 8.90 | 5.87 | 13.13 | 17.05 | 16.49 | 15.45 | 6.56 | 17.42 | 7.03 | 14.28 | 6.27 | 8.41 | 7.63 | 7.55 |
| 2 | 13.75 | 13.47 | 5.41 | 10.64 | 8.86 | 5.57 | 12.68 | 17.43 | 16.63 | 15.54 | 6.25 | 17.12 | 6.64 | 14.01 | 0 | 8.25 | 7.42 | 7.27 |
| 3 | 14.36 | 14.27 | 4.66 | 10.78 | 8.17 | 5.45 | 13.25 | 18.96 | 18.45 | 16.52 | 6.11 | 19.80 | 0 | 14.51 | 5.73 | 7.50 | 8.05 | 7.08 |
| 4 | 9.76 | 11.06 | 3.99 | 7.29 | 4.38 | 3.95 | 8.67 | 15.83 | 14.28 | 12.88 | 4.89 | 17.10 | 0 | 9.29 | 0 | 4.38 | 6.04 | 4.14 |
| 5 | 13.73 | 13.49 | 5.37 | 10.61 | 8.68 | 0 | 12.89 | 17.03 | 16.45 | 15.48 | 6.28 | 17.24 | 6.72 | 14.06 | 6.01 | 8.03 | 7.45 | 7.33 |
| 6 | 14.10 | 14.71 | 5.68 | 11.01 | 9.08 | 0 | 16.16 | 19.64 | 18.02 | 16.72 | 6.66 | 19.88 | 7.31 | 16.18 | 0 | 8.72 | 9.68 | 8.62 |
| 7 | 13.83 | 14.21 | 5.73 | 11.26 | 9.18 | 0 | 16.81 | 19.55 | 17.66 | 17.08 | 6.97 | 20.35 | 0 | 15.88 | 7.95 | 9.25 | 8.96 | 8.21 |
| 8 | 11.39 | 15.61 | 4.99 | 11.83 | 0 | 6.37 | 12.64 | 19.37 | 17.58 | 15.19 | 7.21 | 20.71 | 6.23 | 12.03 | 5.79 | 7.67 | 8.32 | 7.24 |
| 9 | 13.67 | 13.86 | 0 | 10.95 | 8.81 | 5.81 | 12.56 | 16.38 | 16.29 | 15.33 | 6.49 | 17.52 | 6.94 | 13.93 | 6.14 | 8.20 | 7.55 | 7.47 |
| 10 | 15.68 | 13.18 | 0 | 9.41 | 5.98 | 3.33 | 9.07 | 11.85 | 16.65 | 16.39 | 4.73 | 22.28 | 4.07 | 10.15 | 3.52 | 0 | 5.93 | 5.17 |
| 11 | 13.90 | 13.31 | 0 | 10.68 | 8.53 | 5.70 | 12.06 | 16.07 | 15.93 | 15.42 | 6.16 | 17.07 | 6.61 | 13.32 | 0 | 7.93 | 7.34 | 7.21 |
| 12 | 13.67 | 14.32 | 0 | 14.26 | 10.45 | 6.73 | 15.72 | 19.26 | 18.10 | 17.46 | 7.74 | 20.05 | 8.21 | 16.59 | 0 | 0 | 9.15 | 9.62 |
| 13 | 13.65 | 14.29 | 0 | 14.14 | 10.44 | 6.61 | 15.71 | 19.56 | 18.08 | 17.09 | 7.61 | 20.01 | 0 | 16.34 | 8.25 | 10.29 | 9.86 | 8.50 |
| 14 | 13.91 | 14.55 | 0 | 14.31 | 10.22 | 6.75 | 15.66 | 19.42 | 18.19 | 16.89 | 7.93 | 20.13 | 0 | 16.21 | 0 | 11.15 | 9.51 | 9.59 |
| 15 | 9.98 | 10.15 | 0 | 7.60 | 4.10 | 5.40 | 6.67 | 10.81 | 12.82 | 12.28 | 4.23 | 19.78 | 0 | 7.71 | 0 | 0 | 6.10 | 3.22 |
| 16 | 13.68 | 14.02 | 0 | 13.57 | 10.10 | 6.37 | 15.32 | 19.13 | 17.79 | 16.91 | 0 | 18.95 | 9.04 | 16.04 | 6.99 | 10.19 | 9.54 | 8.47 |
| 17 | 14.08 | 13.57 | 0 | 10.87 | 8.96 | 0 | 12.66 | 17.26 | 16.54 | 16.13 | 6.31 | 17.19 | 6.66 | 14.16 | 6.05 | 8.00 | 7.20 | 7.27 |
| 18 | 16.39 | 13.60 | 0 | 10.72 | 8.13 | 0 | 11.05 | 16.83 | 16.54 | 17.90 | 5.41 | 17.31 | 6.07 | 13.07 | 0 | 7.07 | 6.00 | 6.65 |
| 19 | 14.81 | 13.74 | 0 | 10.14 | 8.26 | 0 | 12.06 | 17.03 | 16.92 | 15.58 | 5.60 | 17.75 | 0 | 13.94 | 5.76 | 7.46 | 7.25 | 6.78 |
| 20 | 15.51 | 12.42 | 0 | 9.69 | 7.01 | 0 | 9.74 | 15.51 | 15.60 | 16.19 | 5.62 | 17.17 | 0 | 12.17 | 0 | 6.48 | 6.24 | 5.89 |
| 21 | 13.19 | 11.50 | 0 | 8.66 | 6.35 | 0 | 9.93 | 14.12 | 14.75 | 14.22 | 6.01 | 17.52 | 0 | 11.79 | 0 | 0 | 6.57 | 6.66 |
| 22 | 11.25 | 6.92 | 0 | 4.38 | 5.89 | 0 | 4.81 | 8.77 | 10.22 | 12.53 | 1.66 | 13.73 | 0 | 6.97 | 0 | 0 | 2.67 | 0 |
| 23 | 13.26 | 13.45 | 0 | 13.13 | 10.06 | 0 | 14.23 | 20.81 | 17.77 | 17.25 | 0 | 18.46 | 7.95 | 17.21 | 0 | 9.56 | 8.42 | 7.92 |
| 24 | 14.99 | 10.87 | 0 | 8.52 | 5.98 | 0 | 9.06 | 15.39 | 14.93 | 15.53 | 0 | 14.93 | 0 | 12.81 | 0 | 6.07 | 4.26 | 5.38 |
| 25 | 12.15 | 9.15 | 0 | 5.31 | 4.33 | 0 | 6.47 | 10 | 12.41 | 13.07 | 0 | 17.26 | 0 | 7.39 | 0 | 0 | 4.71 | 5.82 |
| 26 | 13.54 | 13.95 | 0 | 13.97 | 0 | 6.70 | 17.40 | 21.50 | 19.03 | 17.43 | 7.84 | 19.17 | 8.17 | 17.07 | 7.17 | 9.76 | 9.65 | 8.56 |
| 27 | 13.79 | 14.68 | 0 | 10.72 | 0 | 5.26 | 10.61 | 15.60 | 16.62 | 13.93 | 6.29 | 21.38 | 0 | 12.04 | 4.67 | 7.21 | 8.27 | 7.09 |
| 28 | 6.95 | 4.77 | 0 | 2.97 | 0 | 0 | 2.72 | 6.02 | 7.70 | 8.16 | 1.62 | 12.76 | 0 | 3.52 | 0 | 0 | 2.68 | 0 |
| 29 | 3.20 | 4.92 | 0 | 0 | 0 | 0 | 0 | 6.34 | 4.27 | 3.37 | 0 | 12.27 | 0 | 0 | 0 | 0 | 4.32 | 0 |
| 30 | 4.33 | 6.22 | 0 | 0 | 0 | 0 | 0 | 0 | 2.73 | 3.23 | 0 | 6.39 | 0 | 0 | 0 | 0 | 0 | 0 |
| 31 | 3.27 | 0 | 0 | 0 | 0 | 0 | 0 | 6.58 | 4.65 | 3.88 | 0 | 6.40 | 0 | 0 | 0 | 0 | 0 | 0 |
| 32 | 3.98 | 0 | 0 | 0 | 0 | 0 | 0 | 0 | 4.01 | 4.90 | 0 | 8.07 | 0 | 0 | 0 | 0 | 5.68 | 0 |
| 33 | 7.54 | 0 | 0 | 0 | 0 | 0 | 0 | 0 | 6.75 | 5.58 | 0 | 8.88 | 0 | 0 | 0 | 0 | 0 | 0 |
| 34 | 7.12 | 0 | 0 | 0 | 0 | 0 | 0 | 0 | 0 | 7.55 | 0 | 7.21 | 0 | 0 | 0 | 0 | 0 | 0 |
| 35 | 6.25 | 0 | 0 | 0 | 0 | 0 | 0 | 0 | 0 | 0 | 0 | 8.72 | 0 | 0 | 0 | 0 | 0 | 0 |
| 36 | 5.91 | 0 | 0 | 0 | 0 | 0 | 0 | 0 | 0 | 0 | 0 | 0 | 0 | 0 | 0 | 0 | 0 | 0 |
| 37 | 0 | 0 | 0 | 0 | 0 | 5.01 | 0 | 0 | 0 | 0 | 0 | 0 | 0 | 0 | 0 | 0 | 0 | 0 |
| 38 | 0 | 0 | 0 | 0 | 0 | 0 | 0 | 0 | 7.82 | 8.51 | 0 | 9.07 | 0 | 0 | 0 | 0 | 0 | 0 |
| 39 | 0 | 0 | 0 | 0 | 0 | 0 | 0 | 0 | 4.73 | 0 | 0 | 6.11 | 0 | 0 | 0 | 0 | 0 | 0 |
| 40 | 0 | 0 | 0 | 0 | 0 | 0 | 0 | 0 | 0 | 3.07 | 0 | 5.86 | 0 | 0 | 0 | 0 | 0 | 0 |
| 41 | 0 | 0 | 0 | 0 | 0 | 0 | 0 | 0 | 0 | 0 | 0 | 10.47 | 0 | 0 | 0 | 0 | 5.20 | 0 |
| 42 | 0 | 0 | 0 | 0 | 0 | 0 | 0 | 0 | 0 | 0 | 0 | 15.15 | 0 | 0 | 0 | 0 | 0 | 0 |
| 43 | 0 | 0 | 0 | 0 | 0 | 0 | 0 | 0 | 0 | 0 | 0 | 0 | 0 | 0 | 0 | 0 | 5.51 | 0 |

Table S5: Mean vectors used for each cluster in simulation 1.

| class num. \ dim. | 1 | 2 | 3 | 4 | 5 | 6 | 7 | 8 | 9 | 10 | 11 | 12 | 13 | 14 | 15 | 16 | 17 | 18 |
|---|---|---|---|---|---|---|---|---|---|---|---|---|---|---|---|---|---|---|
| 1 | 86.14 | 102.03 | 8.99 | 55.22 | 27.63 | 9.40 | 88.46 | 171.81 | 155.10 | 125.12 | 14.16 | 154.87 | 11.25 | 91.43 | 7.80 | 19.68 | 35.63 | 13.41 |
| 2 | 82.62 | 82.97 | 6.87 | 43.26 | 26.88 | 7.79 | 71.21 | 159.69 | 139.34 | 109.76 | 11.28 | 134.98 | 8.91 | 75.92 | 1 | 16.94 | 29.70 | 11.95 |
| 3 | 84.21 | 73.62 | 9.71 | 36.54 | 22.27 | 8.06 | 64.75 | 144.97 | 123.36 | 96.42 | 11.50 | 138.75 | 1 | 71.04 | 6.00 | 15.97 | 30.30 | 13.02 |
| 4 | 100.60 | 76.69 | 11.91 | 32.29 | 19.68 | 10.36 | 60.23 | 151.95 | 132.23 | 93.69 | 12.62 | 176.36 | 1 | 74.07 | 1 | 12.84 | 27.34 | 14.01 |
| 5 | 83.43 | 85.67 | 7.91 | 45.87 | 25.68 | 1 | 75.75 | 153.81 | 138.65 | 114.60 | 12.26 | 142.19 | 9.98 | 80.97 | 7.13 | 17.81 | 31.26 | 11.76 |
| 6 | 91.96 | 119.80 | 5.88 | 32.63 | 19.63 | 1 | 62.32 | 132.34 | 142.50 | 90.38 | 8.27 | 106.01 | 7.05 | 78.77 | 1 | 11.95 | 19.61 | 8.79 |
| 7 | 82.79 | 82.37 | 7.96 | 46.86 | 24.89 | 1 | 73.59 | 151.39 | 126.46 | 118.88 | 13.58 | 130.97 | 1 | 79.20 | 8.01 | 19.03 | 30.74 | 12.00 |
| 8 | 79.53 | 94.77 | 12.29 | 47.40 | 1 | 11.29 | 84.83 | 150.70 | 120.32 | 126.56 | 15.92 | 156.64 | 11.42 | 79.15 | 9.50 | 19.44 | 32.86 | 13.71 |
| 9 | 84.04 | 95.87 | 1 | 49.26 | 25.74 | 8.76 | 79.18 | 150.99 | 140.75 | 115.76 | 13.25 | 149.44 | 10.39 | 84.58 | 7.24 | 18.24 | 33.93 | 12.76 |
| 10 | 61.44 | 52.69 | 1 | 16.84 | 25.75 | 10.71 | 50.73 | 73.40 | 66.68 | 75.85 | 12.73 | 106.37 | 10.16 | 54.82 | 7.27 | 1 | 28.54 | 14.28 |
| 11 | 85.33 | 82.84 | 1 | 43.11 | 23.75 | 7.65 | 69.31 | 137.69 | 126.49 | 106.37 | 11.93 | 141.01 | 9.93 | 79.97 | 1 | 17.13 | 30.10 | 12.86 |
| 12 | 71.77 | 87.22 | 1 | 48.61 | 26.59 | 9.67 | 74.68 | 144.17 | 128.24 | 102.40 | 13.67 | 138.80 | 12.02 | 78.57 | 1 | 1 | 30.58 | 14.26 |
| 13 | 79.43 | 94.20 | 1 | 48.24 | 27.06 | 9.53 | 80.88 | 157.11 | 136.86 | 109.78 | 13.43 | 140.99 | 1 | 85.50 | 8.15 | 19.00 | 32.55 | 12.79 |
| 14 | 83.47 | 97.27 | 1 | 52.15 | 26.07 | 9.48 | 84.15 | 167.18 | 135.79 | 119.70 | 14.28 | 142.58 | 1 | 83.68 | 1 | 19.96 | 30.71 | 13.13 |
| 15 | 56.30 | 50.15 | 1 | 16.03 | 19.51 | 1.57 | 32.01 | 63.65 | 42.99 | 67.42 | 9.83 | 110.40 | 1 | 47.43 | 1 | 1 | 15.46 | 10.54 |
| 16 | 70.60 | 69.80 | 1 | 33.03 | 19.66 | 6.92 | 60.55 | 117.78 | 111.83 | 101.34 | 1 | 140.73 | 8.81 | 67.63 | 7.22 | 16.34 | 26.52 | 10.39 |
| 17 | 77.57 | 81.80 | 1 | 43.00 | 24.27 | 1 | 70.22 | 143.41 | 130.14 | 113.10 | 11.80 | 134.24 | 9.48 | 76.07 | 6.55 | 16.15 | 30.67 | 11.93 |
| 18 | 67.07 | 60.84 | 1 | 26.77 | 22.28 | 1 | 51.32 | 115.23 | 84.95 | 78.48 | 12.00 | 118.60 | 6.27 | 59.58 | 1 | 11.96 | 30.33 | 12.69 |
| 19 | 78.27 | 77.16 | 1 | 34.97 | 20.68 | 1 | 66.77 | 124.46 | 117.87 | 98.09 | 13.29 | 132.79 | 1 | 69.24 | 6.16 | 15.42 | 29.24 | 13.95 |
| 20 | 61.55 | 53.64 | 1 | 27.36 | 23.32 | 1 | 53.39 | 108.14 | 73.76 | 71.32 | 9.49 | 80.86 | 1 | 58.45 | 1 | 12.43 | 24.03 | 13.26 |
| 21 | 63.88 | 61.95 | 1 | 35.67 | 24.00 | 1 | 55.69 | 115.05 | 97.60 | 81.28 | 8.19 | 110.31 | 1 | 67.68 | 1 | 1 | 25.24 | 9.11 |
| 22 | 52.59 | 27.41 | 1 | 15.76 | 7.32 | 1 | 23.23 | 54.24 | 39.81 | 46.00 | 6.24 | 59.83 | 1 | 36.75 | 1 | 1 | 12.87 | 1 |
| 23 | 56.51 | 58.71 | 1 | 31.04 | 18.74 | 1 | 53.71 | 83.76 | 90.97 | 77.88 | 1 | 73.61 | 8.86 | 51.04 | 1 | 12.38 | 30.58 | 6.43 |
| 24 | 62.64 | 36.83 | 1 | 17.24 | 18.67 | 1 | 27.86 | 74.57 | 61.36 | 65.30 | 1 | 59.36 | 1 | 28.08 | 1 | 7.96 | 15.17 | 9.24 |
| 25 | 60.75 | 47.10 | 1 | 20.27 | 18.11 | 1 | 33.73 | 67.60 | 51.29 | 66.94 | 1 | 77.07 | 1 | 45.69 | 1 | 1 | 19.05 | 4.78 |
| 26 | 71.30 | 80.95 | 1 | 44.81 | 1 | 8.22 | 77.33 | 154.82 | 118.33 | 87.39 | 11.78 | 123.39 | 10.50 | 81.11 | 7.50 | 15.91 | 34.66 | 10.80 |
| 27 | 64.18 | 63.03 | 1 | 23.62 | 1 | 7.92 | 64.74 | 119.51 | 96.42 | 86.99 | 12.07 | 139.27 | 1 | 56.62 | 9.26 | 11.32 | 25.53 | 10.95 |
| 28 | 42.48 | 25.22 | 1 | 10.48 | 1 | 1 | 12.66 | 39.49 | 35.74 | 39.42 | 6.63 | 80.89 | 1 | 16.62 | 1 | 1 | 12.68 | 1 |
| 29 | 22.68 | 17.49 | 1 | 1 | 1 | 1 | 1 | 26.21 | 21.53 | 19.10 | 1 | 55.34 | 1 | 1 | 1 | 1 | 11.37 | 1 |
| 30 | 23.49 | 11.78 | 1 | 1 | 1 | 1 | 1 | 1 | 13.34 | 16.78 | 1 | 31.06 | 1 | 1 | 1 | 1 | 1 | 1 |
| 31 | 18.16 | 1 | 1 | 1 | 1 | 1 | 1 | 13.53 | 23.42 | 22.08 | 1 | 31.15 | 1 | 1 | 1 | 1 | 1 | 1 |
| 32 | 15.68 | 1 | 1 | 1 | 1 | 1 | 1 | 1 | 19.04 | 18.53 | 1 | 28.96 | 1 | 1 | 1 | 1 | 2.83 | 1 |
| 33 | 20.07 | 1 | 1 | 1 | 1 | 1 | 1 | 1 | 23.73 | 28.80 | 1 | 38.57 | 1 | 1 | 1 | 1 | 1 | 1 |
| 34 | 21.40 | 1 | 1 | 1 | 1 | 1 | 1 | 1 | 1 | 31.42 | 1 | 44.20 | 1 | 1 | 1 | 1 | 1 | 1 |
| 35 | 29.86 | 1 | 1 | 1 | 1 | 1 | 1 | 1 | 1 | 1 | 1 | 44.17 | 1 | 1 | 1 | 1 | 1 | 1 |
| 36 | 4.16 | 1 | 1 | 1 | 1 | 1 | 1 | 1 | 1 | 1 | 1 | 1 | 1 | 1 | 1 | 1 | 1 | 1 |
| 37 | 1 | 1 | 1 | 1 | 1 | 0.63 | 1 | 1 | 1 | 1 | 1 | 1 | 1 | 1 | 1 | 1 | 1 | 1 |
| 38 | 1 | 1 | 1 | 1 | 1 | 1 | 1 | 1 | 20.83 | 17.56 | 1 | 36.87 | 1 | 1 | 1 | 1 | 1 | 1 |
| 39 | 1 | 1 | 1 | 1 | 1 | 1 | 1 | 1 | 20.84 | 1 | 1 | 34.08 | 1 | 1 | 1 | 1 | 1 | 1 |
| 40 | 1 | 1 | 1 | 1 | 1 | 1 | 1 | 1 | 1 | 16.79 | 1 | 19.03 | 1 | 1 | 1 | 1 | 1 | 1 |
| 41 | 1 | 1 | 1 | 1 | 1 | 1 | 1 | 1 | 1 | 1 | 1 | 66.84 | 1 | 1 | 1 | 1 | 18.69 | 1 |
| 42 | 1 | 1 | 1 | 1 | 1 | 1 | 1 | 1 | 1 | 1 | 1 | 130.41 | 1 | 1 | 1 | 1 | 1 | 1 |
| 43 | 1 | 1 | 1 | 1 | 1 | 1 | 1 | 1 | 1 | 1 | 1 | 1 | 1 | 1 | 1 | 1 | 10.22 | 1 |

Table S6: Variance vectors used for each cluster in simulation 1.

| cluster num. | 1 | 2 | 3 | 4 | 5 | 6 | 7 | 8 | 9 | 10 |
|---|---|---|---|---|---|---|---|---|---|---|
| $\boldsymbol{\pi}$ | 3.14E-02 | 2.84E-04 | 5.61E-02 | 2.02E-02 | 5.93E-04 | 1.98E-02 | 1.01E-01 | 7.33E-06 | 1.99E-06 | 1.26E-01 |

| cluster num. | 11 | 12 | 13 | 14 | 15 | 16 | 17 | 18 | 19 | 20 |
|---|---|---|---|---|---|---|---|---|---|---|
| $\boldsymbol{\pi}$ | 3.67E-06 | 3.92E-06 | 1.00E-04 | 2.71E-06 | 2.67E-04 | 6.47E-02 | 6.03E-04 | 2.23E-02 | 7.46E-04 | 5.41E-06 |

| cluster num. | 21 | 22 | 23 | 24 | 25 | 26 | 27 | 28 | 29 | 30 |
|---|---|---|---|---|---|---|---|---|---|---|
| $\boldsymbol{\pi}$ | 6.28E-02 | 9.49E-03 | 1.15E-02 | 2.30E-02 | 1.64E-06 | 2.58E-02 | 1.00E-04 | 3.25E-02 | 1.61E-02 | 1.16E-04 |

| cluster num. | 31 | 32 | 33 | 34 | 35 | 36 | 37 | 38 | 39 | 40 |
|---|---|---|---|---|---|---|---|---|---|---|
| $\boldsymbol{\pi}$ | 8.13E-02 | 5.87E-05 | 1.42E-01 | 1.50E-02 | 1.78E-02 | 9.24E-02 | 7.67E-03 | 3.43E-04 | 7.06E-03 | 1.11E-02 |

Table S7: Mixing weights for each cluster used in simulation 2.

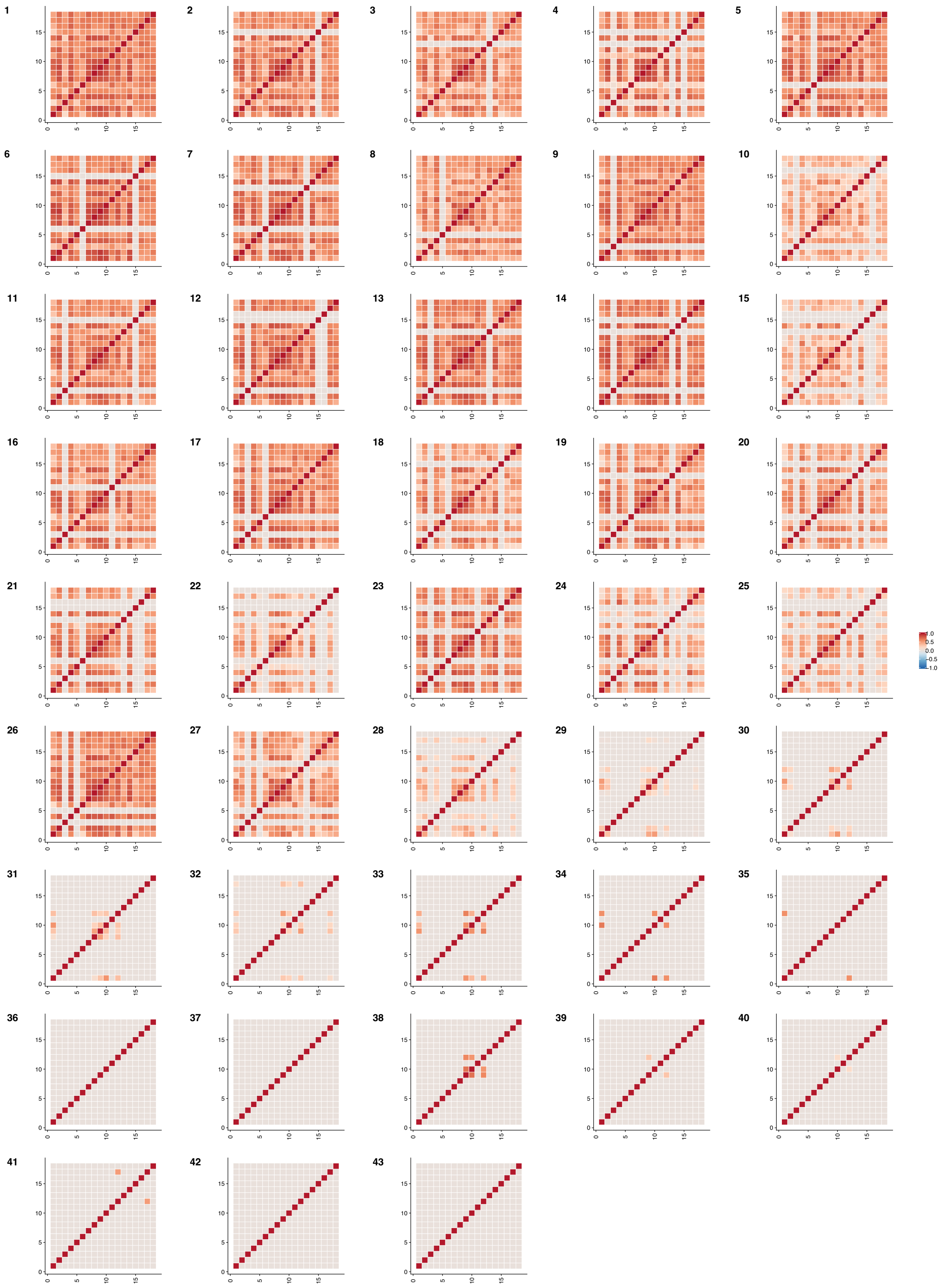

Figure S24: Correlations corresponding to the covariances used for each cluster in simulation 1.

| class num. \ dim. | 1 | 2 | 3 | 4 | 5 | 6 | 7 | 8 | 9 | 10 | 11 |
|---|---|---|---|---|---|---|---|---|---|---|---|
| 1 | 0 | 0 | 1.25 | 1.44 | 2.18 | 0 | 0 | 0 | 0 | 1.91 | 1.27 |
| 2 | 0 | 0 | 1.88 | 2.30 | 2.31 | 0 | 0 | 0 | 0 | 2.18 | 0 |
| 3 | 0 | 0 | 0.62 | 2.23 | 2.00 | 0 | 0 | 0 | 0 | 0 | 0 |
| 4 | 0 | 0 | 1.24 | 1.41 | 0 | 0 | 0 | 0 | 0 | 1.89 | 1.26 |
| 5 | 0 | 0 | 1.80 | 2.17 | 0 | 0 | 0 | 0 | 0 | 2.14 | 0 |
| 6 | 0 | 0 | 1.49 | 0.97 | 0 | 0 | 0 | 0 | 0 | 0 | 0 |
| 7 | 0 | 0 | 1.01 | 0 | 2.49 | 0 | 0 | 0 | 0 | 2.30 | 0.64 |
| 8 | 0 | 0 | 1.88 | -3.01 | 2.32 | 0 | 0 | 0 | 0 | 3.80 | 1.53 |
| 9 | 0 | 0 | 1.84 | -2.95 | 2.34 | 0 | 0 | 0 | 0 | 3.79 | 0 |
| 10 | 0 | 0 | 0.36 | -0.46 | 2.24 | 0 | 0 | 0 | 0 | 0 | 0 |
| 11 | 0 | 0 | 1.88 | -3.02 | 0 | 0 | 0 | 0 | 0 | 2.14 | 3.36 |
| 12 | 0 | 0 | 1.84 | -2.92 | 0 | 0 | 0 | 0 | 0 | 2.13 | 0 |
| 13 | 0 | 0 | 1.82 | -2.89 | 0 | 0 | 0 | 0 | 0 | 0 | 0 |
| 14 | 0 | 0 | 1.84 | -2.93 | 0 | 0 | 0 | 0 | 0 | 0 | -2.97 |
| 15 | 0 | 0 | 0 | 2.22 | 2.71 | 0 | 0 | 0 | 0 | 2.43 | 1.50 |
| 16 | 0 | 0 | 0 | 1.82 | 2.23 | 0 | 0 | 0 | 0 | 1.99 | 0 |
| 17 | 0 | 0 | 0 | 2.20 | 2.71 | 0 | 0 | 0 | 0 | 0 | 0 |
| 18 | 0 | 0 | 0 | 1.50 | 0 | 0 | 0 | 0 | 0 | 1.87 | 1.30 |
| 19 | 0 | 0 | 0 | 2.20 | 0 | 0 | 0 | 0 | 0 | 2.41 | 0 |
| 20 | 0 | 0 | 0 | 2.19 | 0 | 0 | 0 | 0 | 0 | 2.40 | -2.11 |
| 21 | 0 | 0 | 0 | 1.76 | 0 | 0 | 0 | 0 | 0 | 0 | 0 |
| 22 | 0 | 0 | 0 | 1.44 | 0 | 0 | 0 | 0 | 0 | 0 | -1.50 |
| 23 | 0 | 0 | 0 | 1.49 | 0 | 0 | 0 | 0 | 0 | -2.32 | 0 |
| 24 | 0 | 0 | 0 | 1.29 | 0 | 0 | 0 | 0 | 0 | -2.20 | -1.35 |
| 25 | 0 | 0 | 0 | -2.87 | 2.70 | 0 | 0 | 0 | 0 | 2.40 | 3.35 |
| 26 | 0 | 0 | 0 | -1.37 | 2.00 | 0 | 0 | 0 | 0 | 1.66 | 0 |
| 27 | 0 | 0 | 0 | -2.83 | 2.68 | 0 | 0 | 0 | 0 | 0 | 0 |
| 28 | 0 | 0 | 0 | -0.98 | 2.42 | 0 | 0 | 0 | 0 | -2.78 | 0 |
| 29 | 0 | 0 | 0 | -2.23 | 0 | 1.52 | 0 | 0 | 0 | 0 | 0 |
| 30 | 0 | 0 | 0 | -2.89 | 0 | 0 | 0 | 0 | 0 | 2.41 | 2.04 |
| 31 | 0 | 0 | 0 | -0.15 | 0 | 0 | 0 | 0 | 0 | 2.20 | 0 |
| 32 | 0 | 0 | 0 | -2.91 | 0 | 0 | 0 | 0 | 0 | 2.41 | -2.13 |
| 33 | 0 | 0 | 0 | -0.91 | 0 | 0 | 0 | 0 | 0 | 0 | 0 |
| 34 | 0 | 0 | 0 | -1.81 | 0 | 0 | 0 | 0 | 0 | 0 | -1.36 |
| 35 | 0 | 0 | 0 | -2.36 | 0 | 0 | 0 | 0 | 0 | -2.33 | 0 |
| 36 | 0 | 0 | 0 | -1.87 | 0 | 0 | 0 | 0 | 0 | -2.38 | -1.39 |
| 37 | 0 | 0 | -2.16 | -2.39 | 0 | 3.25 | 0 | 0 | 0 | 0 | -1.80 |
| 38 | 0 | 0 | -1.87 | -2.83 | 0 | 0 | 0 | 0 | 0 | 0 | 0 |
| 39 | 0 | 0 | -2.10 | -2.17 | 0 | 0 | 0 | 0 | 0 | 0 | -1.37 |
| 40 | 0 | 0 | -1.93 | -2.75 | 0 | 0 | 0 | 0 | 0 | -2.36 | -1.35 |

Table S8: Mean vectors used for each cluster in simulation 2.

| class num. \ dim. | 1 | 2 | 3 | 4 | 5 | 6 | 7 | 8 | 9 | 10 | 11 |
|---|---|---|---|---|---|---|---|---|---|---|---|
| 1 | 1 | 1 | 0.77 | 1.07 | 0.34 | 1 | 1 | 1 | 1 | 0.40 | 0.86 |
| 2 | 1 | 1 | 0.80 | 1.09 | 0.35 | 1 | 1 | 1 | 1 | 0.43 | 1 |
| 3 | 1 | 1 | 0.97 | 0.20 | 0.20 | 1 | 1 | 1 | 1 | 1 | 1 |
| 4 | 1 | 1 | 0.78 | 1.07 | 1 | 1 | 1 | 1 | 1 | 0.41 | 0.87 |
| 5 | 1 | 1 | 0.79 | 1.07 | 1 | 1 | 1 | 1 | 1 | 0.42 | 1 |
| 6 | 1 | 1 | 0.45 | 0.95 | 1 | 1 | 1 | 1 | 1 | 1 | 1 |
| 7 | 1 | 1 | 1.02 | 1 | 0.09 | 1 | 1 | 1 | 1 | 0.02 | 1.33 |
| 8 | 1 | 1 | 0.78 | 2.24 | 0.35 | 1 | 1 | 1 | 1 | 0.43 | 0.87 |
| 9 | 1 | 1 | 0.70 | 2.08 | 0.32 | 1 | 1 | 1 | 1 | 0.43 | 1 |
| 10 | 1 | 1 | 1 | 2.56 | 0.05 | 1 | 1 | 1 | 1 | 1 | 1 |
| 11 | 1 | 1 | 0.77 | 2.21 | 1 | 1 | 1 | 1 | 1 | 0.41 | 0.87 |
| 12 | 1 | 1 | 0.69 | 1.98 | 1 | 1 | 1 | 1 | 1 | 0.37 | 1 |
| 13 | 1 | 1 | 0.68 | 1.97 | 1 | 1 | 1 | 1 | 1 | 1 | 1 |
| 14 | 1 | 1 | 0.70 | 1.98 | 1 | 1 | 1 | 1 | 1 | 1 | 0.95 |
| 15 | 1 | 1 | 1 | 1.09 | 0.35 | 1 | 1 | 1 | 1 | 0.43 | 0.89 |
| 16 | 1 | 1 | 1 | 0.69 | 0.24 | 1 | 1 | 1 | 1 | 0.27 | 1 |
| 17 | 1 | 1 | 1 | 1.04 | 0.33 | 1 | 1 | 1 | 1 | 1 | 1 |
| 18 | 1 | 1 | 1 | 1.02 | 1 | 1 | 1 | 1 | 1 | 0.40 | 0.84 |
| 19 | 1 | 1 | 1 | 1.07 | 1 | 1 | 1 | 1 | 1 | 0.42 | 1 |
| 20 | 1 | 1 | 1 | 0.96 | 1 | 1 | 1 | 1 | 1 | 0.37 | 0.93 |
| 21 | 1 | 1 | 1 | 0.62 | 1 | 1 | 1 | 1 | 1 | 1 | 1 |
| 22 | 1 | 1 | 1 | 0.74 | 1 | 1 | 1 | 1 | 1 | 1 | 0.95 |
| 23 | 1 | 1 | 1 | 0.72 | 1 | 1 | 1 | 1 | 1 | 1.10 | 1 |
| 24 | 1 | 1 | 1 | 0.83 | 1 | 1 | 1 | 1 | 1 | 1.22 | 0.98 |
| 25 | 1 | 1 | 1 | 2.01 | 0.32 | 1 | 1 | 1 | 1 | 0.37 | 0.85 |
| 26 | 1 | 1 | 1 | 1.69 | 0.21 | 1 | 1 | 1 | 1 | 0.33 | 1 |
| 27 | 1 | 1 | 1 | 1.94 | 0.30 | 1 | 1 | 1 | 1 | 1 | 1 |
| 28 | 1 | 1 | 1 | 3.15 | 0.09 | 1 | 1 | 1 | 1 | 0.78 | 1 |
| 29 | 1 | 1 | 1 | 2.34 | 1 | 0.35 | 1 | 1 | 1 | 1 | 1 |
| 30 | 1 | 1 | 1 | 2.20 | 1 | 1 | 1 | 1 | 1 | 0.41 | 0.85 |
| 31 | 1 | 1 | 1 | 2.51 | 1 | 1 | 1 | 1 | 1 | 0.07 | 1 |
| 32 | 1 | 1 | 1 | 2.21 | 1 | 1 | 1 | 1 | 1 | 0.41 | 1.02 |
| 33 | 1 | 1 | 1 | 1.79 | 1 | 1 | 1 | 1 | 1 | 1 | 1 |
| 34 | 1 | 1 | 1 | 2.34 | 1 | 1 | 1 | 1 | 1 | 1 | 1.05 |
| 35 | 1 | 1 | 1 | 2.85 | 1 | 1 | 1 | 1 | 1 | 1.33 | 1 |
| 36 | 1 | 1 | 1 | 2.50 | 1 | 1 | 1 | 1 | 1 | 1.66 | 1.13 |
| 37 | 1 | 1 | 0.57 | 1.84 | 1 | 1.21 | 1 | 1 | 1 | 1 | 0.83 |
| 38 | 1 | 1 | 0.72 | 1.99 | 1 | 1 | 1 | 1 | 1 | 1 | 1 |
| 39 | 1 | 1 | 0.53 | 2.08 | 1 | 1 | 1 | 1 | 1 | 1 | 0.74 |
| 40 | 1 | 1 | 0.64 | 2.84 | 1 | 1 | 1 | 1 | 1 | 1.54 | 1.55 |

Table S9: Variance vectors used for each cluster in simulation 2.

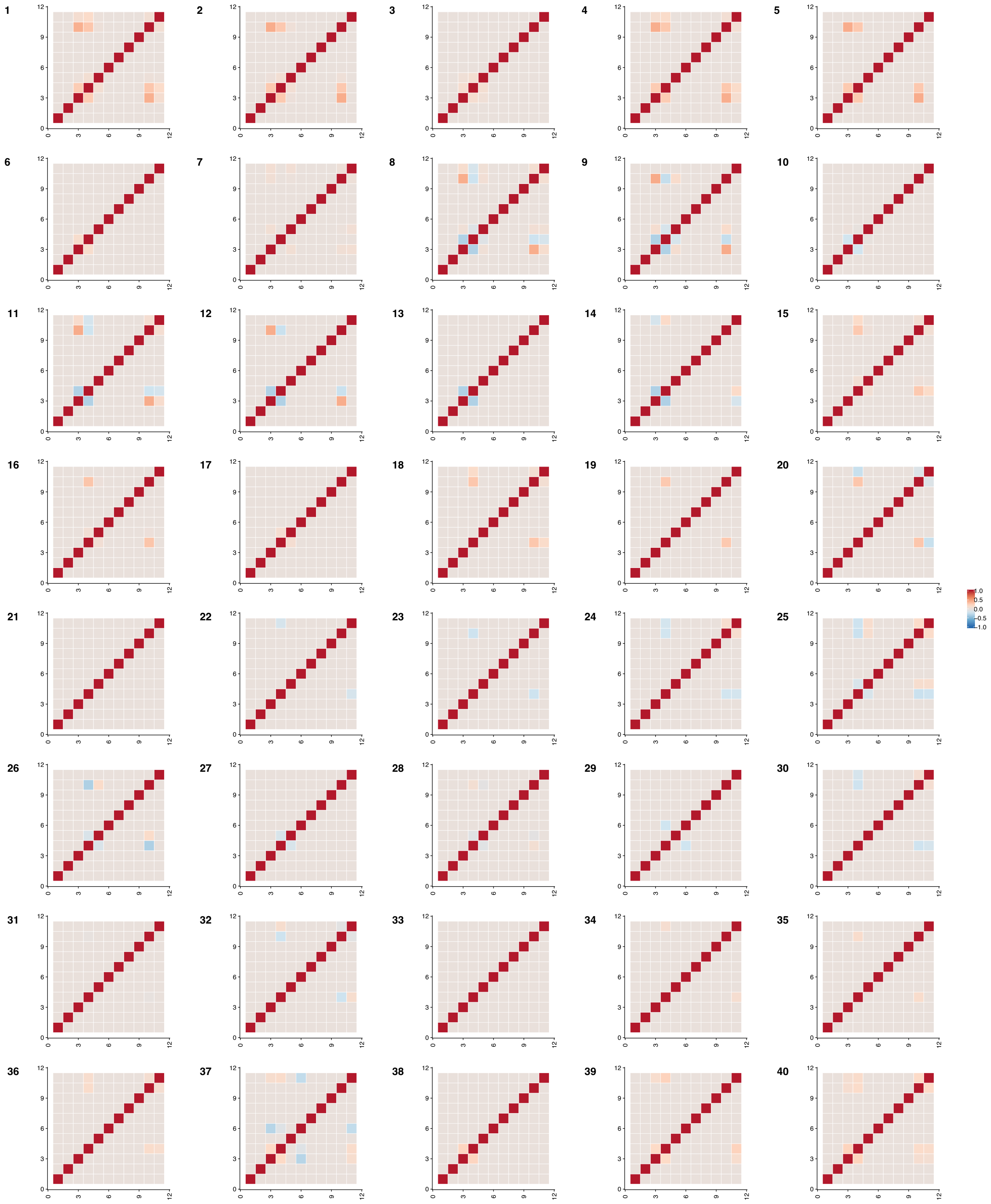


Figure S25: Correlations corresponding to the covariances used for each cluster in simulation 2.

| cluster num. | 1 | 2 | 3 | 4 | 5 | 6 | 7 | 8 | 9 | 10 | 11 |
|---|---|---|---|---|---|---|---|---|---|---|---|
| $\boldsymbol{\pi}$ | 4.15E-01 | 3.52E-04 | 5.22E-04 | 1.79E-03 | 2.39E-02 | 6.8E-06 | 1.47E-05 | 2.53E-05 | 2E-05 | 1.19E-05 | 2.33E-05 |

| cluster num. | 12 | 13 | 14 | 15 | 16 | 17 | 18 | 19 | 20 | 21 | 22 |
|---|---|---|---|---|---|---|---|---|---|---|---|
| $\boldsymbol{\pi}$ | 3.59E-05 | 3.55E-05 | 3.71E-04 | 4.24E-05 | 5.79E-05 | 4.91E-03 | 2.4E-06 | 2.5E-06 | 2.85E-05 | 6.6E-06 | 3.99E-04 |

| cluster num. | 23 | 24 | 25 | 26 | 27 | 28 | 29 | 30 | 31 | 32 | 33 | 34 |
|---|---|---|---|---|---|---|---|---|---|---|---|---|
| $\boldsymbol{\pi}$ | 1.68E-04 | 2.99E-04 | 1.51E-03 | 2.8E-06 | 4.4E-06 | 4.8E-06 | 1.60E-03 | 4.32E-02 | 3.84E-02 | 1.05E-02 | 9.66E-03 | 4.47E-01 |

Table S10: Mixing weights for each cluster used in simulation 3.

| dim. / class num. | 1 | 2 | 3 | 4 | 5 |
|---|---|---|---|---|---|
| 1 | 0.80 | 0.85 | 0.91 | 0.52 | 0.47 |
| 2 | 0 | 0 | 0 | 0 | 0 |
| 3 | 0 | 0 | 0 | 0 | -6.11 |
| 4 | 0 | 0 | 0 | -5.49 | 0 |
| 5 | 0 | 0 | 0 | -1.63 | -1.28 |
| 6 | 0 | 0 | -8.54 | 0 | 0 |
| 7 | 0 | 0 | -8.62 | 0 | -8.55 |
| 8 | 0 | 0 | -8.54 | -8.52 | 0 |
| 9 | 0 | 0 | -8.58 | -8.57 | -7.38 |
| 10 | 0 | -8.28 | 0 | 0 | 0 |
| 11 | 0 | -8.38 | 0 | 0 | -7.17 |
| 12 | 0 | -8.35 | 0 | -7.28 | 0 |
| 13 | 0 | -8.26 | 0 | -7.32 | -8.40 |
| 14 | 0 | -6.99 | -6.83 | 0 | 0 |
| 15 | 0 | -8.20 | -7.66 | 0 | -7.00 |
| 16 | 0 | -8.14 | -7.61 | -7.15 | 0 |
| 17 | 0 | -4.92 | -5.39 | -5.44 | -5.06 |
| 18 | -5.94 | 0 | 0 | 0 | 0 |
| 19 | -5.83 | 0 | 0 | 0 | -8.02 |
| 20 | -5.80 | 0 | 0 | -8.02 | 0 |
| 21 | -5.91 | 0 | 0 | -8.10 | -8.20 |
| 22 | -5.32 | 0 | -6.58 | 0 | 0 |
| 23 | -5.74 | 0 | -7.73 | 0 | -6.22 |
| 24 | -5.33 | 0 | -6.77 | -5.81 | 0 |
| 25 | -5.34 | 0 | -6.02 | -5.63 | -5.21 |
| 26 | -5.90 | -6.16 | 0 | 0 | 0 |
| 27 | -5.93 | -6.17 | 0 | 0 | -7.97 |
| 28 | -5.96 | -6.14 | 0 | -7.97 | 0 |
| 29 | -4.99 | -5.21 | 0 | -5.72 | -5.15 |
| 30 | -2.71 | -3.05 | -3.87 | 1.22 | 1.63 |
| 31 | -2.24 | -2.34 | -2.67 | 0 | 0 |
| 32 | -5.39 | -5.88 | -5.96 | 0 | -5.01 |
| 33 | -5.66 | -6.67 | -6.65 | -5.45 | 0 |
| 34 | -6.95 | -6.94 | -7.00 | -7.42 | -7.25 |

Table S11: Mean vectors used for each cluster in simulation 3.

| class num. \ dim. | 1 | 2 | 3 | 4 | 5 |
|---|---|---|---|---|---|
| 1 | 5.54 | 5.45 | 5.35 | 6.01 | 5.70 |
| 2 | 1 | 1 | 1 | 1 | 1.00 |
| 3 | 1 | 1 | 1 | 1 | 17.92 |
| 4 | 1 | 1 | 1 | 16.76 | 1.00 |
| 5 | 1 | 1 | 1 | 11.75 | 13.46 |
| 6 | 1 | 1 | 18.27 | 1 | 1.00 |
| 7 | 1 | 1 | 19.02 | 1 | 19.59 |
| 8 | 1 | 1 | 18.81 | 19.91 | 1.00 |
| 9 | 1 | 1 | 18.98 | 20.30 | 19.70 |
| 10 | 1 | 16.90 | 1 | 1 | 1.00 |
| 11 | 1 | 17.19 | 1 | 1 | 19.40 |
| 12 | 1 | 17.07 | 1 | 20.11 | 1.00 |
| 13 | 1 | 17.52 | 1 | 20.39 | 19.82 |
| 14 | 1 | 16.57 | 18.06 | 1 | 1.00 |
| 15 | 1 | 17.79 | 18.95 | 1 | 19.62 |
| 16 | 1 | 17.47 | 19.20 | 20.22 | 1.00 |
| 17 | 1 | 15.39 | 14.77 | 16.49 | 15.92 |
| 18 | 18.96 | 1 | 1 | 1 | 1.00 |
| 19 | 19.56 | 1 | 1 | 1 | 19.29 |
| 20 | 19.17 | 1 | 1 | 19.89 | 1.00 |
| 21 | 20.01 | 1 | 1 | 20.41 | 19.63 |
| 22 | 18.82 | 1 | 17.49 | 1 | 1.00 |
| 23 | 19.24 | 1 | 19.19 | 1 | 19.14 |
| 24 | 19.71 | 1 | 18.54 | 19.70 | 1.00 |
| 25 | 18.49 | 1 | 16.94 | 18.14 | 18.68 |
| 26 | 19.39 | 17.18 | 1 | 1 | 1.00 |
| 27 | 20.13 | 17.71 | 1 | 1 | 20.05 |
| 28 | 19.98 | 17.50 | 1 | 20.44 | 1.00 |
| 29 | 18.73 | 16.75 | 1 | 18.16 | 18.16 |
| 30 | 20.04 | 18.37 | 21.15 | 13.46 | 13.79 |
| 31 | 10.98 | 9.99 | 8.33 | 1 | 1.00 |
| 32 | 12.26 | 11.72 | 12.25 | 1 | 14.92 |
| 33 | 13.38 | 10.10 | 11.43 | 14.80 | 1.00 |
| 34 | 6.28 | 6.24 | 6.91 | 6.52 | 5.89 |

Table S12: Variance vectors used for each cluster in simulation 3.

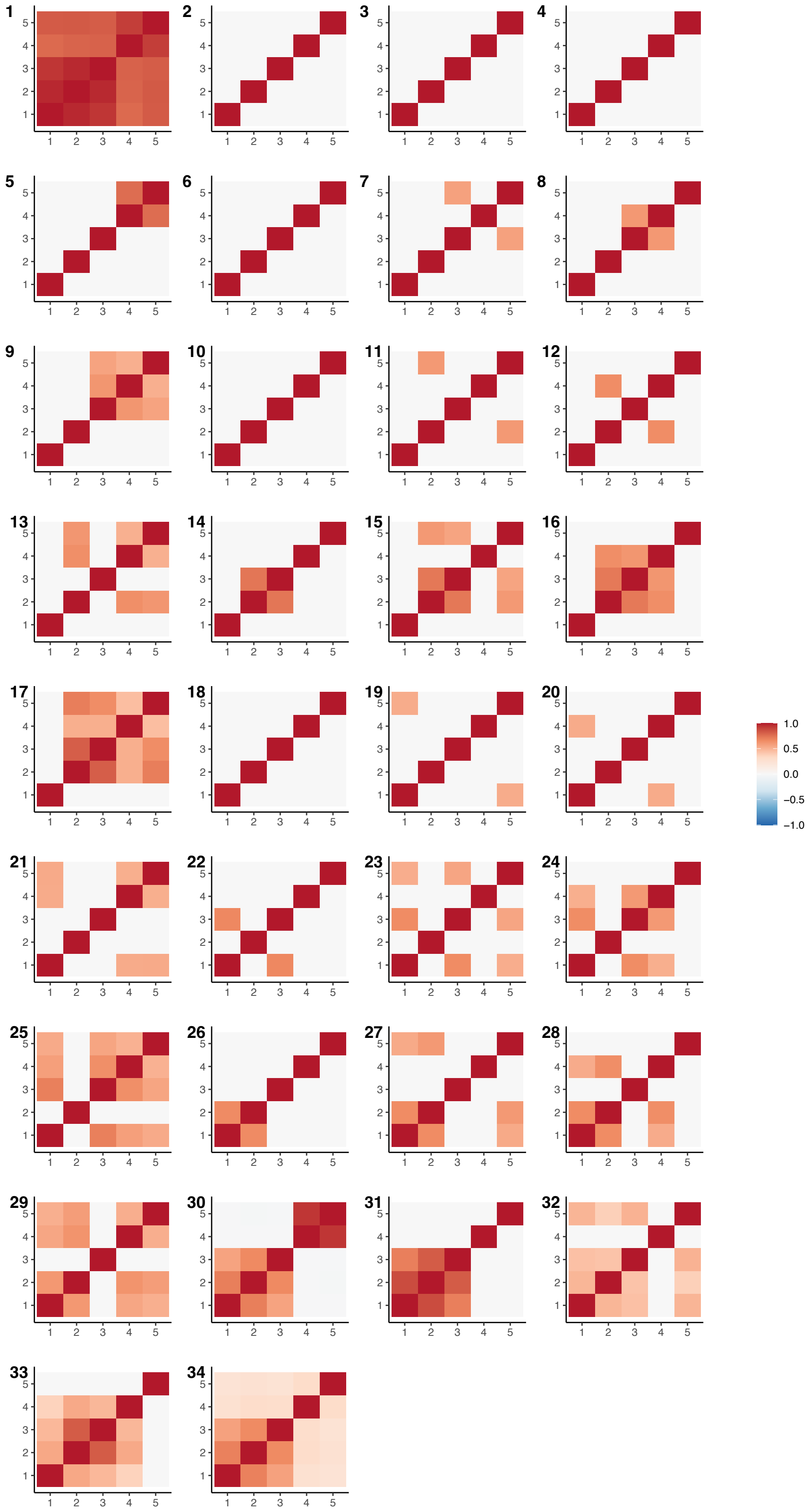

Figure S26: Correlations corresponding to the covariances used for each cluster in simulation 3.

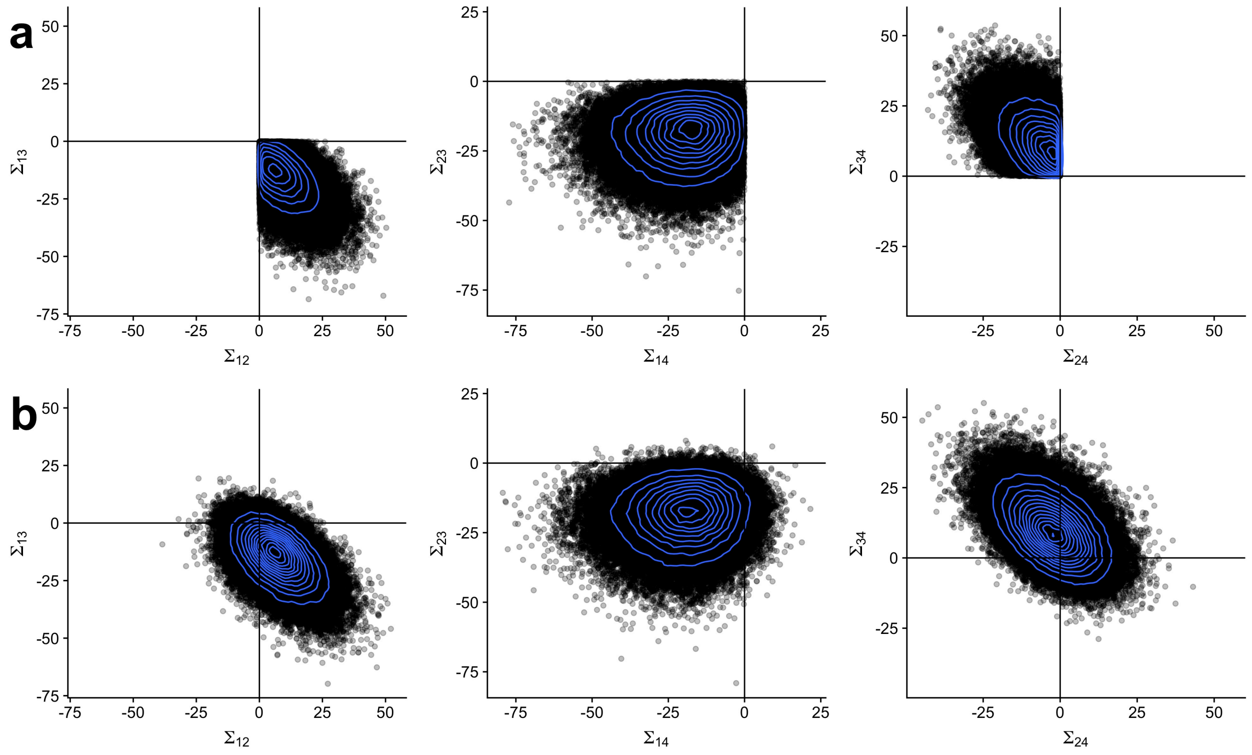


Figure S27: Constrained covariance draws plotted in pairs of off-diagonal elements. **a,** Constrained samples generated with Algorithm 1 and **b,** the corresponding ordinary Wishart samples. Comparing within columns, one observes that data generated with Algorithm 1 closely match their unconstrained counterparts, indicating they serve as an appropriate proposal distribution.

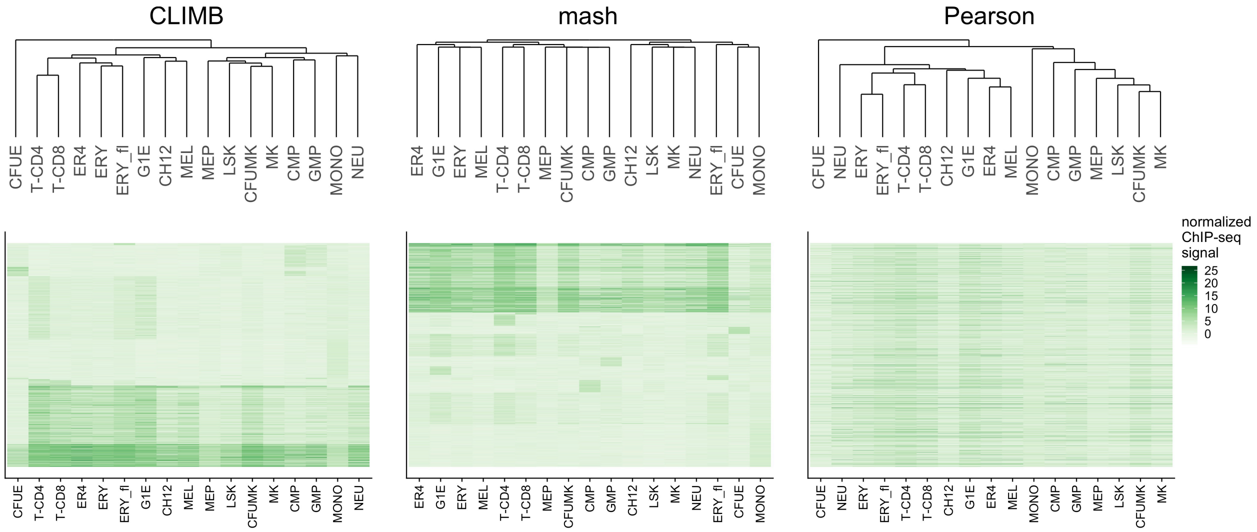


Figure S28: Heatmaps displaying bi-clusterings of chromosome 7 CTCF ChIP-seq data based on CLIMB, mash, and Pearson correlation. The columns, corresponding to different cell populations, are ordered according to the dendrogram for each clustering method. The rows, corresponding to each binding site, are ordered based on class membership (for CLIMB and mash) and Pearson correlation for the other.

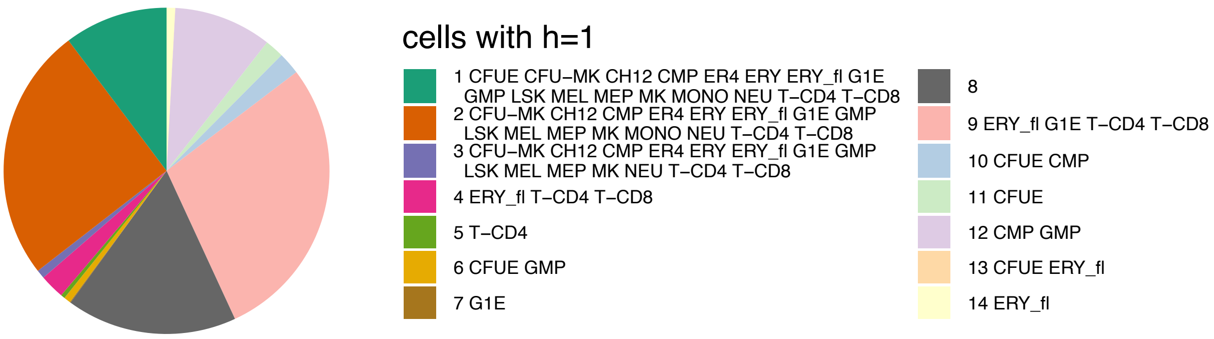


Figure S29: Class sizes in the CTCF ChIP-seq data analysis of chromosome 7. The final model estimated 14 non-empty classes, each described by association vectors with binary elements. The legend shows which cell types in each class were assigned a 1, indicating presence of CTCF.

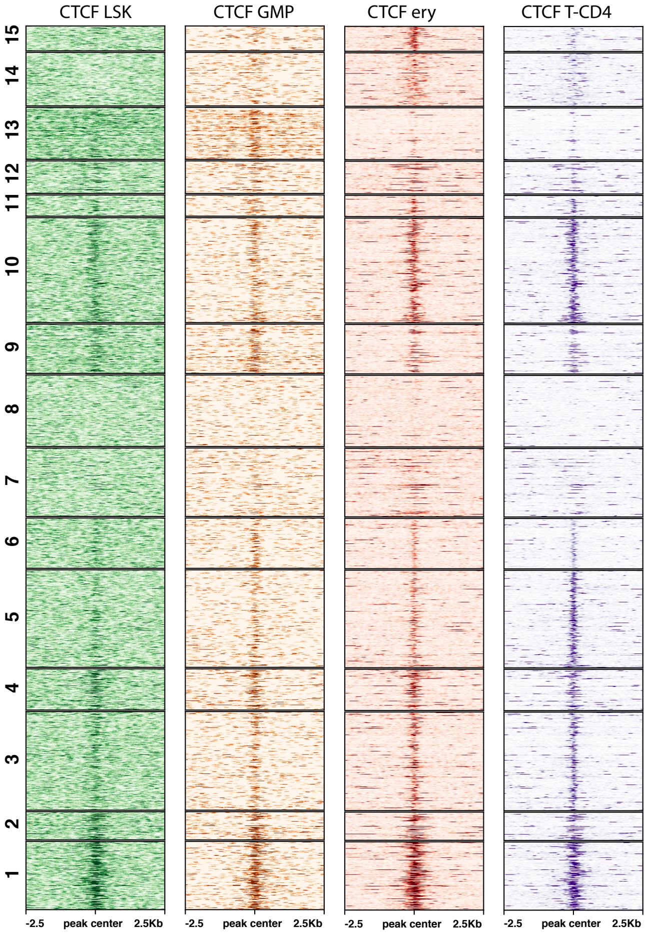


Figure S30: CTCF ChIP-seq signals for binding sites on chromosome 11 in LSK, GMP, ERY, and T-CD4 cells. Same as in Fig. S13, but hierarchically clustered using Pearson correlation and cutting the tree to have 15 clusters.

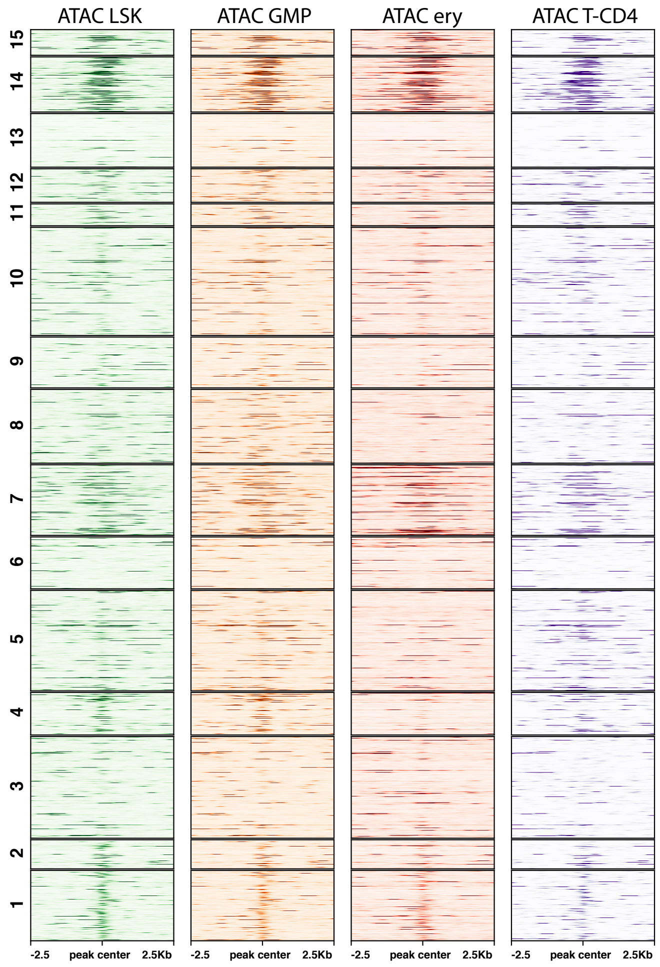


Figure S31: ATAC-seq signals for CTCF binding sites on chromosome 11 in LSK, GMP, ERY, and T-CD4 cells. Same as in Fig. S14, but hierarchically clustered using Pearson correlation and cutting the tree to have 15 clusters.

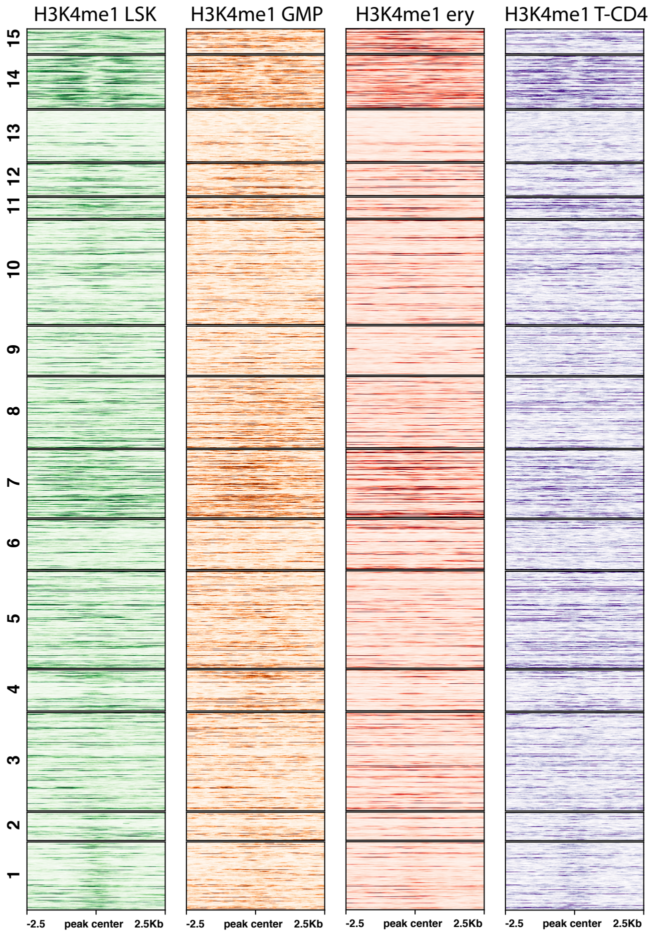


Figure S32: H3K4me1 ChIP-seq signals for CTCF binding sites on chromosome 11 in LSK, GMP, ERY, and T-CD4 cells. Same as in Fig. S15, but hierarchically clustered using Pearson correlation and cutting the tree to have 15 clusters.

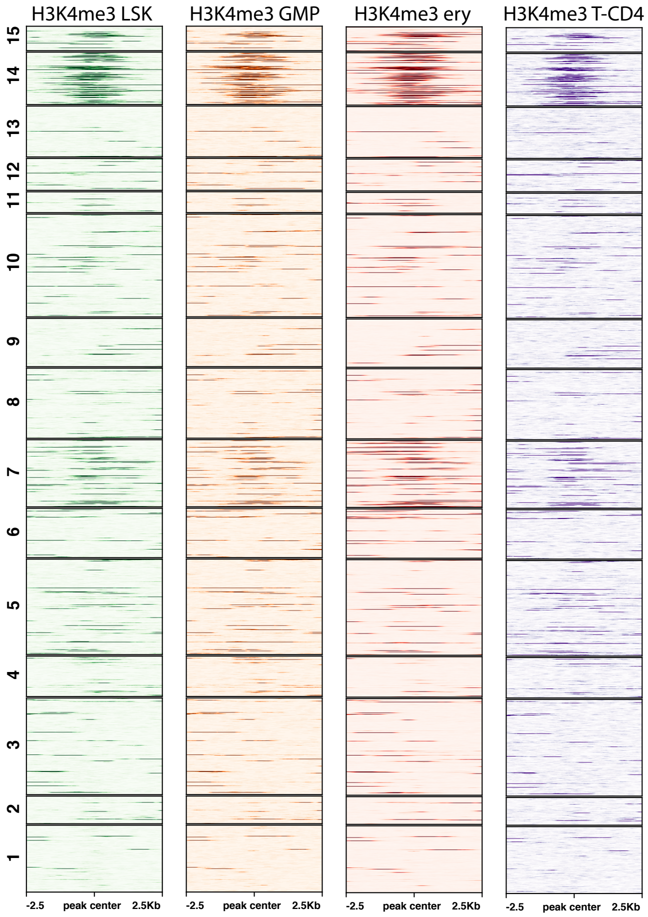


Figure S33: H3K4me3 ChIP-seq signals for CTCF binding sites on chromosome 11 in LSK, GMP, ERY, and T-CD4 cells. Same as in Fig. S16, but hierarchically clustered using Pearson correlation and cutting the tree to have 15 clusters.
